# Deciphering Molecular Language for Spectral Interpretation with a Foundation Model Pretrained on Repository-Wide Metabolomics Data

**DOI:** 10.64898/2026.11.14.766667

**Authors:** Yifan Li, Yunhua Zhong, Pan Liu, Zikun Wang, Jiaqing Xie, Hongxin Xiang, Zihan Chen, Zhiyuan Yuan, Yuqiang Li, Jun Xia

## Abstract

In-depth, scalable structural interpretation of tandem mass spectra from diverse biological tissues and environmental samples is essential for understanding the molecular diversity of cellular environments and the metabolic architecture of living systems. Although community-driven resources such as GNPS and MassIVE have aggregated massive volumes of tandem mass spectra spanning a wide range of organisms, clinical cohorts and environmental surveys, modeling and interpreting this exponentially expanding spectral corpus to distill molecular information and derive meaningful biological insights poses a fundamental challenge for established computational methods. Here we present UltraMS, a dual-polarity foundation model tailored to decipher the molecular language latent in tandem mass spectra through large-scale pretraining on more than 160 million spectra encompassing both positive and negative ion modes. Our findings illustrate that UltraMS is capable of effectively decoding the chemical semantics of molecules underlying mass spectra via self-supervised learning, achieving superior performance across a variety of downstream tasks, including elemental detection, molecular identification, molecular property profiling, and metabolic landscape characterization. Remarkably, UltraMS enables conducting molecular identification with fine-grained spectral interpretability, simultaneously elucidating molecular structures from untargeted MS/MS spectra and providing peak-by-peak substructural attributions via annotation transfer. Deployed on a clinical cohort of human plaque samples, UltraMS demonstrates significant potential in discovering previously unrecognized phenotype-linked metabolites supported by structurally grounded fragment-ion formation pathways, while effectively correcting likely misannotated clinical markers. Furthermore, building on the structure-aware spectral representations learned by UltraMS, we construct Ultra-Atlas, a chemical-property-resolved map that organizes millions of experimental spectra into a unified coordinate system, enabling millisecond-scale molecular profiling and offering practical guidance for exploratory navigation across vast chemical space. Applied to human scalp metabolomics, UltraAtlas effectively captures distinct metabolic variations across individuals, highlighting phenotype-specific chemical signatures while shedding light on the chemical niches of the resident microbiome. Together, UltraMS and UltraAtlas constitute a generalizable framework for structural elucidation and biological discovery from tandem mass spectra, empowering the comprehensive exploration of the largely undeciphered metabolomes across different domains of life.

## 1 Introduction

Liquid chromatography tandem mass spectrometry (LC-MS/MS) represents the prevailing technology for metabolism research and has become a central platform for high-throughput molecular profiling. It is highly sensitive, broadly applicable, and capable of detecting thousands of molecular features from biochemical or environmental specimens via high-throughput scanning [1, 2]. In the standard LC-MS/MS workflow, molecules within a sample are separated by chromatography and then dissociated to generate tandem mass spectrometry(MS), whose fragmentation patterns serve as the primary evidence for structural inference [3]—a process that demands substantial expert knowledge to guide reasoning over the latent correspondence between observed peaks and molecular substructures. Public repositories such as MassIVE host tremendous collections of experimental spectra [4, 5], offering unprecedented opportunities to learn chemically meaningful and transferable spectral representations directly from raw, unannotated spectra in a self-supervised manner. However, unlike protein sequences represented by alphabets of fewer than 30 symbols or words in Natural Language Processing (NLP) drawn from a well-defined vocabulary, each peak in mass spectrometry initially carries only numerical attributes, with its chemical meaning undetermined and varying substantially across diverse spectral contexts. This presents a major bottleneck to effectively tokenizing and embedding mass spectra for large-scale pretraining modern deep learning architectures. In addition, mass spectra are intrinsically sparse, instrument-dependent, and partially informative: the same molecule can produce significantly different signals across collision energies, instruments, chromatographic settings, and ionization modes, making accurate molecular identification from raw mass spectral features challenging and leaving a large proportion of spectra uninterpreted [2, 6]. This unique combination of large-scale data availability despite pervasive spectral noise across heterogeneous instruments and the profound potential to illuminate the dark molecule space makes MS data an especially compelling and attractive opportunity for learning implicit chemical knowledge directly from spectra.

To harness mass spectrometry for diverse downstream analyses, various deep learning methods have been developed with domain-specific architectures [7, 8]. For instance, CANOPUS utilizes a deep neural network alongside molecular formulas and a 4,262-bit structural fingerprint derived from the authoritative SIRIUS toolkit to directly predict over 2,000 compound classes for unknown metabolites, allowing researchers to gain valuable biological insights without relying on exhaustive peak-by-peak annotation [3, 9]. Similarly, MIST leverages spectral peak formula annotations deduced by SIRIUS as chemical prior enhanced inputs to a domain-specific “formula-transformer” to predict molecular substructures reflected by fingerprints, demonstrating its superior ability to detect essential differences in metabolic profiles between diseased cohorts (e.g., IBD) and healthy individuals [10]. Most recently, DeepMet highlighted the capacity of chemical language models to break through the boundaries of the known metabolome; by learning the latent structural patterns of metabolic space, it demonstrated the unprecedented potential to discover dozens of previously unrecognized mammalian metabolites from untargeted metabolomics data [11]. Furthermore, DiffMS integrates a fingerprint-to-molecule decoder pre-trained on 2.8 million fingerprint–structure pairs with a transformer-based spectral encoder to enable de novo molecule identification independent of existing spectral libraries, stepping beyond the limitation imposed on molecule identification by the scarcity of annotated molecular structures in existing mass spectrometry databases [12].

However, these advanced approaches are inherently bound by the small fraction of spectra with verified structural annotations for task-specific supervision. While raw experimental biological data is increasingly abundant across wide-ranging scientific disciplines, reliably annotated spectrum–structure pairs remain prohibitively expensive and constrain the development of robust analytical approaches for untargeted metabolomics. Large-scale self-supervised pre-training has emerged as a transformative solution to this challenge. By learning directly from vast repositories of unannotated data, these models capture underlying principles and extract essential domain insights without labor-intensive human labeling. In the life sciences, spanning protein, gene sequencing, transcriptomics, and single-cell omics—foundation models trained on millions of high-throughput biological sequences and cellular profiles demonstrate remarkable capability in deciphering the “language of life.” [13–16] These foundational models enable highly effective transfer to diverse downstream tasks with effective fine-tuning. More recently, DreaMS introduced a foundation model paradigm for mass spectrometry by incorporating essential spectral domain knowledge as an inductive bias within a Graphormer framework [17]. Through masked peak pre-training on 24 million high-quality spectra, DreaMS constructs a robust embedding space for molecules based on their mass spectra, serving as an adaptable foundation that powers molecular search, element identification, and molecular characterization [18]. However, diverse limitations remain. Notably, DreaMS exhibits a pronounced sensitivity to data quality: incorporating larger, uncurated datasets of variable-fidelity instruments paradoxically degrades model performance, leading to its pre-training restricted in the curated GeMS-A10 subset (24M spectra), despite the availability of broader tiers (such as Level C GeMS) exceeding 100M spectra [18]. Furthermore, its training is strictly limited to positive ion mode data, which creates an intrinsic barrier to downstream scenarios dominated by negative ionization mode, such as the analysis of dissolved organic matter widely present in environmental samples [19]. What’s more, although DreaMS offers an informative global mass spectral embedding through its precursor token, it exhibits limited ability to learn spectral-peak-level embeddings that hold the potential to comprehensively interpret a spectrum and reveal the molecular rationale underlying identification results, posing a fundamental challenge to leverage annotations transferred through spectral peak embeddings and integrating evidence from individual peaks into plausible structural explanations, leaving reasoning across the entire spectrum out of reach for a mass spectrum foundation model.

In this article, we addressed these challenges and developed UltraMS, a large-scale foundation model for tandem mass spectrometry with approximately 200 million parameters, applicable to a wide range of downstream molecular discovery tasks across diverse biochemical domains. We assembled UltraMSdata, an unprecedented pretraining corpus comprising 161 million curated spectra that, for the first time, spans both positive- and negative-ionization modes across diverse metabolic and biological scenarios. We designed a Continuous Multi-resolution Peak Encoding (CMPE) scheme, enabling UltraMS to not only efficiently model spectral peaks without limiting its representation capacity by a fixed-size vocabulary, but also capture the rich chemical semantics implicit in fragmentation patterns while preserving high-precision *m/z* and intensity fidelity. We developed a Sequentially Conditioned Fine-grained Reconstruction (SCFR) decoder that progressively recovers masked spectral peaks from a coarse resolution of 10 Da down to 0.1 Da, sequentially leveraging coarse-level chemical priors learned by CMPE to refine more fine-grained spectral peak estimates during large-scale pretraining, empowering UltraMS to not only reveal core molecular scaffolds by locating the fragment’s broad mass window at the 10-Da scale, but also actively capture subtle mass defects and exact elemental compositions down to a resolution of 0.1 Da by exploiting semantic embeddings learned by CMPE as prior guidance. We show that these novel algorithms tailored for large-scale pretraining substantially improve the modeling capacity for mass spectra, unlocking the opportunity to learn the implicit and valuable chemical knowledge directly from massive amounts of unlabeled raw metabolomics data.

Overall, UltraMS demonstrates the capability of mass-spectral foundation models to decode the latent chemical language of tandem mass spectrometry across three primary aspects. First, pretrained at an unprecedented scale across comprehensive ionization modes spectral repositories, UltraMS establishes a robust foundation for mass-spectral representation learning that benefits a diverse range of downstream works. By delivering state-of-the-art performance across elemental determination, molecular property profiling, spectrum-to-structure annotation, and library matching on diverse benchmarks, our results verify the proposing self-supervised pretraining paradigm as a generalizable engine for computational metabolomics. Second, UltraMS offers superior interpretability through chemically grounded peak- and spectrum-level representations, which both captures the overall molecular identity and assists in retrospectively tracing fragmentation events. By explicitly constructing correspondences between individual spectral peaks and molecular fragment substructures, UltraMS establishes a cornerstone for peak-level structural reasoning and enables transparent rationalization of spectral identification decisions. In a real-world application to human carotid plaque metabolomics, UltraMS not only resolves ambiguous annotations and rectifies misassignments where the original study failed to distinguish subtle structural variations, but also uncovers previously uncharacterized metabolic signatures pointing to potential immunometabolic dysregulation and pro-inflammatory pathways underlying human carotid plaque vulnerability, demonstrating its emergent capacity to derive biologically actionable hypotheses from complex clinical specimens. Third, by leveraging UltraMS’s chemically structured latent space, we construct UltraAtlas, a navigable reference atlas that establishes a unified coordinate system across the global mass-spectrometric landscape, turning millions of collective experimental records of tandem mass spectrometry into an actionable map resolved into characteristic chemical territories for guided chemical exploration. Coupled with an efficient search architecture achieving millisecond-level latency, UltraAtlas enables seamless, confidence-calibrated exploration that not only captures sample-level metabolic phenotypes across diverse cohorts, but also prioritizes individual discriminative molecular features. Demonstrating this utility in an untargeted human scalp metabolomics cohort, UltraAtlas substantially improves chemical-class coverage beyond conventional tools and illuminates valuable metabolic shifts that correlate with host physiological states and resident microbiome dynamics. Based on these findings, we envision that UltraMS and UltraAtlas will greatly deepen our understanding of the chemical language latent in the subtle patterns of tandem mass spectra and help the scientific community accelerate the translation of the growing wealth of mass-spectrometry data into discoveries of molecular diversity and its biological significance.

## 2 Results

### 2.1 Overview of UltraMS and UltraMSdata

We introduced UltraMS, a foundation model with approximately 200 million parameters that learns the chemical language of mass spectra. UltraMS was pretrained on over 160 million tandem mass spectra using a masked-and-recovery pretraining approach. We began by assembling a diverse, multi-domain corpus from publicly available GNPS–MassIVE repositories [4, 5]. To ensure spectral quality, we developed a quality-control algorithm that incorporated centroid evaluation for spectra with uncertain acquisition metadata and instrument-accuracy filtering, yielding 161 million curated spectra that constitute UltraMSdata (Fig. 1a). UltraMSdata is the first pretraining corpus tailored for foundation-model pretraining in mass spectrometry that covers both positive- and negative-ion modes. It encompasses diverse general-purpose applications, including human and animal samples, natural products, and environmental, clinical, plant and food domains. Among the multiple GeMS splits curated alongside the pioneering mass-spectrometry foundation model DreaMS, only GeMS-A10 (24 million spectra) served as its pretraining set while further scaling beyond this subset degraded model performance. UltraMSdata substantially surpasses this scale as a pretraining corpus, comprising 161 million spectra in total—approximately sevenfold more than the previous GeMS-A10, including an additional 11 million negative-mode spectra absent from prior large-scale pretraining corpora. This broadened categorical scope and comprehensive ion-mode coverage empower UltraMS to capture the underlying chemical language across diverse mass-spectrometry regimes and achieve superior performance across a wide range of downstream tasks.

**Figure 1.**
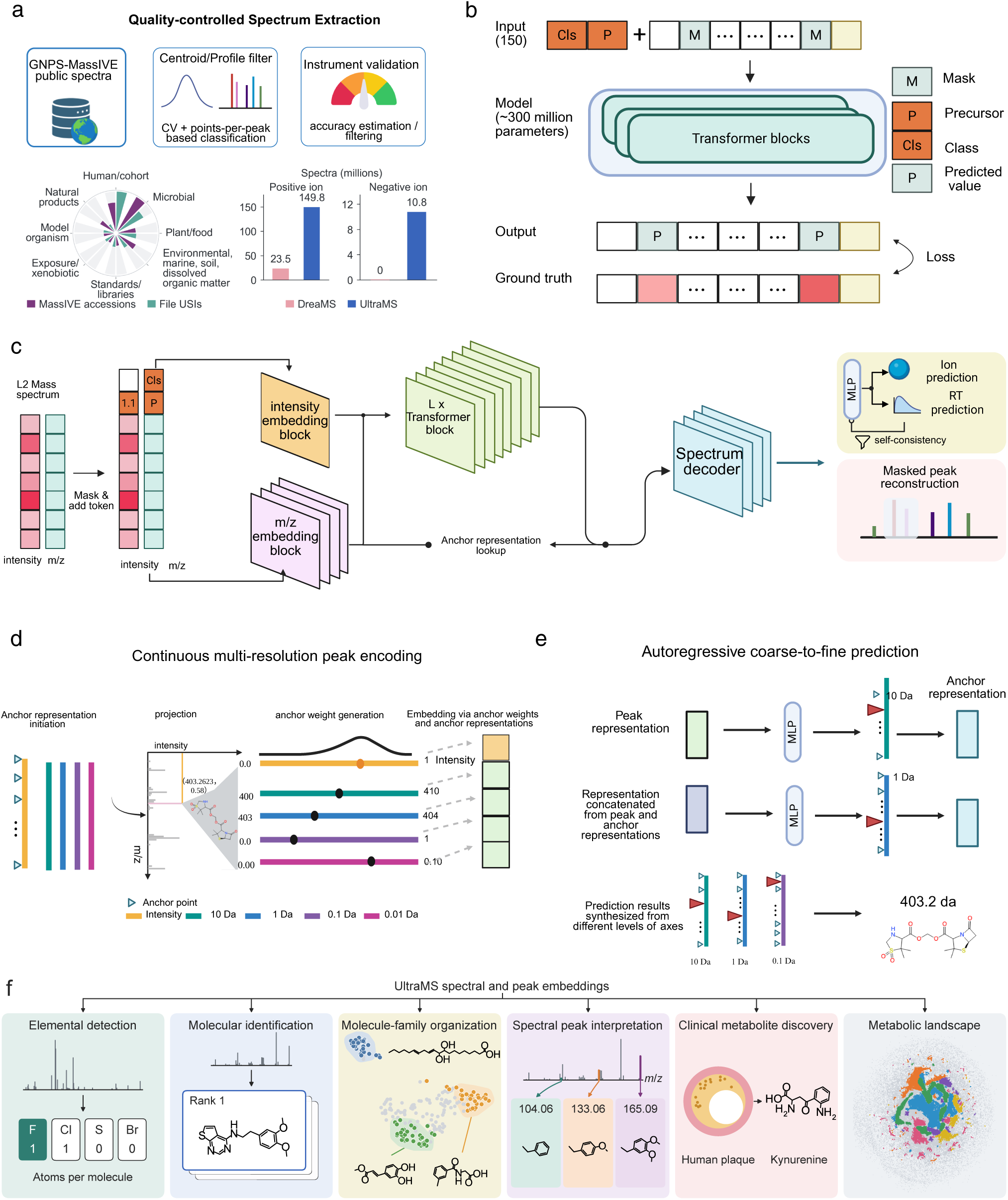
Overview of UltraMSdata and the UltraMS framework. a, Construction of UltraMSdata from public GNPS–MassIVE spectra through quality control. The radial plot highlights the broad spectrum of experimental contexts and data sources underlying UltraMSdata. Bar plots compare positive- and negative-ion spectrum counts with the pretraining corpus of the previous mass-spectrometry foundation model DreaMS. **b,** Masked spectral-peak reconstruction. Selected spectral peaks are masked, and UltraMS reconstructs their *m/z* and intensity by integrating information from the precursor ion and other spectral peaks through self-attention propagation. CLS denotes the token used to obtain the spectrum-level representation. **c,** UltraMS architecture and training tasks. Embeddings of spectral-peak *m/z* and intensity are processed by the transformer to generate spectrum-level and spectral peak-level representations. Masked spectral-peak reconstruction is followed by ion-mode discrimination and retention-time estimation, with self-consistency filtering to exclude unreliable retention-time labels. **d,** Continuous Multi-resolution Peak Encoding (CMPE). Continuous *m/z* and intensity values are represented by weighted combinations of the learnable embeddings of spectral representation anchors. Weights depend on the distance between each *m/z* value and the spectral representation anchor positions, with nearer anchors receiving larger weights. The *m/z* representation combines four resolutions, from 10 to 0.01 Da. **e,** Sequentially Conditioned Fine-grained Reconstruction (SCFR). The decoder first estimates *m/z* from probabilities over 10-Da intervals. This estimate is enhanced by leveraging learnable embedding from CMPE and concatenated with the spectral peak-level representation to predict a finer position within the corresponding interval. This process gradually proceeds through 1- and 0.1-Da resolutions. Ultimately, the predicted positions at all three levels are combined to reconstruct the spectral-peak *m/z*. **f**, Broad applicability of UltraMS across diverse downstream tasks. Learned spectrum- and peak-level representations of UltraMS enable elemental detection, spectral interpretation, untargeted molecular identification, spectrum-based molecular family organization, clinical metabolite discovery, and metabolic landscape characterization.

The core model of UltraMS consists of stacked transformer layers with multi-head attention [20] that simultaneously generate spectral- and peak-level embeddings (Fig. 1b). Training proceeds in two stages: first on UltraMSdata using a masked-and-recovering approach that disrupts and recovers individual spectral peaks, followed by two auxiliary tasks for ion-mode prediction and retention-time regression (Fig. 1c). In the pretraining stage, we introduce Continuous Multi-resolution Peak Encoding (CMPE) (Fig. 1d) as the embedding module that projects mass spectra into representations that are fed into the subsequent transformer. This technique addresses the continuous nature of ion peaks, whose *m/z* values span a near-infinite numerical range and cannot be simply represented by mapping them through a discrete word-lookup table as in NLP. Instead, CMPE projects each spectral peak, consisting of an *m/z*–intensity pair, into an embedding space through multiple resolutions of anchor points uniformly distributed along different *m/z* axes (See details in Methods 4). For spectrum decoding from the hidden states, we propose Sequentially Conditioned Fine-grained Reconstruction (SCFR) (Fig. 1e) to gradually reveal the true *m/z* value from a coarse range to a fine-grained region at the *m/z* axis. SCFR transforms *m/z* reconstruction into a soft, multi-stage process in which the model first predicts a coarse peak location and then leverages coarse-level information from the CMPE embeddings to sequentially identify finer-grained positions until the finest resolution is reached. We show that this design substantially improves *m/z* reconstruction performance compared with the bin-based prediction used in prior models. For the two auxiliary tasks, we design a self-consistency algorithm for retention-time prediction that filters out unreliable RT labels, which typically arise from systematic errors and varying experimental conditions, enabling the model to efficiently leverage the chemically informative RT signal during training. For ion-mode prediction, we employ an MLP prediction head that forces the model to attend to the distinct chemical fragmentation characteristics of different ionization modes. After pretraining, the pretrained model can adapt to new datasets and various downstream tasks. We demonstrate UltraMS’s ability to generate meaningful representations across a variety of essential tasks in mass-spectrometry research, including mass-spectrum annotation, molecular-identity search, clinical-phenotype related metabolite detection and global metabolic-landscape construction (Fig. 1f).

### 2.2 Chemical knowledge emerges hierarchically from self-supervised pretraining on large corpus

In our study, we found that finer-grained chemical understanding progressively emerges from global to spectral peak-level embeddings under the self-supervised learning framework (Supplementary Fig. 1). We first elucidated this foundational advantage by evaluating UltraMS against the pioneering pretrained baseline DreaMS [18] on the core self-supervised task of corrupted peak reconstruction. UltraMS demonstrated superior capability in capturing precise *m/z* values (Fig. 2a). Specifically, UltraMS achieved 37.7% accuracy compared to 3.5% for DreaMS on MassSpecGym, and even more than doubled the accuracy on DreaMS’s own training domain GeMS-A10 (55.2% versus 23.8%). Additionally, we observed that UltraMS exhibits robust representations against diverse algorithmically simulated noise perturbations (Supplementary Fig. 2). We next examined whether the model effectively captures the underlying ionization mode directly from its global representations. UltraMS improved test classification accuracy from 88.8% to 94.9% compared with DreaMS, lifting both [M − H]^−^ and [M + H]^+^ F1 scores by approximately 6 percentage points (Fig. 2b). This demonstrates a deepened capacity to model the physical generation process across distinct ionization regimes. Furthermore, we investigated whether incorporating negative-mode data during pretraining provides concrete representation benefits for negative-mode spectra. We conducted validation experiments on the Spectraverse dataset [21] following the same contrastive learning protocol on MoNA reported by DreaMS (see detailed method in Supplementary Material Sections S1.7 and S1.8). We visualized the resulting embeddings alongside DreaMS and raw spectral baselines, coloring data points according to molecular cluster (Fig. 2c). Notably, UltraMS exhibited markedly sharper cluster boundaries and consistently outperformed baseline methods across quantitative metrics including NMI, ARI, DB index and the Silhouette score. This demonstrates that introducing negative-mode mass spectra into pretraining establishes an essential foundation for comprehensively deciphering mode-specific fragmentation pathways across diverse ionization regimes.

**Figure 2.**
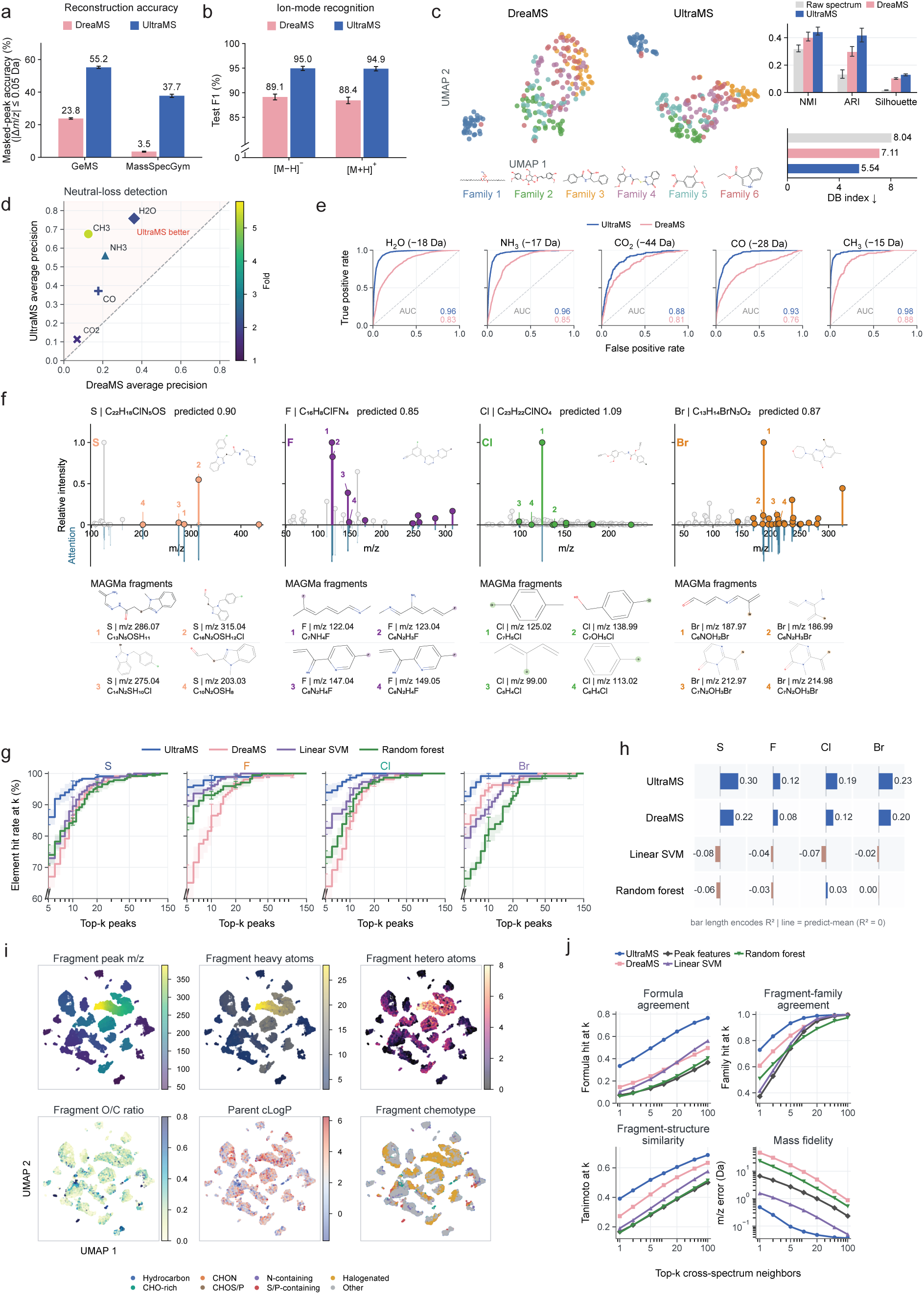
UltraMS demonstrates its superior mass spectral understanding ability by encoding meaningful chemical information at spectrum and spectral peak levels. **a**, Performance of masked spectral peak *m/z* re-construction on GeMS-A10 and MassSpecGym. Accuracy measures the proportion of masked spectral peaks whose *m/z* values are reconstructed within 0.05 Da of their true values after masking 15% of the input spectral peaks. **b**, *F*_1_ scores for positive- and negative-ion classification using spectrum-level embeddings on a balanced test set of 10,000 MSnLib spectra. **c**, Molecular structural organization of negative-ion spectra from SpectraVerse in the embedding space of various mass spectral foundation models, colored by molecular fingerprint-defined clusters. Normalized mutual information (NMI) and adjusted Rand index (ARI) measure agreement with these molecular clusters; silhouette scores measure their separation in the embedding space. The Davies–Bouldin index (DBI) measures within-cluster dispersion relative to between-cluster separation, with lower values indicating more compact and better-separated clusters. Error bars show 95% confidence intervals from bootstrap resampling within each molecular cluster. **d,e**, Evaluation of the recognition of characteristic neutral losses from MassSpecGym spectra, assessed by average precision (**d**) and receiver-operating-characteristic curves (**e**). **d** shows the UltraMS-to-DreaMS average-precision ratio. A neutral loss is treated as present when there exists a precursor-to-spectral-peak *m/z* difference that matches the mass of the corresponding neutral-loss type within 0.02 Da. **f**, Element-specific attention highlights spectral peaks containing S, F, Cl or Br. The upper panels show spectral peak intensities, and the lower panels show corresponding attention weights. The numbers displayed beside the spectral peaks correspond to the same numbers beside the substructures annotated by MAGMa. **g**, Element-specific spectral peak ranking. For each target element, hit rate at *k* measures the fraction of spectra in which at least one of the *k* highest-ranked spectral peaks has an annotated substructure containing that element. **h**, Performance of prediction of S, F, Cl and Br atom counts in the parent molecule, evaluated by the coefficient of determination (*R*^2^). **i**, Two-dimensional visualization of spectral peak-level embeddings. Each point represents an individual spectral peak (*n* = 60, 000), projected via UMAP based on its UltraMS embedding. Points are colored by the *m/z*, heavy-atom count, heteroatom count, O/C ratio and chemotype of the annotated spectral peak substructure, and by parent-molecule lipophilicity (cLogP), illustrating the spatial and chemical organization captured in the embedding space. **j**, Spectral peak matching across different spectra using embedding similarity. For each query spectral peak, spectral peaks from other spectra are ranked by their embedding similarity to the query. Formula and substructure-family hit rates measure the fraction of queries for which at least one of the top *k* spectral peaks has the same molecular formula or substructure-family annotation as the query, respectively. Structural similarity indicates the substructure ECFP4 fingerprint Tanimoto similarity.

The mass difference between the precursor ion and each product ion, known as a neutral loss, directly reflects the chemical nature of the uncharged moiety released during fragmentation. Because specific neutral losses usually map onto specific functional groups, they constitute a diagnostic vocabulary that chemists have long relied upon to read spectra and reconstruct potential fragmentation process [22, 23]. To evaluate whether the unsupervised pretraining of UltraMS captures such essential chemical prior in its embeddings, we tested its ability to predict the occurrence of a given neutral loss from the precursor ion. On MassSpecGym, UltraMS demonstrates greater chemical understanding by delivering superior identification, achieving a mean area under the receiver operating characteristic curve (AUC) of 94.3% compared with 82.5% for DreaMS across five common neutral losses (Fig. 2e), and a 1.7-to 5.4-fold higher average precision for each neutral loss type (Fig. 2d). This result indicates that unsupervised pretraining on raw spectra is sufficient for UltraMS to internalize the mechanistic chemical principles underlying neutral loss patterns, without explicit supervision on fragmentation rules. Predicting the elemental composition of a molecule directly from its tandem mass spectrum generally serves as a critical prerequisite for unknown metabolite identification [3]. While the presence of the four principal organic elements (C, H, O, N) can be reliably inferred by existing models, the identification of less frequent heteroatoms remains challenging yet carries broad practical significance (Supplementary Material Section S1.8). We employed an attention layer to directly predict the counts of four heteroatoms (S, Cl, F, Br) using UltraMS and DreaMS on MassSpecGym (see detailed method in Methods). Fig. 2f displays the learned attention map cases across molecules containing each of the four heteroatom types. The model adapts attention scores successfully to local peaks by assigning higher weights to peaks that truly contain the target element, while remaining robust to noise peaks without diverting attention. In the spectral peak-level evaluation, UltraMS consistently outperforms all other models across each element type by assigning at least one correct peak among the Top-*k* highest attention scoring candidates (Fig. 2g). In the overall evaluation, both UltraMS and DreaMS demonstrate substantial advantages over all baselines, while classical models operating on raw features yield near-zero or even negative *R*^2^ scores (Fig. 2h; see detailed discussion of performance in Supplementary Material Section S1.8). This underscores the critical role of latent chemical knowledge acquired during self-supervised pretraining in enabling models to decipher complex mass spectra.

Finally, apart from elemental detection, we aim to conduct a further probing tailored for the chemical characteristics of the fine-grained spectral peak-level embeddings generated by different foundation models. Specifically, we collect spectral peaks that can be reliably annotated from MSnLib. These spectral peaks are fully annotated for their substructures with the MAGMa algorithm [22], yielding 60,000 reliably annotated spectral peaks in total (see detailed method in Supplementary Material Section S1.8). We first depict UltraMS’s spectral peak-level embedding landscape using UMAP [24] and color them from different perspectives (Fig. 2i). We observe that fragments represented by these spectral peaks form meaningful local manifolds that distinguish chemically similar groups, such as clearly layered *m/z* regions accompanied by different numbers of heavy atoms, the cLogP and chemotype of fragments, and heteroatoms and the O/C ratio, showing that UltraMS presents a well-defined embedding space encompassing diverse chemical information implied in spectra. Nearest Top-*k* neighbor ranking using UltraMS preserves fragments with the same formula clustered together among the neighbors and maintains the best structural similarity while keeping the most consistency of fragment family and least mass deviation with substantially higher fidelity than other baselines (Fig. 2j). Additionally, we observed that UltraMS retains high embedding similarity among chemically equivalent substructures represented by spectral peaks that stably occur across varying collision energies for the same compound, demonstrating its ability to capture canonical and persistent fragmentation patterns for a given molecule via its spectral peak-level embeddings (Supplementary Fig. 3). We also demonstrate that the spectral peak-level embeddings encoded by UltraMS serve as an effective bridge to identify shared substructures across tandem mass spectra of different compounds, revealing a promising capacity for annotation transfer from well-characterized reference standards to unannotated molecules (Supplementary Fig. 4). Overall, our extensive benchmarks and results elucidate the comprehensiveness and efficacy of UltraMS as a general method to tackle diverse chemical tasks with advanced performance.

### 2.3 Pretrained spectr al representations enable accurate molecular identification and fine-grained structural reasoning

Inference of molecular identities from their corresponding spectra serves as the tunnel from unseen microscopic particles to human-interpretable chemical objects, offering substantial importance for uncovering key components inside samples across various biological and chemical research areas and practical applications, such as novel species metabolite discovery, clinical diagnostics, environmental pollution detection, and drug and toxicant screening [2]. We applied UltraMS to essential molecule identification tasks via two basic paths: spectrum-to-molecule direct prediction and spectrum-to-spectrum library search. Due to the incompatibility between raw spectra and molecular representations such as SMILES strings, we introduced the pretrained molecular foundation model ChemBERTa [25] to encode molecules into embedding representations for spectrum-to-molecule prediction (Methods). The original MSnLib represents one of the most authoritative mass spectral libraries [26], and is well suited as a reference source for library search or as an identification performance evaluation platform under diverse ion modes. Therefore, we chose MSnLib as the major benchmark and partitioned the complete library by molecular formula into training, validation and test sets, which we release through Zenodo (DOI: 10.5281/zenodo.23963489; see detailed method in Supplementary Material Section S1.8). Additionally, to conduct comprehensive testing we also report the performance of UltraMS across various general public benchmarks including NPLIB1 [9, 10] and MassSpecGym [27] and compare with diverse models.

We primarily evaluate different models on spectrum-to-molecule identification performance under two distinct protocols. The first strictly requires the model to identify the ground truth mixed within large isomer candidate sets averaging approximately 2,804 candidates from a single provided spectrum. The second simulates a more realistic identification process: to determine a molecule’s structure, chemists generally do not merely measure its mass spectrum under one specific collision energy but rather gather sub-structural evidence across different molecule collision conditions or different instruments to decide the correct structure. Therefore, we collected all spectra of the same molecule across collision conditions for each molecule as a basic test unit, where models primarily independently recognize the molecule from each spectrum and then vote for the best candidate to determine its true identity. Overall, UltraMS achieves superior results under both setups, with notable improvement in top-*k* accuracy, reaching 27.15% Top-1 accuracy in single-spectrum identification, compared with 20.25% for DreaMS and 17.92% for the best-performing other baseline; this advantage broadens further under multiple-spectrum voting, with 42.56% for UltraMS versus 33.49% for DreaMS and 24.35% for the best-performing other baseline (Fig. 3a). Nevertheless, even though UltraMS preserves its superior ranking with the distribution of the true molecule’s similarity scores concentrated closer to the top for the corresponding spectra, single-spectrum identification accuracy reaches only 43.54% at Top-20 and merely 56.92% at Top-100, whereas multiple-spectrum voting substantially increases the corresponding identification accuracy to 64.87% and 76.41%, respectively (Fig. 3b), highlighting the difficulty of resolving the correct structure from limited and noisy single-spectrum evidence and emphasizing the necessity of realistic molecule identification using multi-condition spectra. Moreover, following each test molecule across its spectra, we found that UltraMS recovers the correct structure from at least one acquisition condition for the majority of molecules, more than any other model (Supplementary Fig. 6; see detailed discussion in Supplementary Material Section S1.3). We further demonstrate that UltraMS consistently maintains superior performance over a broad set of baselines across diverse molecular identification paradigms, such as fingerprint-based discrimination and benchmark challenges of varying difficulty levels (Supplementary Fig. 5).

**Figure 3.**
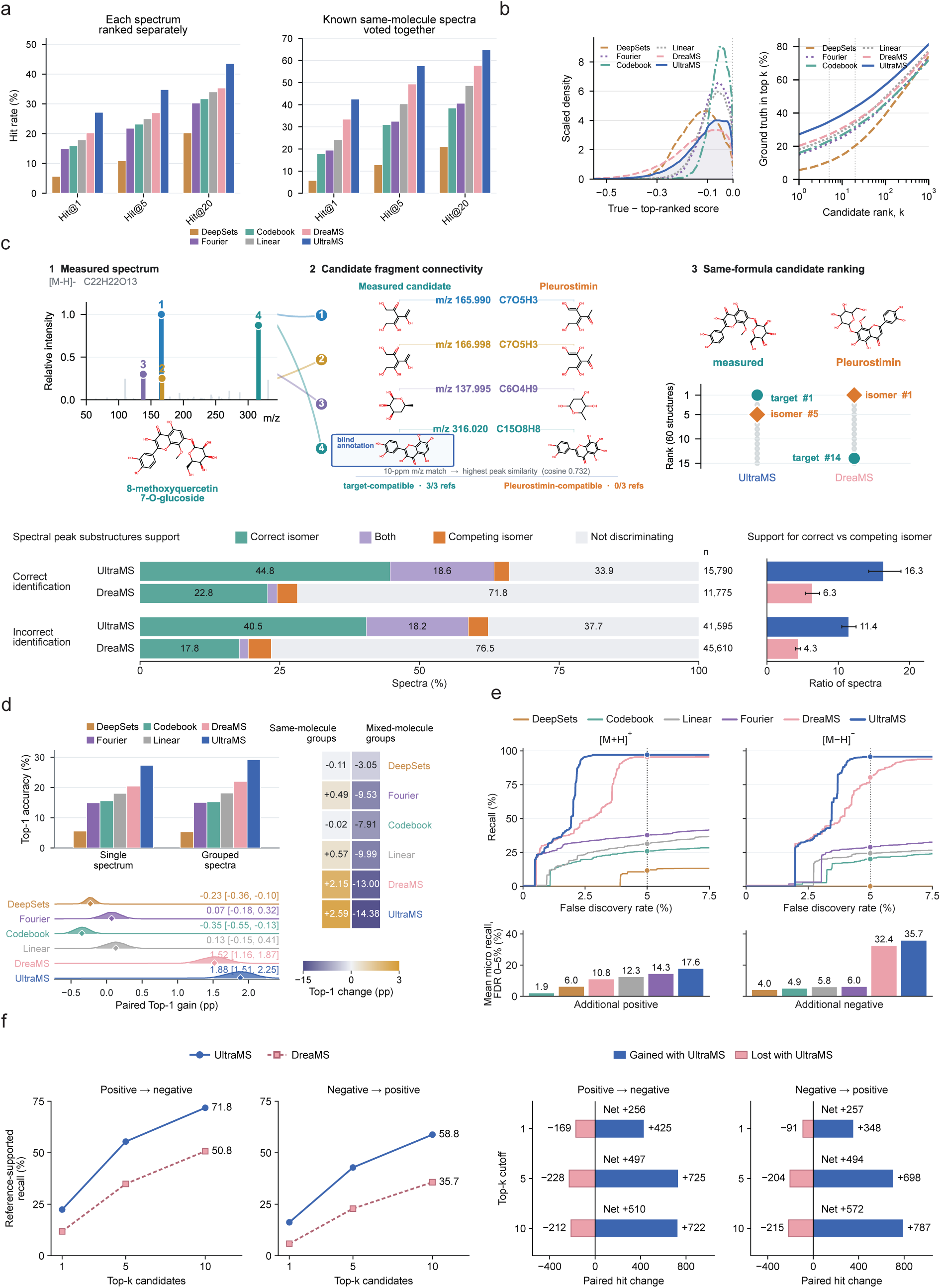
Learned chemical semantics enable molecular identification with fine-grained interpretation. UltraMS identifies molecules by ranking candidate structures and searching spectral libraries; its spectral peak-level embeddings also transfer a substructure annotation that helps distinguish same-formula isomers. **a**, Evaluation of molecular identification from tandem mass spectra. Left panel: Models rank candidate molecules from individual MSnLib spectra (single-spectrum mode). Right panel: When multiple spectra originate from the same molecule, a consensus voting strategy aggregates their individual rankings to yield a unified identification. Hit@*k* indicates the percentage of queries where the ground-truth molecule ranks within the top *k* predictions. **b**, Error analysis and cumulative identification accuracy. Left panel: Distribution of similarity score margins, calculated as the cosine similarity of the ground-truth molecule minus that of the top-ranked incorrect candidate (more negative values indicate stronger model preference for incorrect candidates). Right panel: Cumulative single-spectrum identification accuracy as a function of candidate rank. **c**, Case study of structural isomer discrimination. Top panel: UltraMS correctly ranks 8-methoxyquercetin 7-*O*-*β*-D-glucopyranoside above its structural isomer Pleurostimin (ranked first by DreaMS). Central panel: Substructures annotated by MAGMa across the two candidate structures. UltraMS transfers a discriminative substructure annotation from mass-matched reference peaks to the measured peak at *m/z* 316.020 using peak-level embeddings, independent of the query molecule’s ground-truth identity. Supplementary Fig. 7 shows the property distributions of the top 20 candidates, with outlined symbols marking the ground-truth structure. TPSA, topological polar surface area; cLogP, calculated octanol–water partition coefficient. Lower panels: Substructure-level discrimination of structural isomers. Percentage of MSnLib test spectra where transferred peak annotations support the correct structure over the top-ranked competing isomer, evaluated across correctly and incorrectly identified queries for UltraMS and DreaMS. **d**, Molecular identification via unsupervised spectral clustering. Spectra are clustered based on ion adduct, precursor *m/z*, retention time, and cosine similarity, followed by consensus voting within each cluster to determine molecular identities. Upper left panel: Comparison of Top-1 identification accuracy between single-spectrum and clustered-spectrum evaluations. Lower left panel: The distribution of paired accuracy changes shows the performance variation using bootstrap resamples, calculated as its cluster-based Top-1 accuracy minus its single-spectrum accuracy. Right panel: Effect of clustering errors, showing that incorrectly grouping different molecules into the same cluster causes a substantially larger accuracy drop than the gains achieved by correctly clustered spectra. Brackets denote 95% confidence intervals derived from cluster bootstrap resampling; pp, percentage points. **e**, Evaluation of spectrum-to-spectrum library search under empirical false discovery rate control. Upper panels: Empirical FDR–recall curves for [M + H]^+^ and [M *−* H]*^−^*. FDR is defined as the proportion of accepted identifications that are incorrect, and recall denotes the fraction of library-matched queries correctly identified. Dotted lines mark the 5% FDR threshold. Lower panels: Mean recall evaluated across the 0–5% FDR window for additional positive- and negativeion adducts. **f**, Reciprocal cross-ionization molecule identification via spectrum-to-spectrum library search. Comparison of identification accuracies between UltraMS and DreaMS across positiveand negative-ion spectra sets.

Figure 3c showcases a spectrum-to-molecule identification example of the molecule 8-Methoxyquercetin 7-*O*-*β*-D-glucopyranoside, which was reported as a putatively identified differential metabolite of *Thesium chinense*, a medicinal plant whose standardized preparation is clinically used for inflammatory respiratory disorders [28, 29]. This molecule serves as an important profiling marker for *Thesium chinense* bioactivity assessment and geographical discrimination in quality evaluation [28]. However, isomers of methoxylated flavonoid *O*-glycosides share highly similar fragmentation patterns, and localizing their substituents often requires retention-time and spectral matching against precious purified isomer standards [30], or 2D NMR [31]. The spectral peak annotation algorithm MAGMa was run to annotate the spectra using the respective presumed structures of the true molecule and one isomer called Pleurostimin. Annotation results further demonstrate a challenging case that even spectral peak-level substructure differences are only confined to a single oxygen atom position. In the test, UltraMS primarily detected the correct structure and ranked it at top-1 among numerous formula-identical candidates exhibiting extremely similar and hard-to-differentiate chemical properties. In comparison, DreaMS was misled and ranked another isomer, Pleurostimin, as its top selection and placed the true structure largely back at position 14. Additionally, UltraMS further provides fine-grained clues by assigning the spectral peak at 316.020 *m/z* to the correct substructure with the precise essential oxygen atom combination position using its peak embedding from peak-level library search without knowing the final molecular structure, confirming the correct molecule and unlocking chemical fragmentation interpretation simultaneously. We further extended this substructure-level analysis to isomer discrimination across the full MSnLib test set (Fig. 3c bottom panel). On the MSnLib test set, we evaluated how selectively transferred spectral peak annotations distinguish the correct structure from competing isomers (Methods). Among spectra whose corresponding molecular identities were correctly identified by each model, 44.8% showed support exclusively for the correct molecular identity with UltraMS, compared with 22.8% with DreaMS. This advantage persisted even among incorrectly identified spectra, with corresponding proportions of 40.5% and 17.8% (see detailed discussion of performance in Supplementary Material Section S1.3). We next examine a stricter multiple-spectrum molecule identification protocol based on unsupervised spectral clustering across diverse models. In this evaluation, the model is blind to the molecular attribution of each spectrum and is required to first determine whether multiple spectra belong to the same molecule before utilizing all spectra within one group to select the best candidate (Fig. 3d). UltraMS demonstrates the greatest improvement at 1.88 percentage points (95% group-bootstrap CI: 1.51–2.25), compared with 1.52 percentage points for DreaMS (95% CI: 1.16–1.87). Notably, none of the other non-pretrained baselines achieve stable performance gains and instead degraded their accuracy, further highlighting the importance of versatile chemical knowledge implicitly acquired during pretraining for enabling more robust identification. The right panel contrasts performance under correct spectral grouping versus erroneous merging of spectra, indicating that misalignment substantially degrades accuracy across all models and largely outweighs the positive effect of correct grouping. These results emphasize the criticality of spectrum-to-spectrum clustering accuracy for multiple-spectrum identification.

Spectrum-to-spectrum library search treats molecule identification as a retrieval task, in which the cosine similarity is computed directly between a query spectrum and the reference spectra of comparable precursor mass within a predefined library, representing the prevailing paradigm for molecule identification. However, operating directly on raw spectra leaves it exposed to unresolved ambiguity arising from variation in acquisition conditions such as various collision energy that drives the molecule along distinct fragmentation pathways [32], leading to poor performance and low recall, particularly under cross collision energy settings. We evaluate model performance on the public MSnLib spectra library across diverse ion modes (Methods). UltraMS achieves promising recall while maintaining the lowest false discovery rates under different acceptance thresholds compared with the other baselines. Notably, both pretrained spectral language models, UltraMS and DreaMS, are able to achieve maximum recalls of 97.13%/95.70% and 95.44%/93.73% for [M+H]^+^/[M − H]^−^, respectively, while the best model without pretraining delivers maximum recalls of merely 74.54%/71.99%. Furthermore, these advantages also extend to more diverse ion modes. With the average FDR constrained to 0–5%, UltraMS achieved mean micro recalls of 17.58% and 35.71% for additional positive- and negative-ion adducts, respectively, compared with 14.25% and 32.38% for the best-performing alternative (Fig. 3e). Furthermore, in a more challenging cross-collision-energy searching task, UltraMS maintained over 92% Recall@1 across both ionization modes (Supplementary Fig. 8; see detailed discussion of performance in Supplementary Material Section S1.3). Positive- and negative-ion spectra provide complementary views of a molecule, yet their distinct ion species and fragmentation pathways often result in minimal peak overlap, making the same chemical identity difficult to recognize across polarities. We therefore tested whether the representations learned by UltraMS could support spectrum-to-spectrum library search across ionization polarities (Fig. 3f; see detailed method in Supplementary Material Section S1.8). Among the 2,421 positive-ion and 2,476 negative-ion queries whose correct molecules were present in the opposite-polarity library, UltraMS ranked the correct molecule first for 22.4% and 16.2% of queries, respectively, compared with 11.9% and 5.9% for DreaMS. Within the top ten candidates, UltraMS ranked the correct molecule for 71.8% of positiveion and 58.8% of negative-ion queries, compared with 50.8% and 35.7% for DreaMS (see detailed discussion of performance in Supplementary Material Section S1.3). A query-by-query comparison revealed that UltraMS substantially expanded identification coverage over DreaMS, with top-ten identifications gained outpacing those lost by more than three-to-one across both ionization modes (722 versus 212 in positive mode, net +510; 787 versus 215 in negative mode, net +572; Fig. 3f, right panel). These results demonstrate that full-polarity pretraining enables UltraMS to learn shared, polarity-invariant representations of chemical structure, allowing positive- and negative-ion data to be unified within a shared coordinate space via shared molecular identity rather than treated as isolated repositories. Collectively, pretrained on a universal mass spectrometry corpus spanning diverse practical scenarios, UltraMS learns the underlying chemical language expressed from molecular fragmentation and encodes spectra acquired under a wide range of conditions into chemically meaningful embeddings. Its spectral peak-level representations further elevate molecular identification from one-shot spectral matching toward mechanistic chemical reasoning, establishing pretrained spectral foundation models as a versatile cornerstone for broad downstream applications including metabolite discovery, clinical diagnostics, and environmental screening.

### 2.4 Grounding molecular identification with spectral peak-level substructure annotations

A molecule identification proposed from a tandem mass spectrum becomes chemically convincing when the spectrum itself can be explained. Deep learning models for molecular identification typically return a similarity score or a ranked list of candidate molecules without interpreting the underlying spectral peaks, leaving chemists in the dark about how individual spectral peaks support the proposed molecule and limiting their ability to verify the result using chemical logic. We therefore asked whether a large-scale pretrained model can, beyond assigning a molecular identity, open that molecule identification to chemical interpretation by revealing what each spectral peak represents (Fig. 4). As demonstrated across our analyses (Fig. 2i,j), the peak-level embeddings of UltraMS inherently encode meaningful chemical information. Leveraging this generalizable representation, we reasoned that this learned capacity could be systematically scaled to annotate individual spectral peaks: a query peak can be assigned the substructure of the reference peak whose embedding is most similar to its own. We evaluated this approach on the original MSnLib test partition (see detailed method in Supplementary Material Section S1.4). Spectral peaks of the query and reference spectra were annotated by MAGMa using their known parent molecules. For this nearest-neighbor annotation approach to succeed, the underlying embedding space must be organized by chemical substructure—meaning that spectral peaks representing similar substructures should lie close together (Fig. 4a). Specifically, within each query spectrum, we compared the cosine similarity of peak-level embeddings against the Tanimoto similarity of their annotated substructures. UltraMS achieved a mean Pearson correlation coefficient (PCC) of 0.379 (95% CI 0.374–0.383), outperforming DreaMS 0.077 (0.073– 0.082) across 95.5% of parent molecules (4,536 of 4,749). We then directly transferred substructure annotations to each query peak from the reference peak with the highest embedding similarity across the entire library.

**Figure 4.**
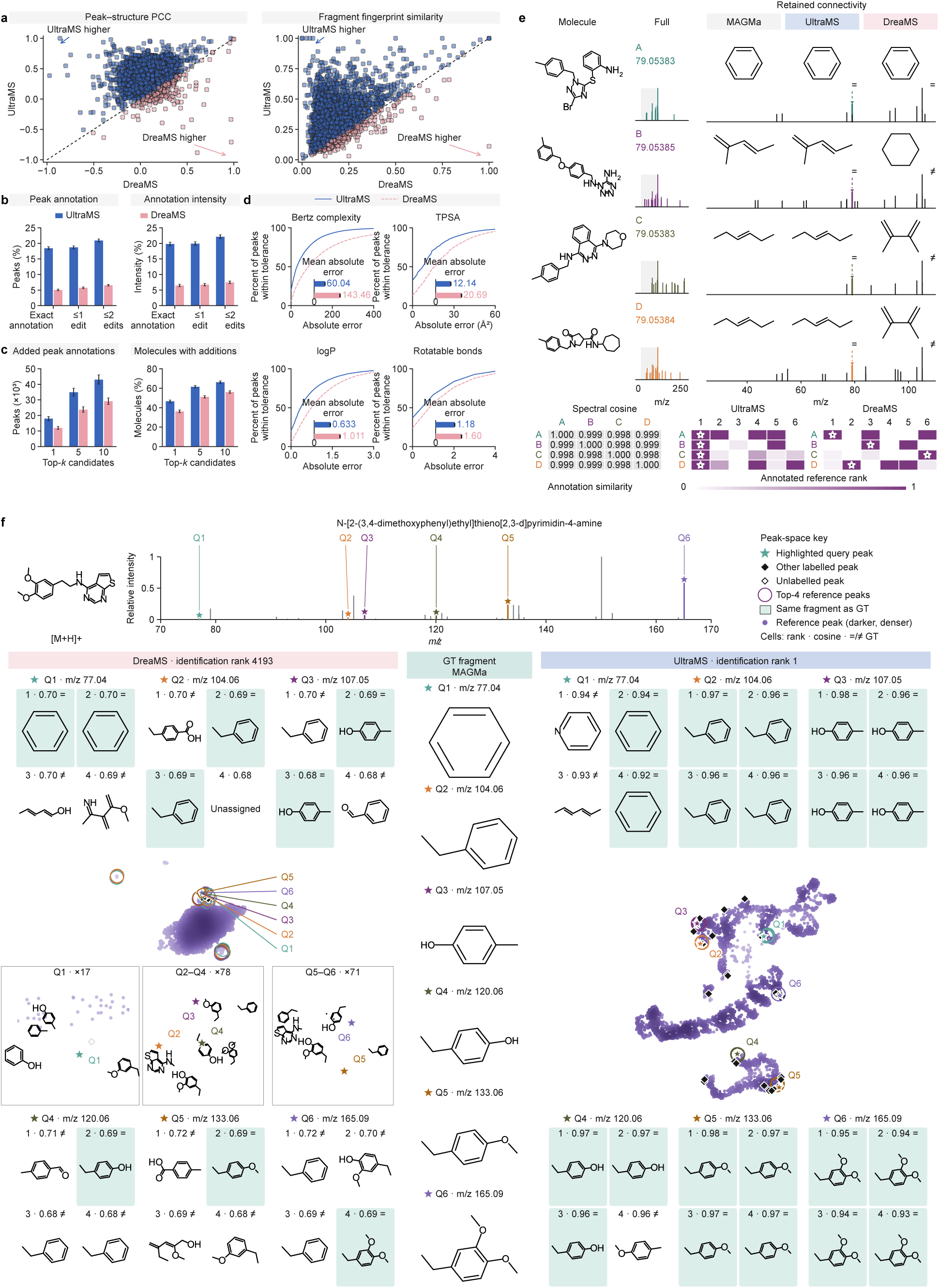
Spectral peak embeddings enable chemical interpretation of tandem mass spectra. **a**, Molecule-level comparisons of the correlation between within-spectrum embedding cosine and substructure Tanimoto similarities (left; *n* = 4, 749 molecules), and Tanimoto similarity between transferred and MAGMa annotations (right; *n* = 4, 764). Each point represents one molecule. **b**, Fractions of spectral peaks and their original intensity explained by exact annotations or annotations within one or two atom or bond edits. **c**, Additional annotations beyond MAGMa and the percentage of test molecules with additions, using the top 1, 5 or 10 references from distinct molecules. Transfer annotation acceptance required connectivity and bond types supported by the query molecule and agreement with the observed *m/z* within 10 ppm. **d**, Cumulative distributions of absolute errors in Bertz complexity, topological polar surface area (TPSA), logP and rotatable bond count. **e**, Four highly similar spectra with a shared spectral peak near *m/z* 79.054 and three MAGMa-annotated substructures. Similarity matrices show spectral cosine and annotation similarity; stars mark the first identical substructure among annotated references in retrieval order. **f**, UMAP projections of query and reference spectral peak embeddings for an example molecule, alongside the four highest-ranked reference annotations for each of six query spectral peaks. Stars represent query spectral peaks.

Evaluating these predicted annotations against the MAGMa annotations, UltraMS achieved a mean Tanimoto similarity of 0.295 (95% CI 0.291–0.299), outperforming DreaMS 0.157 (0.154–0.160) across 92.7% of parent molecules (4,414 of 4,764). We next evaluated the transferred annotations under the strictest standard: whether they faithfully reveal the ground-truth substructures provided by MAGMa (Fig. 4b). Across 599,185 query spectral peaks evaluated by both models, UltraMS assigned the exact substructure to 18.4% of spectral peaks (95% CI, 18.0–18.9%), outperforming DreaMS (5.0%; 95% CI, 4.8–5.3%) by 3.7-fold. This advantage persisted when spectral peaks were weighted by their intensity or when the transferred and true substructures were allowed to differ by up to two atom or bond edits (Fig. 4b; see detailed discussion of performance in Supplementary Material Section S1.4).

Crucially, we demonstrate that the structural representations encoded by pretrained UltraMS are able to effectively serve as a bridge to scale and amplify the annotation capabilities of existing domain tools. Specifically, we leveraged existing MAGMa annotations across the reference library to transfer substructure interpretations to spectral peaks that MAGMa left unannotated (Fig. 4c; see detailed method in Methods and Supplementary Material Section S1.4). Among the top ten candidates retrieved from distinct reference parent molecules, UltraMS yielded 28,617 additional peak annotations—a 6.6-fold increase over DreaMS (4,309 annotations). Furthermore, UltraMS obtained at least one additional annotation for 57.5% of test parent molecules (95% confidence interval [CI], 56.3–58.9%), compared with only 19.2% for DreaMS (95% CI, 18.1–20.3%). Moreover, we compared physicochemical descriptors between transferred and MAGMa-annotated substructures (Fig. 4d). UltraMS achieved lower macro-averaged mean absolute errors across all four descriptors (see detailed discussion of performance in Supplementary Material Section S1.4). Together, these results demonstrate that UltraMS-guided annotation transfer significantly expands peak-level chemical interpretability while preserving structural fidelity. In tandem mass spectrometry molecular identification, a particularly challenging scenario arises when spectral peaks share identical nominal *m/z* values within nearly indistinguishable spectral backgrounds (Fig. 4e). To intuitively explore how UltraMS navigates these deceptive regimes, we probed four spectra with extreme pairwise cosine similarities (0.9977–0.9994) that all share a prominent peak near *m/z* 79.0538. UltraMS correctly identified the ground-truth substructure at Rank 1 across all four cases. In contrast, DreaMS placed the true annotation at Rank 1 in only a single instance; for the remaining three targets, the correct substructures were relegated to ranks 2, 3, and 6. This shows that UltraMS genuinely decodes the contextual chemical language within spectra, interpreting the underlying molecular logic rather than only relying on superficial spectral resemblance, and thereby elevating peak interpretation to definitive structural discrimination. We further exhibited the latent embedding space for an individual query spectrum alongside its reference chemical evidence (Fig. 4f). In the UMAP projection of peak representations, UltraMS organized the six high-intensity query peaks into distinct, chemically coherent neighborhoods. By contrast, DreaMS collapsed chemically divergent substructures into a nearly undifferentiated cluster. Across the four top-ranked reference peaks for each of the six representative query peaks, UltraMS achieved 21 out of 24 concordant annotations with the ground-truth MAGMa substructures, whereas DreaMS achieved only 9 of 24. Furthermore, this spectral-peak-level structural accuracy translated directly into parent molecule identification: UltraMS prioritized the true molecule at Rank 1, whereas in DreaMS the correct candidate was lost at Rank 4,193. Together, this demonstrates a compelling paradigm: by bridging spectral representation learning with grounded chemical references, UltraMS is able to transform spectral identification from an empirical, blind matching process into an interpretation-driven, evidence-backed deduction.

### 2.5 UltraMS deciphers clinical plaque metabolomes to reveal phenotype-associated metabolic signatures

We further confirmed the benefits of using UltraMS in a clinical cohort for metabolic research. We used data from a publicly available carotid-plaque cohort consisting of 72 participants, including 30 symptomatic participants who experienced stroke-related symptoms within one month before surgery and 42 asymptomatic participants who experienced no such event during the preceding six months [33]. The plaque LC–MS/MS data comprised 13,116 unique molecular features, including 8,457 positive-ion and 4,659 negative-ion features derived from the human carotid-plaque LC–MS/MS results. We first visualized the overall UltraMS embedding manifold from multiple perspectives by coloring points according to ionization polarity, precursor *m/z*, retention time, feature prevalence, MS2 support and feature intensity. Notably, UltraMS spectral embeddings exhibited distinctive clustering properties across these key mass-spectral attributes, organizing into coherent local neighborhoods characterized by precursor *m/z* or retention time, and even revealing regions enriched in molecular features with greater prevalence, stronger MS2 support and higher feature intensity across samples (Fig. 5a). We next sought to investigate whether molecular features across symptomatic and asymptomatic plaque groups exhibited phenotype-dependent organization via their spectral embedding projections in the UltraMS representation space. We then observed that molecular features with the same phenotype enrichment in the corresponding group also tended to accumulate within distinct local regions rather than being randomly dispersed (Fig. 5b). We subsequently showed that this neighborhood phenomenon robustly extended to individual runs. By examining local regions of molecular features in individual LC–MS runs, we demonstrated that local embedding clusters contained group-specific features and exhibited consistent abundance directionality (Fig. 5c). By mapping two representative embedding clusters comprising molecular features differentiated by *m/z* back to their native chromatographic retention-time coordinates, we observed that these molecular features exhibited markedly more similar retention times than those outside the cluster (Fig. 5d), demonstrating UltraMS’s ability to faithfully reflect intrinsic chemical characteristics, including physicochemical polarity and structural relatedness, as evidenced by the convergent retention times.

**Figure 5.**
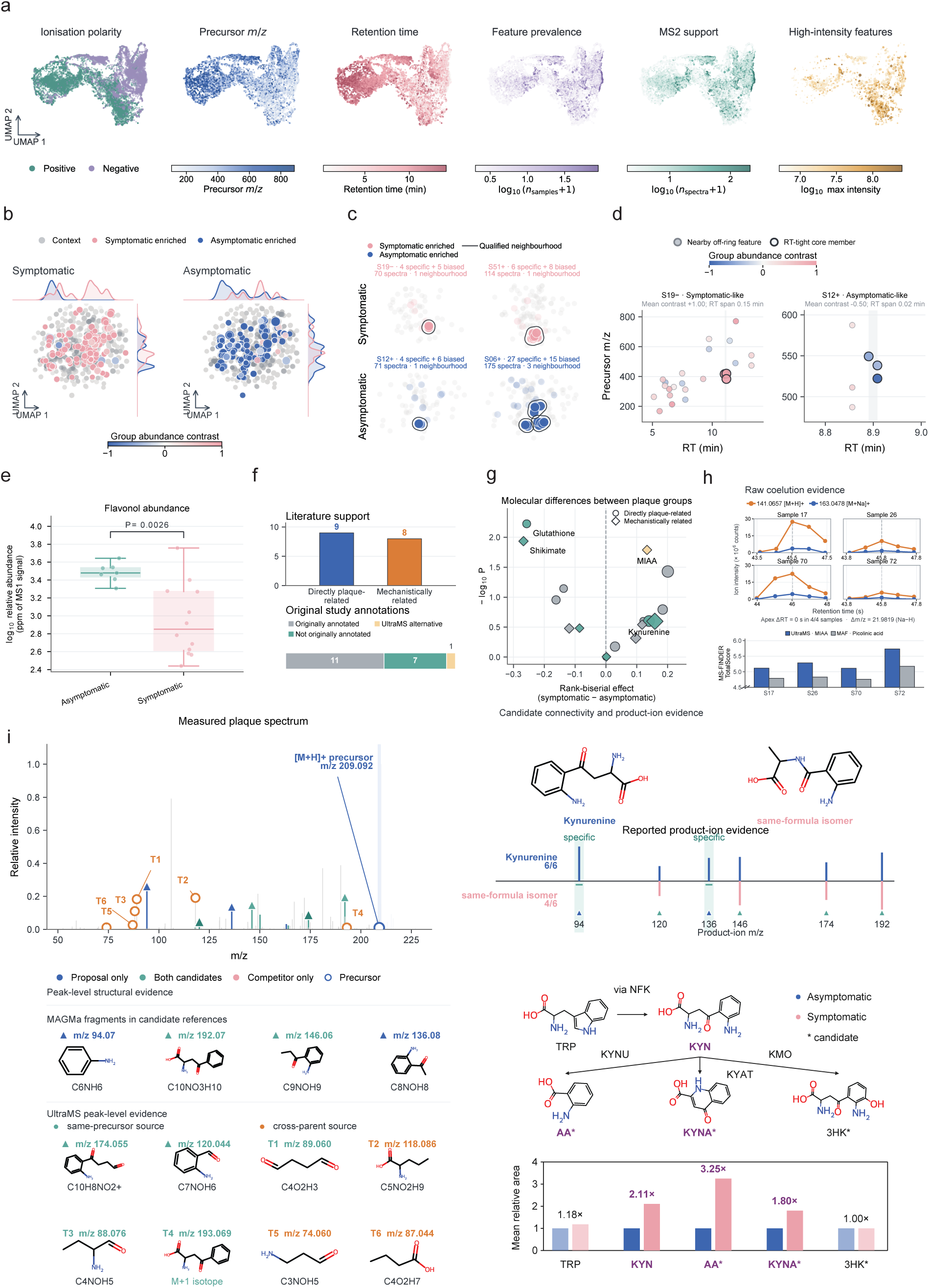
UltraMS enables phenotype-specific plaque metabolomics and elucidates structurally unrevealed metabolites through interpretable peak-level substructure attribution. UltraMS transforms untargeted tandem mass spectrometry data from clinical plaque specimens into a coherent representation space, resolving phenotype-associated metabolic signatures between symptomatic and asymptomatic patients. Orthogonal validation via extracted-ion chromatography and in silico fragmentation, coupled with peak-level embedding transfer, enables robust structural elucidation of clinically relevant metabolites and discriminates critical structural isomers. **a**, Organization of clinical plaque spectra in the UltraMS representation space. Two-dimensional UMAP projection of 13,116 molecule features. Individual panels display molecule features color-coded by ionization polarity, precursor *m/z*, retention time, sample prevalence, MS^2^ spectral support, and MS1 signal intensity. **b**, Distribution of plaque molecule features by phenotype-associated abundance across 11,351 feature-linked spectra. Points are color-coded by the relative abundance contrast of normalized MS1 intensity, defined as (*µ*_symptomatic_ *− µ*_asymptomatic_)*/*(*µ*_symptomatic_ + *µ*_asymptomatic_), with pink and blue indicating enrichment in symptomatic and asymptomatic plaques, respectively. Point size scales with MS1 intensity, and marginal density plots show feature distributions along the UMAP coordinates. **c**, Phenotype-enriched spectral neighborhoods in representative plaque samples. Four representative samples display localized clusters of molecule features selectively enriched in or specific to the indicated clinical phenotype. Outlines highlight these neighborhoods within the UltraMS representation space. **d**, Retention time and precursor *m/z* profiles for two selected spectral neighborhoods. **e**, Relative abundance of flavonols uncovered by UltraMS across plaque samples with detectable signals. Comparison of MS1 signal intensity between asymptomatic (*n* = 7) and symptomatic (*n* = 12) plaque samples. In the box plots, center lines represent medians, box limits indicate interquartile ranges (IQRs), whiskers extend to 1.5 *×* IQR, and individual points denote distinct samples. *P* = 0.00262 (unadjusted two-sided Mann–Whitney *U* -test). **f**, Literature validation of characterized metabolites. Upper panel: 17 characterized metabolites categorized by supporting evidence from carotid plaque studies and broader vascular literature. Lower panel: Categories of 19 metabolite assignments: annotations consistent with the original study, novel annotations uncovered by UltraMS, and UltraMS-proposed revisions to the original study’s identification assignments. **g**, Phenotypic associations of the 17 literature-supported candidate metabolites. Participant-level rank-biserial effect size (*x*-axis; positive values denote higher abundance in symptomatic plaques) and statistical significance (*−* log_10_ *P*, *y*-axis; unadjusted two-sided Mann–Whitney *U* -tests). Point size scales with the number of supporting spectra, shape indicates the type of literature support, and colors correspond to the annotation categories defined in **f**. **h**, Reassessment of an ambiguous metabolite assignment through orthogonal evidence. Extracted-ion chromatograms (EICs) across four representative plaque samples demonstrate co-eluting ions at *m/z* 141.0657 and 163.0478, consistent with protonated ([M+ H]^+^) and sodiated ([M+ Na]^+^) adducts of methylimidazoleacetic acid (MIAA), as proposed by UltraMS. This structural revision accounts for a signal originally assigned as an ammonium adduct of picolinic acid. Across all four spectra, in silico fragmentation via MS-FINDER yields consistently higher compatibility scores for MIAA than for picolinic acid. **i**, Peak-level substructure attribution and isomer discrimination for UltraMS-identified kynurenine. The experimental plaque spectrum matches all six characteristic product ions of kynurenine, compared with only four for the top-ranking structural isomer sharing the same molecular formula. Lower left panel: Substructure annotations transferred to query spectral peaks via UltraMS peak-level embeddings. Labels T1–T3 and T5–T6 denote transferred substructure hypotheses, whereas T4 corresponds to an isotopic peak. Pathway diagram: Summary of the kynurenine pathway derived from an independent carotid plaque cohort [45].

We then probed whether the phenotype-associated organization of plaque spectra encoded by UltraMS could further reveal clinically informative molecular differences between symptomatic and asymptomatic plaque samples. We first encoded the spectra corresponding to molecular features using UltraMS and searched them against a reference-library collection comprising three structure-annotated public datasets: SpectraVerse, MassSpecGym and MSnLib. This analysis ultimately identified 506 high-confidence molecular features with structural annotations (see detailed method in Methods). We then classified these molecular features into chemical classes using NPClassifier [34] according to their annotated structures. We observed that flavonols produced the strongest phenotype-associated separation across all eligible chemical classes, with a 4.2-fold higher median relative abundance in asymptomatic detections than in symptomatic detections (two-sided Mann–Whitney *P* = 0.00262; Fig. 5e). We further found that this UltraMS-derived class-level clue was consistent with independent human and experimental evidence: higher flavonol intake has been associated with lower carotid-plaque burden [35] and the flavonol quercetin suppresses plaque-associated inflammatory signaling and plaque development in experimental models [36]. To move beyond broad chemical classes and focus on specific, mechanistically interpretable molecular identities, we performed a rigorous structural inspection of the 506 annotated molecular features. We then conducted targeted literature searches, yielding 17 biologically interpretable molecules. According to the scope of the supporting literature, nine were directly plaque-related, with evidence from plaque-focused studies, whereas eight were mechanistically related through experimental carotid or other vascular studies (Fig. 5f). Furthermore, we compared our molecular annotations with those reported in the original MTBLS12303 study [33], showing that nearly all of our annotation results aligned precisely with the existing annotations, with identical structures for nine molecules. Additionally, seven molecules were newly annotated by UltraMS and had not been uncovered in the original study. We next examined how these molecular candidates differed between plaque phenotypes by quantifying their relative abundance changes across subgroups, where negative values indicated lower abundance in the symptomatic cohort (Fig. 5g). We found that glutathione showed the clearest association with asymptomatic plaques among the seven additional annotations revealed by UltraMS. It was detected by UltraMS in 12 of 42 asymptomatic plaques but only 1 of 30 symptomatic plaques (abundance difference, 0.257; *P* = 0.00595). Further literature investigation showed that its higher abundance in asymptomatic plaques was consistent with the established protective effect of glutathione in plaque biology [37–40] (Supplementary Note S1.5). Additionally, shikimate (shikimic acid) was detected by UltraMS in 15 of 42 asymptomatic plaques but only 3 of 30 symptomatic plaques (see detailed discussion in Supplementary Material Section S1.5).

We present a particularly revealing case in which UltraMS produced a result that differed from the original molecular annotation reported in the MTBLS12303 study. We found that the corresponding molecular feature was detected in four symptomatic plaque samples but in none of the asymptomatic plaque samples. It had originally been assigned to picolinic acid [*M* + NH_4_]^+^, whereas UltraMS selected methylimidazoleacetic acid (MIAA) [*M* + H]^+^ as a better structural annotation than picolinic acid from its reference library (Fig. 5g). We therefore returned to the original data to explore which structure better explained the complete ion pattern. Here, we observed that the ions at *m/z* 141.0657 and 163.0478 co-eluted and reached the same chromatographic apex in all four samples (Fig. 5h), closely matching the expected co-elution pattern of coexisting protonated and sodium-adduct ions. Their mass difference of 21.9821 Da aligned well with the theoretical 21.9819-Da difference between [*M* + H]^+^ and [*M* + Na]^+^, identifying them as a characteristic ion pair from the same approximately 140.058-Da neutral molecule, which can be well explained by MIAA [*M* +H]^+^ at *m/z* 141.0657 and MIAA [*M* +Na]^+^ at *m/z* 163.0478. In contrast, the original picolinic acid interpretation failed to explain this paired signal (see detailed discussion in Supplementary Material Section S1.5). This reassignment was further supported orthogonally by MS-FINDER [41], which consistently favored MIAA over picolinic acid across all four plaque spectra (Fig. 5h; see detailed discussion in Supplementary Material Section S1.5). Additionally, a more direct biological connection is evident: MIAA is a stable product of histamine turnover [42], and the corresponding histamine signal has been linked to symptomatic carotid disease in humans [43] and carotid disease in experimental models [44]. This case illustrates how UltraMS can move beyond proposing annotations and utilize the inherent mass spectral chemistry to rethink an ambiguous existing assignment and prioritize a more reliable molecular explanation.

Finally, we aimed to probe whether UltraMS spectral peak embeddings could provide a deeper structural explanation for a molecule without a corresponding annotation in the original study (Fig. 5i). In particular, kynurenine (KYN) was confidently and consistently prioritized by UltraMS across five plaque spectra as a noteworthy metabolic signal, despite not being revealed in the original study. We attempted to evaluate the rationality of this annotation by interpreting the observed fragmentation pattern using MAGMa. However, we found that only four spectral peaks could be clearly annotated by MAGMa, leaving much of the measured spectrum without an explicit peak-level substructure explanation. Notably, we also found that several of these unexplained peaks represent known signature peaks of identifying KYN. To overcome this challenge, we exploited the spectral peak embeddings learned by UltraMS as an annotation-transfer medium to interpret the remaining high-intensity unannotated peaks (Methods). UltraMS elevated the interpretable intensity to 68.4%, representing a 5.1-fold increase compared with the initially interpretable MS2 intensity fraction. Furthermore, this transfer substantially improved the comprehensive interpretation of the complete spectrum, resolving eight additional interpretable spectral peaks with structurally compatible annotations. More importantly, two of these additional peaks corresponded to the characteristic KYN peaks at *m/z* 120.044 and 174.055, ultimately providing clear substructure interpretations for all six signature spectral peaks previously reported for protonated KYN [46, 47]. To extend beyond isolated metabolite characteristics, we leveraged UltraMS to interrogate the broader KYN pathway within human carotid plaques (Fig. 5i, right panel). Multi-node annotation delivered by UltraMS revealed that KYN and its key downstream intermediates, anthranilic acid (AA) and kynurenic acid (KYNA), were elevated by 2.11-, 3.25-, and 1.80-fold in symptomatic relative to asymptomatic plaques. This tissue-resolved pathway view coherently unifies previously separate lines of clinical and molecular evidence (see detailed discussion in Supplementary Material Section S1.5). Here, operating directly on human plaque metabolomic spectra, UltraMS demonstrates its integrative capacity to efficiently resolve multiple intermediates and connect disconnected clues into a coherent pathway chain of evidence. Overall, UltraMS demonstrated its remarkable capacity to provide a chemically reasonable avenue towards interpreting molecular dissociation patterns and its potential to offer valuable insights for clinical research based on corresponding metabolic profiles.

### 2.6 A million-scale navigable mass spectral atlas illuminates metabolic microenvironments and uncovers microbial niches on human scalp via learned chemical geography

One prominent advantage of UltraMS is that it extends library-matching-based molecular identification beyond the comparison of raw spectral similarity to deeper identification driven by chemical semantic understanding. Particularly, spectra acquired from the same molecule under similar experimental conditions can exhibit substantial peak overlap and high raw spectrum cosine similarity. However, practical applications are more challenging, as variation in collision energy, experimental noise or even operational deviation causes one molecule to produce markedly different mass spectra through diverse molecule fragmentation patterns [32]. To address this challenge, we began by enhancing the spectral representation capacity of UltraMS by incorporating broadly available public spectrum-molecule pairs into contrastive learning (see detailed method in Supplementary Material Section S1.7), and evaluated molecule identification via spectrum-to-spectrum similarity ranking on 6,159 challenging spectrum queries for which neither precursor mass nor mechanically counting overlapping peaks can easily distinguish the correct molecule from the interfering candidates. UltraMS outperformed all comparison methods across all three complementary evaluation criteria. Its advantage was first evident in recognizing the correct molecular identity across markedly diverse spectra: UltraMS achieved a mean molecule identification AUROC of 0.764, compared with 0.662 for DreaMS and 0.484–0.533 for direct spectral cosine similarity matching and neural network encoding baselines (Fig. 6a). This improvement also extended beyond exact molecular identification to yielding recommendations with higher structural similarity: among the spectrum pairs in the top 10% of embedding similarity for each method, UltraMS captured remarkably more coherent chemical relationships, with a mean ECFP4 Tanimoto similarity [48] of 0.938, substantially higher than DreaMS at 0.75 (Fig. 6b). Notably, UltraMS embedding similarity further provided an effective readout of molecular structure, most pronounced among pairs with high embedding similarity, with a pooled Pearson *r* of 0.39, compared with 0.17 for DreaMS (Fig. 6c). Complementary evaluation on the remaining, less challenging subset of the independent test set further showed that UltraMS is particularly advantageous when raw spectral cosine similarity no longer offers a viable shortcut and molecular identity must instead be uncovered from the underlying chemical semantics (Supplementary Fig. 9; see detailed discussion of performance in Supplementary Material Section S1.6).

**Figure 6.**
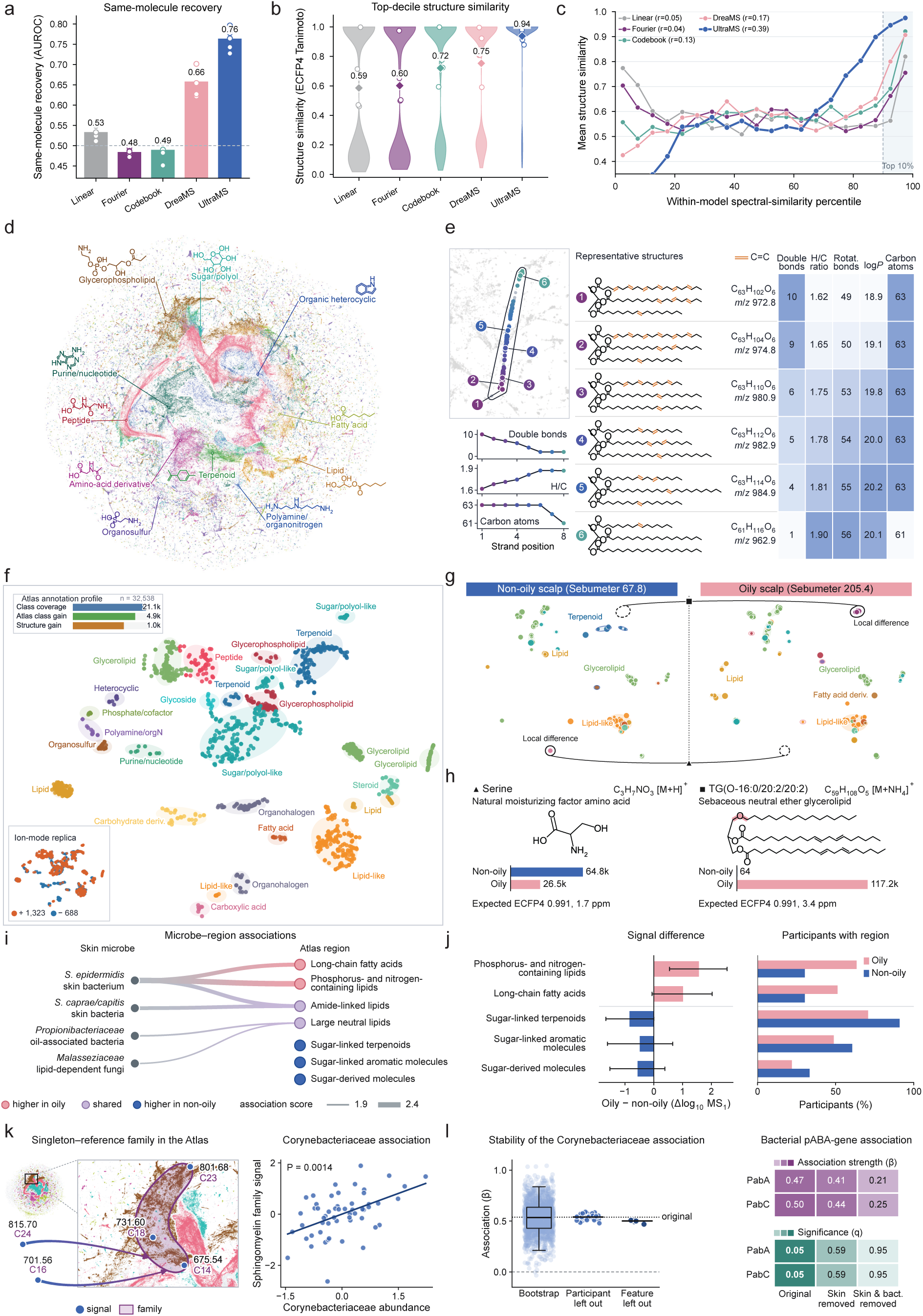
UltraAtlas establishes a multi-million-scale, chemically meaningful, navigable landscape and demonstrates its utility in profiling scalp phenotypic metabolic microenvironments and revealing microbial ecological niches across a human cohort. **a**, Recovery of molecular identity under substantial spectral divergence. Benchmark evaluation across 6,159 query spectra paired with same-molecule reference spectra exhibiting raw cosine similarity *<* 0.4, accompanied by hard-to-discriminate interfering spectra strictly matched for precursor *m/z*, ionization polarity, and raw spectral similarity. Bars indicate mean per-query area under the receiver operating characteristic curve (AUROC) in distinguishing authentic same-molecule pairs; open circles denote individual folds across five-fold molecule-disjoint cross-validation. **b**, Structural fidelity of candidate molecules identified through top-ranking spectral retrieval. Violin plots depict the distribution of molecular structural similarity (ECFP4 Tanimoto coefficient) between compounds corresponding to the top 10% highest-scoring retrieved spectral pairs across evaluated methods. Diamonds and open circles denote pooled means across all spectral pairs and means from individual validation folds, respectively. **c**, Alignment between spectral similarity and molecular structural resemblance. Curves depict mean ECFP4 Tanimoto similarity across 20 bins of spectral similarity, determined independently for each method. The shaded region highlights the top decile (top 10%) of spectral pairs. Values in the legend (*r*) indicate pooled Pearson correlation coefficients between spectral similarity and molecular structural similarity. **d**, Global chemical organization of UltraAtlas. Overview of UltraAtlas comprising approximately 4.20 million navigation nodes. Colors highlight 13 major chemical classes across class-annotated nodes, revealing distinct, chemically organized regions within the shared spectral “geometric map”. **e**, Continuous chemical semantic transitions along a local triglyceride-like trajectory. Numbered molecular structures (1–6) correspond to sequential positions along an O_6_ triglyceride-like strand. Heatmap and trajectory curves show progressive shifts in physicochemical properties along this path: estimated acyl unsaturation decreases alongside concurrent increases in the H/C ratio and number of rotatable bonds, while total carbon content remains largely conserved. **f**, Mapping and annotation expansion of scalp metabolic features via UltraAtlas. Distribution of 2,011 scalp molecule features mapped onto 28 representative class-resolved chemical regions; colors denote chemical classes, with the lower left inset showing ionization polarity distribution. Upper left inset: Annotation expansion across all 32,538 detected scalp molecule features. Class coverage reflects the integration of original and UltraAtlas annotations, whereas class and structure gains denote features for which UltraAtlas uniquely provided chemical class or structural assignments absent from original study annotations. **g**, Comparative scalp metabolic landscapes between distinct phenotypic samples in shared UltraAtlas coordinates. Scalp molecule features from representative non-oily and oily participants mapped onto the unified reference landscape. Distinct symbols highlight representative features selected for structural elucidation in **h**. **h**, Structural elucidation for the marked scalp features. Expected structural similarity (ECFP4 Tanimoto coefficient) is quantitatively calibrated from UltraMS embedding similarity. Bar plots display raw MS1 peak intensities of each feature across both participants(independent scales). **i**, Mapping of scalp microbe–metabolite associations across UltraAtlas-defined chemical regions. Edges denote associations reported across 55 participants with corresponding metabolomic and microbial profiles; line width reflects association strength. **j**, Phenotype-associated metabolic signatures statistics across UltraAtlas-resolved chemical class regions in the full scalp cohort (*n* = 41 oily and *n* = 33 non-oily participants). Left: difference in mean log_10_(MS1 intensity + 1) between oily and non-oily participants; error bars represent 95% bootstrap confidence intervals derived from participant resampling. Right: percentage of participants in each group with detected molecule features in the chemical class region. **k**, Five GNPS singleton molecule features mapped to an SM family in UltraAtlas, labelled by *m/z* and *N* -acyl chain length (left). The combined SM family signal was positively associated with *Corynebacteriaceae* abundance across 55 participants after covariate adjustment, including skin sebum and hydration (right; each point represents one participant). **l**, Stability of the SM–*Corynebacteriaceae* association under participant-level bootstrap resampling and leave-one-participant/feature-out analyses (left). Associations with the pABA-biosynthesis genes *pabA* and *pabC* before and after adjustment for skin sebum, hydration and *Corynebacteriaceae* abundance (right; top, association coefficients *β*; bottom, Benjamini–Hochberg-adjusted *q*).

This ability to recognize latent structural coherence underlying chemical organization from mass spectra motivated us to explore whether UltraMS could further map large numbers of spectra from unidentified molecules directly and efficiently into biochemically meaningful and well-defined chemical class clusters. UltraMS achieved a mean class-wise AUROC of 0.933 across the 10 RDKit [49] structural classes and 0.923 across the 20 ChEBI [50] biochemical classes with sufficient reference molecules (Supplementary Figs. 10–12; see detailed method in Supplementary Material Section S1.6). At an empirical FDR of no more than 5%, class-specific confidence thresholds were established for all 10 RDKit classes and 19 of the 20 ChEBI classes, defining 29 classes that UltraMS could reliably assign to spectra of previously unseen molecules. Grounded in this validated framework, we constructed a global mass-spectral map called UltraAtlas (Fig. 6d) using 163.7 million experimental tandem mass spectra from UltraMSdata with 1.2 million structurally annotated reference spectra from MSnLib, Spectraverse and MassSpecGym. These spectra formed approximately 4.2 million navigation nodes, making every spectrum in UltraMSdata rapidly reachable through its corresponding chemical neighborhood (see detailed method in Methods). The resulting map revealed a class-resolved chemical geography: 13 major chemical classes formed characteristic territories, exposing both distinct molecular families and broader chemical relationships within neighborhoods. To further evaluate the ability of UltraAtlas to provide meaningful navigation through chemical geography, we extracted one million spectra from the 201.2-million-spectrum GeMS-C collection [18], leaving 750,464 spectra as the external validation corpus after excluding query–Atlas pairs with a raw-spectrum cosine similarity of more than 0.95 (Supplementary Fig. 13). The mean expected ECFP4 Tanimoto similarity reached 0.972 across the 750,464 spectrum queries, compared with only 0.155 for the random matched pairs (difference, 0.817; 95% CI, 0.808–0.831; *P* = 1.0 10^−5^; Supplementary Fig. 14; see detailed method in Supplementary Material Section S1.6). Together, these results demonstrate that, even when raw spectral agreement no longer provides a direct cue for uncovering molecular identity, UltraAtlas is able to efficiently map massive numbers of diverse spectra from uncharacterized molecules to local regions with consistently high expected structural similarity within its learned chemical geometry. In tandem with this, UltraAtlas further resolved meaningful chemical variation within the distinctly shaped regions across its map. In one lipid-rich region, navigation nodes aligned along the principal direction of the local Atlas geometry while retaining a shared O_6_ TG-like composition (Fig. 6e). Along this direction, estimated acyl unsaturation declined smoothly from 10 to 1, accompanied by an increase in H/C ratio from 1.62 to 1.90 and in the number of rotatable bonds from 49 to 56, while oxygen stoichiometry remained fixed and carbon content was largely conserved, showing that its learned chemical geometry goes beyond identifying molecular relationships to reveal the direction and extent of chemical variation underlying mass spectra.

To establish the practical utility of UltraAtlas, we applied it to the independent MSV000088283 human scalp study [51], which yielded 32,538 non-redundant molecule features with associated tandem mass spectra. UltraAtlas first depicted the overall metabolic landscape by resolving these otherwise unorganized molecule features into spatially coherent regions spanning distinct chemical classes within its unified spectral coordinate system (Fig. 6f). In addition, the original GNPS/SIRIUS/CANOPUS analysis provided class-level information for 16,236 molecule features (49.9%), and UltraAtlas supplied supported class-level information for a further 4,863 molecule features that lacked an original label, delivering a substantial 30.0% increase in chemical-class annotation (Supplementary Fig. 15; see detailed method in Supplementary Material Section S1.6). Beyond overall dataset profiling, the shared chemical-geography projection of UltraAtlas enables direct, interpretable comparisons between individual samples, effectively highlighting phenotypic metabolic divergence. In particular, we observed that the UltraAtlas maps of representative non-oily and oily samples exhibited sharply contrasting local morphologies (Fig. 6g): the non-oily map specifically contained a discrete carboxylic-acid region, whereas the oily map contained a salient lipid region. In the representative non-oily sample, UltraAtlas delivered serine as its molecular recommendation with an expected ECFP4 Tanimoto similarity of 0.991, identifying a major amino acid in the skin’s natural moisturizing system that helps retain water [52]. Its unique presence on the non-oily Atlas map and substantially higher abundance than in the oily sample revealed a stronger natural-moisturizing-factor signal on the non-oily scalp, consistent with previous evidence that excess scalp oil dilutes metabolites from the outer skin relative to oil-poor scalp surfaces [51] (Fig. 6h). In contrast, the distinct region in the oily sample returned TG(O-16:0/20:2/20:2) as the UltraAtlas molecular recommendation, with an expected ECFP4 Tanimoto similarity of 0.991 and a substantially higher MS1 abundance than in the non-oily sample (Fig. 6g, h). This molecule belongs to a skin-enriched class of large neutral lipids released with natural skin oil [53, 54]. Together, these results show the capacity of UltraAtlas to convert a visible difference between individuals into a biologically grounded contrast between water-retaining chemistry and oil-associated lipid accumulation.

We subsequently probed how the visible metabolic patterns revealed by UltraAtlas could extend our interpretation of metabolic characteristics across the wider scalp ecosystem. For the 55 participants with both metabolic and microbial measurements, we leveraged UltraAtlas to identify the molecular regions in which the spectral features linked to diverse skin microbes in the original study [51] accumulated (Fig. 6i). We found that *Staphylococcus epidermidis*, a common bacterium on human skin, was linked to regions containing phosphorus- and nitrogen-containing lipids, long-chain fatty acids and amide-linked lipids, whereas *Propionibacteriaceae*, a group of bacteria commonly associated with oily skin, and *Malasseziaceae*, a family of skin fungi, were both linked to large neutral lipids found in skin oil. Quantification across all 74 participants using MS1 intensity and participant prevalence further demonstrated a consistent trend: phosphorus- and nitrogen-containing lipids and long-chain fatty acids were more abundant in oily participants, whereas sugar-linked terpenoids, sugar-linked aromatic molecules and sugar-derived molecules were more abundant in non-oily participants (Fig. 6j). Together, these findings demonstrate that the spatial patterns resolved by UltraAtlas carry genuine biological significance, faithfully reflecting the established ecological niches and nutritional requirements of the scalp microbiome (see detailed discussion in Supplementary Material Section S1.6). Moreover, five molecule features, originally represented as disconnected singletons in the GNPS molecular network, formed a compact sphingomyelin (SM) family cluster in UltraAtlas (Fig. 6k, left), and UltraAtlas suggested that they share a common SM backbone and differ primarily in their *N* -acyl chain lengths (C_14_–C_24_). Guided by this chemical family assignment, we next examined its ecological associations with the skin microbiome: the SM family signal was positively associated with *Corynebacteriaceae* relative abundance after covariate adjustment, including skin sebum and hydration (*β* = 0.54, *P* = 0.0014; Fig. 6k, right), consistent with prior evidence from porcine models, in which multiple SM species positively covaried with *Corynebacterium* species [55]. The family-level association remained robust when each participant or each of the five SM family molecule features was left out in turn, and under participant-level bootstrap resampling (Fig. 6l, left). To further explore the functional basis of this microbe–metabolite association, we profiled microbial functional potential using the paired scalp shotgun metagenomes (see detailed method in Methods and Supplementary Material Section S1.6): across 4,895 candidate functions, the strongest positive associations with the SM family signal converged on *pabA* and *pabC* (Fig. 6l, right), which encode key enzymes for the synthesis of *para*-aminobenzoic acid (pABA), an indispensable metabolite powering downstream nucleotide production essential for sustaining bacterial cell division and population expansion. Removing the effect of *Corynebacteriaceae* relative abundance weakened the associations between both genes and the SM family signal by approximately half. Collectively, these findings suggest an interdependent tripartite metabolic structure that bridges host human scalp sphingomyelins, *Corynebacteriaceae* abundance and microbial pABA biosynthesis, revealing a plausible model from otherwise unorganized and disconnected MS data, in which an SM-enriched microenvironment favours the growth of *Corynebacteriaceae*, whose proliferation relies on pABA-dependent pathways.

## 3 Discussion

Inspired by the recent success of foundation models in decoding domain-specific ’languages’ across the life sciences [13–15, 56], we investigated whether large-scale pre-training can capture the intrinsic molecular semantics encoded in tandem mass spectra, which is vastly abundant and largely unannotated, with substantially diverse fragmentation patterns and generally pervasive background noise, yet conceals an immense reservoir of chemical intelligence. Combining structurally grounded spectral encoding with a chemical-prior-aware decoding architecture, we present UltraMS, a large-scale foundation model with 200 million parameters pretrained on 161 million full-polarity mass spectra. Through this design, UltraMS reconciles spectral peak features with discrete structural semantics, transforming uncurated, multi-instrument archives into an informative and unified chemical latent space, establishing an adaptable foundation for navigating diverse chemical landscapes and supporting molecular exploration in a scalable and interpretable manner.

We demonstrate the benefits of pretraining on 161 million full-polarity spectra with comprehensive experiments in both open public benchmarks and realistic application study settings. The pretrained model showcases strong capabilities in unraveling molecular structural information and chemical properties to organize spectra into a chemically meaningful molecular landscape, presenting well-defined clustering patterns in accordance with key molecular attributes such as mass consistency, elemental composition, and molecular hydrophobicity. In addition, the representations learned by UltraMS exhibit excellent performance in capturing fragmentation principles, such as predicting general neutral losses and tracking essential atom transfers between specific spectral peaks. Furthermore, the pretrained model’s knowledge can be transferred to a wide range of downstream tasks through task-specific adaptation and benefit realistic applications by resolving ambiguous annotations and uncovering valuable biological insight in complex disease cohorts. In a variety of tasks such as elemental detection, molecular identification, molecular property profiling, spectral peak interpretation, and metabolic landscape characterization, the pretrained UltraMS model consistently outperforms existing baselines, including domain-specific deep learning models trained from scratch and pioneering pretrained foundation models. This demonstrates the potential of pretraining on large-scale mass spectra without demanding expensive and labor-intensive annotation, enabling accurate molecular exploration and the acquisition of meaningful biological insights. Leveraging UltraMS’s superior ability to depict molecular space across massive spectral archives, our developed application, UltraAtlas, constructs a navigable reference map spanning millions of spectra for guided chemical exploration. Its optimized search architecture not only enables millisecond-level spectral search and mapping, but also empowers the global profiling of metabolomic landscapes while prioritizing phenotype-discriminative molecular features, providing a transformative platform for large-scale chemical exploration in untargeted metabolomics.

Looking forward, several promising directions could further expand the horizons of mass spectrometry via large pretraining foundation models. First, we plan to scale pretraining to substantially larger spectral archives encompassing greater diversity in instrumentation, ionization mode, biological specimen types and application regimes. Additionally, leveraging the transferable latent representations learned by UltraMS also offers an exciting avenue for augmenting existing computational tools with chemically informative peak-level annotations by providing valuable instruction to guide rational structure assembly for tasks such as de novo molecular structure elucidation. Furthermore, a promising frontier lies at the intersection of domain-specific spectral foundation model and general large language models [57–59] (LLMs). Developing effective approaches to transform spectral embeddings into a chemically grounded representation space clears the path for LLMs to iteratively reason and gradually infer structural information over fragmentation events. This empowers researchers to harness advanced AI reasoning to systematically illuminate the vast unexplored “chemical dark matter” [6] and uncover novel metabolites across metabolic repositories. Ultimately, we envision this self-supervised pretraining paradigm becoming a foundation of mass spectrometry research to leverage generalizable chemical knowledge and meaningful biological insights from the exponential accumulation of mass spectra in global repositories for future discoveries.

## Acknowledgments

This research is supported by National Natural Science Foundation of China Project (No. 623B2086), TeleAI of China Telecom, Ant Group, CIPS-SMP research funding for Large Language Models and Tencent. We thank Wen-Feng Zeng (Westlake University) and Cheng Chang (National Center for Protein Sciences, Beijing) for their valuable suggestions.

## Author contributions

Yi.L. and Ju.X. conceived the study. Yi.L. designed the methodology, assembled the UltraMSdata pretraining corpus, and developed, implemented and pretrained UltraMS. Yi.L. designed and performed the benchmark, spectral peak interpretation, clinical plaque metabolomics and scalp metabolomics analyses, and constructed UltraAtlas. Yi.L. and Y.Z. prepared the figures. Yi.L. wrote the manuscript. Y.Z., P.L., Z.W., Ji.X. and Ju.X. helped refine the manuscript. Ju.X. supervised the project and acquired funding. All authors discussed the results and reviewed the manuscript.

## 4 Method

### 4.1 UltraMS model architecture

#### Architecture overview

UltraMS represents a tandem mass spectrum at two complementary levels: a spectrum-level embedding that summarizes the complete fragmentation pattern to represent the global molecule information and contextualized peak-level embeddings that retain fine-grained substructure information about individual spectral peaks. The model contains three major components (Fig. 1b–e). A Continuous Multi-resolution Peak Encoding (CMPE) module converts the *m/z* and intensity values in the continuous space into learnable representation. A transformer encoder [20] contextualizes the spectral peaks within the spectrum. Finally, a Sequentially Conditioned Fine-grained Reconstruction (SCFR) decoder reconstructs masked spectral peaks from other contextualized spectral representations, providing the self-supervised signal used to pretrain the model.

#### Embedding module

We represent a spectrum as a set of spectral peaks 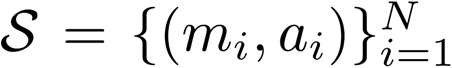, where *m_i_* is the measured *m/z* and *a_i_* is the intensity normalized to the maximum intensity in that spectrum. CMPE converts these continuous values into embeddings without assigning every possible value to an independent discrete token. CMPE uses four resolution levels indexed by *r* ∈ *{*0, 1, 2, 3}. At resolution level *r*, it defines an ordered set of spectral representation anchors

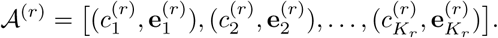

Each spectral representation anchor pairs a fixed reference position *c_k_*^(*r*)^ with a learnable embedding *e_k_*^(*r*)^; the embedding list [*e*_1_^(*r*)^, *e*_2_^(*r*)^,*…*] is therefore aligned with the position list [*c*_1_^(*r*)^, *c*_2_^(*r*)^, *…*]. Each resolution level maps *m_i_* to one coordinate *v_i_*^(*r*)^ and compares that coordinate with the *K_r_* spectral representation anchors in *A*^(^*^r^*^)^, where *K_r_* is selected separately for each resolution level. The first two levels use *m_i_* itself and place the spectral representation anchors along the full mass axis at approximately 10-Da and 1-Da intervals, respectively. The next two levels locate the same peak within its current 1-Da and 0.1-Da intervals. The four coordinates are therefore

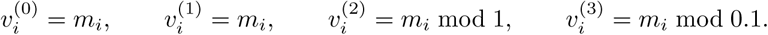

For example, a peak at *m/z* = 403.26 is presented to the four levels as

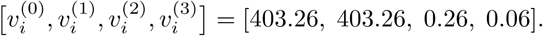

The first two coordinates are compared with spectral representation anchors spanning the full mass axis, giving the largest weights to those near 400 and 403, respectively. The latter two are compared with spectral representation anchors within [0, 1] and [0, 0.1], giving the largest weights to those near 0.26 and 0.06. Thus, the first two levels locate the peak on the full mass axis, whereas the latter two preserve its fractional positions within 1-Da and 0.1-Da intervals.

At each resolution level *r*, CMPE converts the coordinate *v_i_*^(*r*)^ into a resolution-level-specific embedding **z***_i_*^(*r*)^ by aggregating the learnable embeddings of the spectral representation anchors, with nearer spectral representation anchors receiving larger weights:

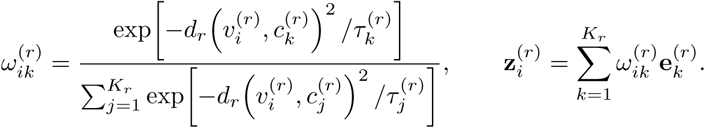

For the two levels defined over the full mass axis, distance is the absolute difference *d_r_*(*v, c*) = |*v*−*c*|. For the levels defined within the 1-Da and 0.1-Da intervals, the shorter circular distance is used: *d_r_*(*v, c*) = min(|*v* − *c*|*, P_r_* − *v c*), where *P_r_* is 1 or 0.1 Da, respectively. The learnable *τ_k_*^(*r*)^ is the temperature parameter for spectral representation anchor *k*: a lower temperature concentrates the weight on the nearest spectral representation anchors, whereas a higher temperature distributes it across more spectral representation anchors. Concatenating the four interpolated representations gives

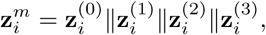

where || denotes concatenation. Normalized intensity is encoded by a separate set of spectral representation anchors positioned over [0, 1], using the same interpolation to obtain **z***_i_^a^*. The final embedding of spectral peak *i*, and hence the vector passed to the transformer, is

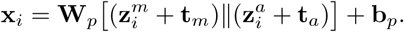

Here, **t***_m_* and **t***_a_* are learnable type embeddings added to the *m/z* and intensity representations, respectively, before the two representations are concatenated. The matrix **W***_p_* then maps the concatenated representation into the transformer space, and **b***_p_* is the learnable bias vector. The precursor ion is embedded through the same mapping and is distinguished from spectral peaks by a learned precursor-type embedding.

#### Encoder

The spectral peaks are ordered by increasing *m/z*. UltraMS then places a learned [CLS] embedding and the precursor-ion embedding before the CMPE embeddings of the spectral peaks. After adding a learned positional embedding **p***_j_* at each position, the encoder input is

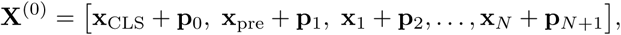

where **x**_CLS_ is a learned embedding, **x**_pre_ is the precursor-ion embedding and **x***_i_* is the CMPE embedding of spectral peak *i*. The [CLS] element does not correspond to a measured ion; it serves as a learned position for collecting information from the complete spectrum.

The fundamental task of tandem mass spectrometry is to decode the latent molecular information from complex dissociation phenomena. However, critical structural determinants are implicitly entangled within subtle, highly coupled fragmentation events and peak co-occurrence patterns across the spectral background. Multihead self-attention of transformer blocks directly addresses this chemical complexity by simultaneously modeling relationships among individual spectral peaks [20], enabling the encoder to learn the latent chemical language reflected in diverse fragmentation patterns and decipher the structural information collectively encoded across the complete spectrum:

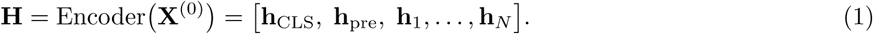

The complete input sequence **X**^(0)^ is processed by a stack of transformer encoder layers, denoted by Encoder(). Each hidden state **h***_i_* corresponds to one spectral peak. Its CMPE input initially describes the *m/z* and intensity of particular peak. After self-attention, **h***_i_* also incorporates information from the precursor ion and all other peaks in the spectrum. The representation of a peak therefore depends on how it fits into the complete fragmentation pattern, allowing the same or similar peak to carry different structural meanings in different spectral contexts. The precursor position is updated in the same way to produce **h**_pre_. At the spectrum level, the [CLS] position gathers information from the precursor ion and all fragment peaks, and its output **h**_CLS_ is used as the spectrum-level embedding. During pretraining, the output at each masked peak position is passed to the SCFR decoder, which reconstructs the missing peak’s *m/z* and intensity from the precursor ion and the remaining fragmentation pattern.

#### Decoder

The SCFR decoder reconstructs the *m/z* and normalized intensity of each masked peak from its contextualized representation **h***_i_*. It predicts mass at three successive resolutions: 10 Da, 1 Da and 0.1 Da. Each finer level uses both **h***_i_* and the mass estimated at the preceding level.

The first level predicts probabilities over 200 10-Da intervals spanning *m/z* = 0–2000 and calculates a mass estimate from their centres:

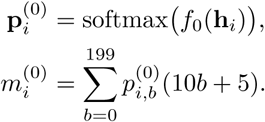

Here, *b* indexes the intervals, *p_i,b_*^(0)^ is the predicted probability of interval *b*, and 10*b* + 5 is its centre. For example, probabilities of 0.7 and 0.3 for the 400–410 and 410–420 Da intervals, with zero probability elsewhere, give *m_i_*^(0)^ = 0.7 × 405 + 0.3 × 415 = 408 Da.

The decoder then represents *m_i_*^(0)^ using the same *m/z* component of CMPE defined above. We denote this mapping by **z***^m^*(*m*), with **z***^m^*(*m_i_*) = **z***_i_^m^*. The second level combines this mass representation with **h***_i_* and predicts ten 1-Da positions:

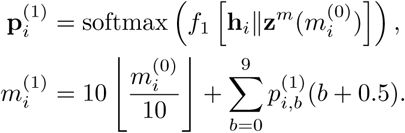

The first term gives the start of the 10-Da interval containing *m_i_*^(0)^. The second term calculates an average position estimate using the predicted probabilities to weight the centres of the ten 1-Da intervals, located at 0.5, 1.5*, …,* 9.5 Da from the start of the 10-Da interval. Adding these two terms gives the refined mass estimate *m_i_*^(1)^.

The third level uses the refined mass *m_i_*^(1)^ in the same way to predict ten positions at 0.1-Da resolution:

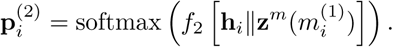

The function *f*_0_ is a learned linear transformation. Both *f*_1_ and *f*_2_ contain two learned linear transformations separated by a GELU activation, with separate parameters for each level.

The ten entries of p*_i_*^(2)^ give the probabilities of positions 0, 0.1*, …,* 0.9 Da within a 1-Da interval. To obtain the final mass prediction, the decoder selects the position with the highest probability at each of the three levels:

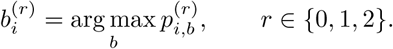

Here, *b_i_*^(0)^ identifies the predicted 10-Da interval, *b_i_*^(1)^ gives the integer-Da position within it, and *b_i_*^(2)^ gives the position in 0.1-Da increments within that 1-Da interval. The reconstructed mass is therefore

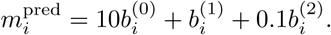

For example, predicted class indices of 40, 3 and 2 give *m_i_*^pred^ = 400 + 3 + 0.2 = 403.2 Da.

Normalized intensity is reconstructed by choosing among ten possible values: 0, 0.1*, …,* 0.9. A learned linear transformation *f_a_*, followed by a softmax, maps **h***_i_* to the probabilities of these ten values:

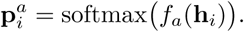

The decoder selects the intensity value with the highest predicted probability:

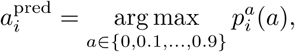

Where *p_i_^a^*(*a*) is the predicted probability of intensity value *a*.

During training, each finer level is given the correct 10-Da or 1-Da interval of the masked peak with probability 0.5 to stabilize training and reduce error propagation across levels. In the remaining cases, each finer level is conditioned on the mass estimated by the preceding level, maintaining the same coarse-to-fine prediction protocol as during inference. Gradients are not propagated through these mass values or their CMPE representations. During prediction, both transitions use the estimated masses. The three mass distributions and the intensity distribution are supervised using the measured values of the masked peak.

#### Pretraining task

UltraMS adopts self-supervised masked peak reconstruction (MPR) as the core training objective. Specifically, the model continually recovers missing spectral peaks by leveraging the precursor ion and remaining spectral peak context, directly capturing latent structural semantics and underlying chemical dissociation logic across large-scale unannotated spectra.

For each masked peak *i*, SCFR produces three mass distributions, **p***_i_*^(0)^, **p***_i_*^(1)^ and **p***_i_*^(2)^, together with the intensity distribution **p***_i_^a^*. Training increases the probabilities assigned to the measured mass and intensity of that peak.

For mass reconstruction, we denote the correct class at each resolution by *b_i,true_*^(*r*)^, corresponding to the predicted class *b_i_*^(*r*)^ defined in SCFR.

The mass reconstruction loss *l_i_^m^* adds the negative log-probabilities assigned to these three correct classes:

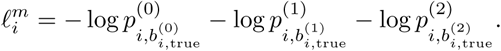

Each term therefore evaluates one SCFR prediction: the 10-Da interval, the 1-Da position or the 0.1-Da position. Assigning a higher probability to the correct class reduces its contribution to the loss.

For intensity reconstruction, the spectral peak’s normalized intensity *a_i_* is rounded to the nearest 0.1 and restricted to the ten available values 0, 0.1*, …,* 0.9. We denote the resulting target value by *a_i_*^target^. Using the intensity probability *p_i_^a^*(a) defined in SCFR, the intensity loss is

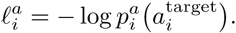

We apply difficulty weighting to the mass reconstruction loss. We define the difficulty score *d_i_* using the normalized entropy of SCFR’s first-level distribution p*_i_*^(0)^. This is motivated by the coarse-to-fine design of SCFR, which first locates the masked peak within a broad mass interval, capturing the structural information about the fragment’s core chemical scaffold, providing a readily available overall estimate of reconstruction difficulty.

A prediction concentrated on one specific *m/z* interval has an uncertainty score near zero, whereas probabilities spread evenly across all *m/z* intervals give an uncertainty score near one. The uncertainty score *d_i_* and the corresponding peak weight *w_i_* are

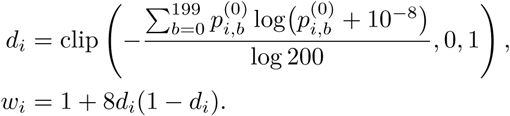

The small constant 10^−8^ prevents taking the logarithm of zero, and clipping restricts the difficulty score to [0, 1]. The weight is largest at intermediate difficulty: it reaches 3 when *d_i_* = 0.5 and approaches 1 when *d_i_* approaches either zero or one. The same weight is applied to all three mass predictions for that peak.

Ultimately, let denote the set of masked peaks in a training batch, and let be their number. The masked peak reconstruction objective averages the weighted mass loss and the intensity loss over these peaks:

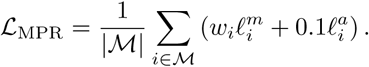

Here, *w_i_l_i_^m^* is the difficulty-weighted loss for the three SCFR mass predictions, and 0.1*ℓ^a^*is the intensity loss with a fixed coefficient of 0.1.

#### Auxiliary training tasks

Following MPR pretraining, UltraMS incorporates retention time and ion mode as auxiliary supervision to enhance spectrum-level representations with chromatographic behaviour and ionization mode awareness. Both tasks use the embedding **h**_CLS_*_,s_* of the complete, unmasked spectrum *s*, while MPR continues on a masked version of the same spectrum.

For retention-time prediction, a learned function *f*_RT_ maps **h**_CLS_*_,s_* to a predicted retention time normalized by 600. The measured retention time *t_s_*, expressed in seconds, is divided by the same factor for training. The prediction loss is:

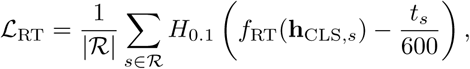

where *R* is the set of spectra with eligible retention-time labels in the minibatch. *H*_0_._1_ denotes the Huber loss, which applies a quadratic penalty to small errors and a linear penalty beyond a normalized error of 0.1, corresponding to 60 seconds.

For ion-mode prediction, a separate learned function followed by a sigmoid gives the positive-ion probability *p_s_*^ion^. Let *y_s_* = 1 denote positive ion mode and *y_s_* = 0 denote negative ion mode. The loss over the labelled spectra *I* is

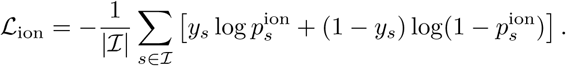

Training first incorporates the auxiliary task of retention-time prediction for five epochs, followed by eleven epochs of joint training with MPR and the auxiliary tasks of retention-time and ionization-polarity prediction. The joint training set contains equal numbers of positive- and negative-ion spectra. The two objectives are

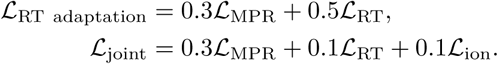

During joint training, we apply self-consistent retention-time supervision: a spectrum contributes gradients to *L*_RT_ only when its recorded retention time is in (0, 1500] seconds and its absolute prediction error is below 60 seconds. Otherwise, it contributes no retention-time gradient. This design prioritizes learnable retention-time relationships while reducing the influence of potentially misleading supervision arising from large systematic deviations caused by changes in instrument state or variations in experimental procedures.

4.2 Downstream methods

#### Element prediction and neutral-loss detection

We predict the numbers of S, Cl, F and Br atoms in the parent molecule and use the attention weights from each prediction to identify element-associated spectral peaks. For each element *E ∈* { S, Cl, F, Br }, a learned query assigns attention weights *α_i,E_* to the contextualized spectral peak-level embeddings **h***_i_* using scaled dot-product attention. A learned linear projection **W***_v_*, shared across the four elements, transforms the spectral peak embeddings before they are combined using these weights. The combined representation is concatenated with the spectrum-level embedding **h**_CLS_ and passed to an element-specific regression network *f_E_*. The predicted number of atoms of element *E*, denoted by *n_E_*^pred^, is

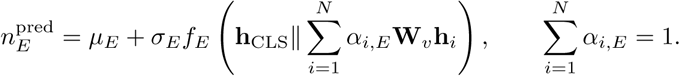

Here, *i* indexes the *N* spectral peaks, and *α_i,E_* is the attention weight of spectral peak *i* for element *E*. The network *f_E_* predicts a standardized atom count; the training-set mean *µ_E_* and standard deviation *σ_E_* of the atom counts for element *E* convert this output back to the original scale. Each *f_E_* contains two fully connected layers with a 2,048-dimensional input, a 1,024-dimensional hidden layer and a scalar output. Layer normalization and dropout with a rate of 0.1 precede the first fully connected layer; GELU activation and dropout with the same rate are applied between the two layers. The four elements have separate queries and regression networks but share the attention projections. These components are trained using mean squared error against standardized atom counts derived from the parent molecular formula.

We then rank spectral peaks by *α_i,E_* and assess their chemical relevance using spectral peak substructures annotated by MAGMa at an *m/z* tolerance of 10 ppm. We label a spectral peak as containing element *E* if at least one of its annotated substructures contains that element.

For neutral-loss detection, we predict whether a spectrum contains spectral peaks consistent with the loss of H_2_O, NH_3_, CO_2_, HF, HCl, CO or CH_3_ from the precursor ion. For each neutral loss, we subtract its theoretical mass from the precursor-ion *m/z* to obtain the expected spectral peak *m/z*. If the spectrum contains a spectral peak within 0.02 Da of this value, the training label is 1; otherwise, it is 0. A separate attention pooling module combines the spectral peak-level embeddings **h***_i_* into a summary of the spectral peaks. This summary is concatenated with the spectrum-level embedding **h**_CLS_ and passed to a classifier *f*_NL_. The classifier contains two fully connected layers with a 2,048-dimensional input, a 1,024-dimensional hidden layer and seven outputs, one for each neutral loss. Layer normalization and dropout with a rate of 0.1 precede the first fully connected layer; GELU activation and dropout with the same rate are applied between the two layers. Each output is passed through a sigmoid function to give the probability of the corresponding neutral loss. The attention pooling module and classifier are trained using binary cross-entropy, allowing multiple neutral losses to be detected in the same spectrum.

#### Visualization of spectrum-level embeddings

We visualize spectra using the spectrum-level embeddings obtained by applying the embedding projection learned during spectrum-to-spectrum contrastive training to the encoder output **h**_CLS_. We calculate cosine distances between these embeddings and use uniform manifold approximation and projection (UMAP) to map them into two dimensions while preserving local neighbourhood relationships. The resulting coordinates determine the positions of the spectra, while molecular structures and measured properties are used to colour the points.

For spectra with known molecular structures, we calculate Morgan fingerprints and apply Butina clustering [60] using Tanimoto distance to define molecular groups. Spectra are coloured according to the group of their parent molecule, allowing us to examine whether spectra from structurally related molecules cluster together in the UMAP visualization.

For realistic experimental samples, spectra are linked to their corresponding LC–MS molecule features and coloured by properties such as precursor-ion *m/z*, retention time, ion mode and abundance measured by MS1 intensity.

We divide each molecule feature’s intensity by the total molecule feature intensity within the same analytical run and apply a square-root transformation to obtain a normalized intensity. Experimental samples are grouped by their phenotype. To compare the abundance of each molecule feature between two phenotype groups, we calculate the difference between that molecule feature’s mean normalized intensities in the two groups. We then divide this intensity difference by the sum of the same molecule feature’s mean normalized intensities in the two groups to obtain its abundance contrast and reflect it through the colors of the corresponding spectral points in the UMAP visualization.

#### Visualizing fragmentation in the spectral peak-level embedding space

Alongside the spectrum-level embedding **h**_CLS_, we use learned spectral peak-level embeddings **h***_i_* to connect individual spectral peaks with the chemical properties of their corresponding substructures. UltraMS processes each complete spectrum and produces **h***_i_* at spectral peak *i* by incorporating information from the precursor ion and the other spectral peaks.

We begin by associating these embeddings with spectral peak substructures annotated by MAGMa. Given a spectrum and a candidate parent molecule, MAGMa simulates fragmentation of the candidate molecule to explain the observed spectral peaks. The resulting substructures are matched to spectral peaks according to their theoretical and measured *m/z*, providing substructure labels that describe how the candidate molecule can give rise to the observed spectrum. These labels provide the elemental composition and chemical structure information for each annotated spectral peak. We visualize the embeddings using UMAP with cosine distance and colour the spectral peak points by *m/z*, substructure composition, heavy-atom and heteroatom counts, and parent-molecule lipophilicity to display how these chemical properties are organized in the embedding space.

#### Spectral peak annotation transfer via structure-aware embeddings

We extend substructure annotations from reference spectra to query spectral peaks through *m/z* matching and spectral peak embedding similarity. UltraMS processes both query and reference spectra to obtain their spectral peak-level embeddings **h***_i_* and **h***_j_*. Specifically, for query spectral peak *i* and reference spectral peak *j*, we calculate their embedding similarity *s_ij_*

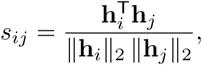

where **h***_i_*and **h***_j_*are the UltraMS spectral peak-level embeddings of the query and reference spectral peaks, respectively, and ∥ · ∥_2_ denotes the Euclidean norm.

For multiple spectra acquired for the same query molecule feature, we group corresponding spectral peaks by *m/z* and use their median *m/z* as the target value *m_i_*^target^ for query spectral peak *i*. We search reference spectra acquired in the same ion mode. Among reference spectral peaks with substructure labels annotated by MAGMa, we select those within 10 ppm of the target value:

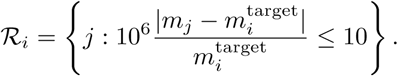

Here, *m_j_* is the measured *m/z* of reference spectral peak *j*, and *_i_* contains the indices of the labeled reference spectral peaks matching the target *m/z*. We rank these reference spectral peaks by their embedding similarity *s_ij_* to query spectral peak *i*.

We transfer spectral peak substructures annotated by MAGMa from the high-ranking reference spectral peaks to the corresponding query spectral peaks. This assigns chemical meaning to query spectral peaks through matching *m/z*, similar UltraMS spectral peak-level embeddings and existing substructure annotations. Additionally, we compare the transferred substructures with candidate parent molecules and identify substructures present in some candidates but absent from others. These substructure clues that come from transferred annotations of spectral peaks provide assistance in distinguishing the candidate molecules based on the chemical information in the observed spectrum.

#### Spectrum-to-molecule identification

We identify candidate parent molecules by comparing spectrum-level embeddings from UltraMS with molecular embeddings generated from candidate molecule structures. For spectrum *s*, a learned embedding projection *f*_mol_ maps **h**_CLS_*_,s_* into the molecular embedding space of ChemBERTa. The embedding projection contains two fully connected layers separated by GELU activation, layer normalization and dropout, together with a linear skip connection whose output is added to that of the two-layer branch. Candidate molecule structures are represented by the [CLS] output of the fixed ChemBERTa-100M-MLM model applied to their canonical SMILES. ChemBERTa [25] is trained on 100 million molecules from ZINC20 [61] and efficiently represents molecules as embeddings with abundant chemical information. We denote the projected spectrum embedding and the molecular embedding by **u***_s_* and **v***_c_*, respectively. Their cosine similarity gives the score for candidate molecule *c*:

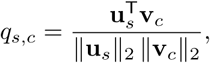

where ∥ · ∥_2_ denotes the Euclidean norm.

For a training batch containing *B* spectra, let *c*_1_*, …, c_B_* denote their known parent molecules, with *c_s_* being the parent molecule of spectrum *s*. The training objective is

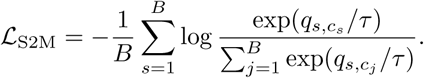

For each spectrum *s*, the numerator is the exponentiated similarity score for its correct parent molecule *c_s_*, while the denominator sums these scores over all *B* molecules in the batch. The temperature *τ* controls the distribution of probabilities assigned to candidate molecules: lower values make the distribution more concentrated, whereas higher values make it more uniform.

After training, when we perform molecular identification, for each query spectrum, we calculate *q_s,c_*for the candidate molecule structures in its predefined candidate set, constructed according to precursor-ion *m/z* using a reference database such as PubChem, and rank them in decreasing order to select the highest-ranked candidate molecule as the identification result.

#### Substructure-level discrimination of structural isomers

For each MSnLib test query, the competing isomer was selected as the highest-ranked incorrect candidate of the baseline model with the same molecular formula but different atomic connectivity. A transferred substructure annotation was considered to support the ground-truth structure when the query and reference spectral peaks agreed in *m/z* within 10 ppm and the substructure could be mapped onto that molecule—preserving atom types, connectivity, and bond types—but not onto the competing isomer.

#### Consensus molecular identification from spectra of the same molecule

Spectra acquired from the same molecule at different collision energies or on different instruments reveal different fragmentation patterns, providing substructure evidence for molecular identification. To integrate this evidence, we combine the candidate molecule rankings obtained from these spectra to identify the parent molecule.

For each spectrum, we apply the spectrum-to-molecule identification method to calculate the similarity score between the spectrum and each candidate molecule. We then select the 50 candidate molecules with the highest similarity scores, or all candidate molecules if fewer than 50 are available. Each spectrum contributes one vote to its highest-ranked candidate molecule, and the candidate molecule receiving the most votes is selected as the identification result.

When multiple candidate molecules receive the same highest number of votes, we use their ranks across the individual spectra to determine the final identification. For each of these candidate molecules, we calculate

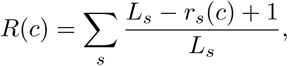

where *c* is a candidate molecule, *r_s_*(*c*) is its rank for spectrum *s*, and *L_s_* is the number of candidate molecules selected from that spectrum. Among the candidate molecules with the same highest vote count, we select the candidate molecule with the highest *R*(*c*) as the identification result.

#### Molecular identification through spectral clustering

For an input spectrum whose molecular structure is to be determined, we first compare it with spectra that have the same adduct and matching precursor-ion *m/z* and retention time. Raw spectral cosine similarity is used for complete-linkage clustering, which places spectra in the same group only when every pair has a cosine similarity over a special threshold. The threshold value is selected using the validation set and then fixed for test evaluation. After the input spectrum is grouped with other spectra most likely to originate from the same molecule, UltraMS uses the chemical information captured from each spectrum to rank the candidate molecules. Specifically, we apply the spectrum-to-molecule identification method described at “Consensus molecular identification from spectra of the same molecule.” to determine the final molecule identification result.

#### Molecular identification through fingerprint prediction

We predict molecular fingerprints from UltraMS representations and compare them with candidate fingerprints to identify the parent molecule. We train two separate fingerprint predictors using two types of input. The first uses the spectrum-level embedding obtained by applying the embedding projection to **h**_CLS_. The second concatenates this spectrum-level embedding with two readout of the spectral peak-level embeddings **h***_i_*: their mean and their intensity-weighted mean. Each predictor is a three-layer fully connected network with GELU activations and sigmoid outputs that give the probability of each fingerprint bit being active. The spectrum representations remain fixed, and the predictors are trained to predict the parent molecule’s radius-two Morgan fingerprint using weighted binary cross-entropy.

After training, we identify the parent molecule from the candidate set by comparing the predicted fingerprint with each candidate’s binary fingerprint. For each candidate molecule, we calculate the soft Tanimoto similarity *q* between its binary fingerprint and the predicted fingerprint:

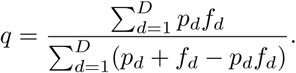

Here, *D* is the number of fingerprint bits, *p_d_* is the predicted probability that bit *d* is active, and *f_d_* is the corresponding binary value calculated from the candidate molecule. A higher *q* indicates greater agreement between the predicted and candidate fingerprints and serves as the criterion for selecting the final molecule as the identification result by combining the candidate ranking results obtained from the spectrum-level embedding predictor and from the predictor using the spectrum-level embedding concatenated with the two spectral peak-level readouts.

#### Spectrum-to-spectrum molecule identification

We identify the parent molecule of a query spectrum by comparing its UltraMS spectrum-level embedding with those of reference spectra with known molecular structures. For query spectrum *s*, the embedding projection maps its encoder output **h**_CLS_*_,s_* through layer normalization and two fully connected layers separated by a GELU activation to obtain **e***_s_*. Applying the same embedding projection to the encoder output **h**_CLS_*_,t_* of reference spectrum *t* gives **e***_t_*. We train these embeddings to bring spectra from the same molecule closer together using contrastive learning. When each molecule contributes two spectra, we use an InfoNCE loss [62] averaged over both comparison directions. When each molecule contributes multiple spectra, we use a supervised contrastive loss that treats all other spectra from the same molecule in the batch as positive examples and spectra from different molecules as negative examples.

After training, we apply the same model and embedding projection to query and reference spectra. For query spectrum *s* and reference spectrum *t*, we calculate their cosine similarity as

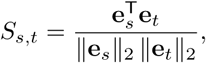

where **e***_s_* is the spectrum-level embedding of query spectrum *s* after the embedding projection, **e***_t_* is the spectrum-level embedding of reference spectrum *t* after the same embedding projection, and ∥ · ∥_2_ denotes the Euclidean norm. We rank reference spectra in decreasing order of *S_s,t_* and select the known parent molecule of the highest-ranked reference spectrum as the molecular identification result. We evaluate protonated and deprotonated ions separately, using reference spectra with the same adduct as the query spectrum.

#### Empirical FDR for spectrum-to-spectrum molecular identification

We use the empirical false-discovery rate (FDR) to estimate the proportion of incorrect molecular identifications accepted at a given spectral embedding similarity threshold. For a query with a known molecular structure, the identification is correct when the query and its highest-ranked reference spectrum have the same canonical SMILES. We accept the identification when the embedding similarity between these two spectra is at least *θ*, and calculate the empirical FDR and identification recall as

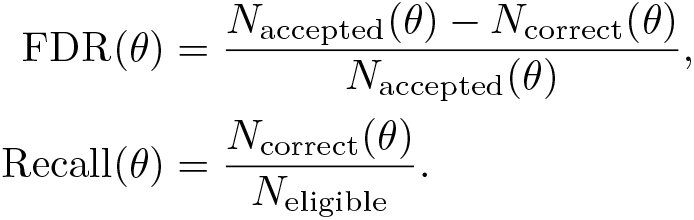

Here, *θ* is the embedding similarity threshold, *N*_accepted_(*θ*) is the number of accepted identifications, and *N*_correct_(*θ*) is the number of correct identifications among them. *N*_eligible_ is the number of queries whose parent molecule has at least one reference spectrum in the library. Otherwise, an accepted identification is counted as incorrect if the query has no same-molecule reference in the library.

#### Aligning spectral similarity with molecular structural similarity

We use spectra with known parent molecules to learn how UltraMS spectrum-level embedding similarity relates to molecular structural similarity. We represent each parent molecule by a 2,048-bit Morgan fingerprint with radius two (ECFP4) and calculate structural similarity as the Tanimoto similarity between two fingerprints: the number of shared active bits divided by the number of bits active in either fingerprint. For each reference spectrum used as a query, we exclude that spectrum from the reference library and select the remaining reference spectrum with the highest embedding similarity in the same ion mode.

For each query spectrum, we denote its embedding similarity to the highest-ranked reference spectrum by *S* and calculate the ECFP4 Tanimoto similarity between their parent molecules. We use isotonic regression to fit a non-decreasing function *g* that estimates molecular structural similarity from embedding similarity:

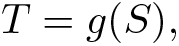

where *T* is the expected ECFP4 Tanimoto similarity between the query and reference parent molecules. We fit *g* by minimizing the weighted squared differences between the estimated and known Tanimoto similarities. The non-decreasing function aims to map higher embedding similarity to equal or higher expected structural similarity.

We evaluate this mapping by five-fold cross-validation, keeping all query spectra from the same molecule in the same fold. We fit *g* on four folds and evaluate it on the remaining fold using the mean absolute difference between estimated and known Tanimoto similarities. After validation, we fit *g* using all reference queries and apply it to external query spectra in independent test dataset. For each external query spectrum, the embedding similarity to its highest-ranked reference spectrum is converted into an expected ECFP4 Tanimoto similarity between their parent molecules.

4.3 Construction and application of UltraAtlas

#### Identify molecular classes directly from spectra

We predict molecular classes from UltraMS spectrum-level embeddings to provide chemical information for spectra whose parent structures are unknown. To describe both broad structural features and biochemical categories, we adopt two molecular-class systems using reference molecules from MSnLib, Spectraverse and MassSpecGym. The first comprises ten structural classes defined using RDKit rules for functional groups, elemental composition and ring structures. The second comprises 24 biochemical classes defined by ChEBI parent terms. Detailed ChEBI annotations are traced through their ancestor terms and assigned to these classes according to a predefined priority order. To avoid learning class patterns from only a few examples and support generalization to unseen molecules, we require each class to contain at least 100 molecules and 500 spectra. These requirements give ten RDKit classes and 21 ChEBI classes. We further exclude one ChEBI class with fewer than five validation molecules, leaving 20 ChEBI classes for molecule-disjoint evaluation.

For spectrum *s*, we use the spectrum-level embedding **e***_s_* obtained by applying the embedding projection learned during spectrum-to-spectrum contrastive training to the UltraMS encoder output **h**_CLS_*_,s_*. For each reference molecule, we average the normalized embeddings of its spectra and further leverage the resulting mean to obtain its molecular representation. We train a separate classifier for each class system, using layer normalization and two fully connected layers separated by GELU activation and dropout. Softmax converts the outputs into class probabilities, and the class with the highest probability is the predicted class within that system. To give less frequent classes greater weight during training, the loss for each molecule is weighted in proportion to 1*/*√*N*, where *N* is the number of training molecules in its class.

We evaluate class prediction using five-fold molecule-disjoint cross-validation. In each fold, the classifier is trained on four folds, with an internal validation subset used for model selection, and evaluated on the remaining fold. We pool the held-out predictions and select a probability threshold for each predicted class that accepts the largest number of class assignments while keeping the empirical FDR at or below 5%. A molecule’s class assignment is accepted when the probability of its predicted class is at least the threshold for that class.

#### Constructing UltraAtlas

We construct UltraAtlas by combining spectra from UltraMSdata with structure-annotated reference spectra from MSnLib, Spectraverse and MassSpecGym. To establish the initial map, we select three million spectra from UltraMSdata through stratified sampling across precursor-ion *m/z*, ion polarity, charge, collision energy and acquisition metadata. These three million spectra are combined with approximately 1.2 million reference spectra with an annotated parent molecular structure from the three databases to form the initial navigation nodes. We obtain their UltraMS spectrum-level embeddings **e***_s_* and apply principal component analysis followed by UMAP to jointly represent these nodes in two dimensions, organizing the chemical relationships captured by their embeddings into a shared chemically informative “geographical map”. Additionally, chemical-class labels derived from known reference structures and UltraMS embedding-based class predictions are used to colour the resulting map.

We then assign the spectra in UltraMSdata to navigation nodes according to their embedding similarities. This enables each node to provide access to a collection of related spectra. Moreover, to increase the spectral diversity represented by the navigation nodes, we expand the navigation nodes over several rounds. In each round, we take the spectrum least similar to each navigation node from its assigned spectra and make it a new navigation node, ultimately yielding approximately ten million navigation nodes with nearly 153 million additional spectra assigned to and distributed around them.

For each new navigation node, we select eight original nodes with the highest embedding similarities from the local map region of its original node. Its two-dimensional position in UltraAtlas is calculated as

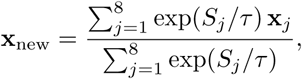

where **x***_j_* is the position of the *j*-th original node, *S_j_* is its embedding cosine similarity to the new node, and *τ* controls the weighting. More similar nodes therefore contribute more strongly to the position of the new node. For an incoming query spectrum, UltraAtlas first identifies relevant regions through the navigation nodes and then compares the query with the individual spectra contained in those regions. These spectra are ranked by embedding similarity to provide the final Atlas results.

#### Profiling sample chemistry and interpreting local differences using UltraAtlas

We map experimental samples into UltraAtlas by encoding their tandem mass spectra with the UltraMS model and embedding projection. For each query spectrum from an experimental sample provided as an MGF or mzML file, we identify its highest-ranked UltraAtlas result and place the molecule feature associated with this query spectrum at that result’s position in UltraAtlas. Chemical-class annotations are transferred from these results to the query spectra and further used to colour the molecule features. To offer an overall metabolic profile of the input sample, UltraAtlas then identifies and displays class-enriched regions in the resulting map to demonstrate the chemical organization of the sample.

To compare metabolic differences across experimental samples, we adopt a unified UltraAtlas coordinate system, allowing direct comparison of corresponding chemical regions across sample maps. Point sizes are scaled by the log-transformed MS1 intensities of molecule features to show local variation in molecule feature abundance.

### 4.4 Real-world biological and clinical metabolic datasets

MTBLS12303 contains LC–MS data from carotid artery plaques removed during surgery from 72 patients: 30 had recently experienced a stroke or transient ischaemic attack, and 42 had not experienced either event recently. The plaques were analysed using C18 and HILIC chromatography to characterize their chemical composition. The dataset combines measurements from the plaques themselves with patient symptom records and clinical measurements, including blood markers of inflammation and cholesterol levels. These data support comparisons of plaque chemistry between patients with and without recent symptoms and assessment of its relationship to their clinical characteristics. We restricted this analysis to molecular features within the upper decile of MS1 intensity and retained model search results only when their similarity scores exceeded the benchmark-derived threshold corresponding to an empirical FDR of less than 5%, which was established independently in Fig. 3. This filtering reduced the 13,116 initial features to 1,785 qualified features and ultimately identified 506 high-confidence molecular features with structural annotations.

MSV000088283 contains LC–MS/MS data from scalp-surface samples collected from 74 healthy volunteers: 33 with non-oily scalp and 41 with oily scalp. Each sample was analysed using C18 and HILIC chromatography in both positive and negative ion modes to capture a broad range of chemicals on the scalp surface. Scalp oiliness was measured for each participant, and samples for microbial analysis were collected at the same visit. The dataset combines scalp chemical profiles with measured oiliness and accompanying microbiome data, supporting comparisons between scalp types and investigation of chemical variation associated with local microbes. Each participant contributed paired positive- and negative-ion LC–MS/MS acquisitions using both C18 and HILIC chromatography. For each mode-run, peaks in the MS1 spectra representing molecular features were linked to the corresponding MS2 spectra using pyOpenMS [63] with default settings.

### 4.5 Statistics and reproducibility

No statistical method was used to predetermine sample size. Benchmark partitions were disjoint with respect to the evaluated molecules: the MSnLib benchmark was partitioned by molecular formula, and chemical-class assignment and structural-similarity calibration were evaluated by five-fold molecule-disjoint cross-validation, with all spectra from the same molecule kept in the same fold. Classifier thresholds and the cosine-similarity threshold were selected on reference data and applied unchanged to query data. Model training used a fixed random seed.

Unless stated otherwise, 95% confidence intervals are percentile intervals from nonparametric bootstrap resampling of the independent unit of each analysis (parent molecule, molecule, spectral group, query or participant). Comparisons between models used paired resamples.

For the carotid plaque cohort (30 symptomatic and 42 asymptomatic patients), abundance differences between phenotypes were assessed with two-sided Mann–Whitney *U* -tests. Individual metabolites were compared across all 72 patients, with undetected features assigned zero intensity, and the flavonol class was compared among the samples in which it was detected. *P* values are not adjusted for multiple comparisons.

For the scalp cohort, group differences between oily (*n* = 41) and non-oily (*n* = 33) participants are reported as differences in mean log_10_(MS1 intensity + 1) with participant-level bootstrap confidence intervals. Metabolite– microbe associations were estimated in the 55 participants with paired metabolomic and metagenomic profiles by linear regression of standardized variables, adjusting for age, sex, ethnicity, skin sebum, skin hydration and LC–MS acquisition order, with heteroskedasticity-robust (HC3) standard errors. Multiple testing was controlled by the Benjamini–Hochberg procedure within each prespecified family of tests. The robustness of the association between the sphingomyelin family signal and *Corynebacteriaceae* was assessed by 55 leave-one-participant-out fits, five leave-one-feature-out fits and 2,000 participant bootstrap resamples.

### 4.6 UltraAtlas web resource

UltraAtlas can be searched at https://www.yifanliscience.com/ultraatlas/ and explored interactively at https://www.yifanliscience.com/ultraatlas/atlas. Users can upload their own tandem mass spectra to retrieve the closest UltraAtlas regions together with their predicted structural and biochemical classes, and can navigate the full Atlas to inspect individual entries, their representative spectra, class annotations, structure properties and associated study context. The resource is freely accessible without registration.

### 4.7 Code availability

The UltraMS source code, including the model architecture, pretraining and fine-tuning scripts, inference utilities, benchmark evaluation code, and source data, is available at https://github.com/Dsadd4/UltraMS under the Apache License 2.0. UltraMS can be installed from PyPI (pip install ultrams), and the pretrained model checkpoints are available on Hugging Face (https://huggingface.co/collections/dsadd4/ ultrams-6ab4cfaa860cbbd1f9a6b9b6).

### 4.8 Data availability

All tandem mass spectra used for pretraining are publicly available from GNPS/MassIVE (https://massive.ucsd.edu). The UltraMSdata pretraining corpus was assembled from 824,828 open-format files in 3,062 MassIVE datasets listed in a GNPS2 file-level USI export dated 10 August 2025; the complete list of file USIs is available at Zenodo (doi:10.5281/zenodo.24071782). Reference spectra were obtained from MSnLib [26] (doi:10.5281/zenodo.11163380), for which the formula-level and molecule-level benchmark partitions used in this study are available at Zenodo (doi:10.5281/zenodo.23963489) and Hugging Face (https://huggingface.co/datasets/dsadd4/UltraMS-MSnLib-Benchmark); Spectraverse [21] (doi:10.5281/zenodo.17870921); MassSpecGym [27] (https://huggingface.co/datasets/roman-bushuiev/MassSpecGym); NPLIB1 [9, 10]; and Mass-Bank of North America (MoNA; https://mona.fiehnlab.ucdavis.edu). The GeMS-C spectra used for external search are available at https://huggingface.co/datasets/roman-bushuiev/GeMS [18]. The carotid plaque metabolomics data are available from MetaboLights under accession MTBLS12303 [33]. The scalp metabolomics data are available from MassIVE under accession MSV000088283 (doi:10.25345/C5DK1G), and the paired scalp shotgun metagenomes are available from the NCBI Sequence Read Archive under BioProject PRJNA1013332 [51]. Biochemical classes were defined using ChEBI [50]. The MSnLib evaluation subset, fragment annotations and peak representations used for the spectral-interpretation analyses are available at Zenodo (doi:10.5281/zenodo.23227485). Source data are provided with this paper.

## Ethics

This study analysis is based on publicly available, de-identified human datasets; no new samples were collected, no participants were contacted, and no identifiable information was accessed. The carotid plaque metabolomics data (MetaboLights MTBLS12303) were generated in a study approved by the Ethics Committee of Assistance Publique–Hôpitaux de Marseille (No. CSE24-4) and conducted in accordance with the Declaration of Helsinki; the committee waived the requirement for written consent for this observational study, and all patients were informed of their participation by mail with the possibility of withdrawal [33]. The scalp metabolomics (MassIVE MSV000088283) and metagenomic (SRA PRJNA1013332) data were generated under protocols TC-0118-003-095 A and B, based on parent protocol TC-0315-001-006, which were approved by the BASF Tarrytown Consumer Testing Center Institutional Review Board; informed consent was obtained from all volunteers [51].

## Appendix

### S1 Supplementary Methods

The main Methods section describes the UltraMS architecture, objectives and applications. The supplementary material below records the data-construction algorithm, benchmark denominators, evaluation rules and Source Data boundaries needed to reproduce the claims. Detailed rules are written as methods text, and only denominator-defining or configuration-defining information is tabulated.

#### S1.1 UltraMSdata pretraining dataset construction

To construct the pretraining corpus UltraMSdata, we began from a GNPS2 file-level USI export dated 10 August 2025 and downloaded the associated files. We retained GNPS-labelled MassIVE (MSV) open-format records with .mzML or .mzXML extensions, yielding 824,828 file USIs from 3,062 MassIVE accessions. These accessions span diverse experimental contexts: human/cohort, environmental/marine/soil, microbial/microbiome, plant/food/agricultural, chemical-standard/library, exposure/xenobiotic, and model-organism datasets. This distinguishes UltraMSdata from the DreaMS GeMS construction, which selected files from MassIVE datasets explicitly labelled as metabolomics studies bearing the GNPS prefix in their names (as of November 2022). The DreaMS file-level source comprises 338,649 file USIs from 2,211 MassIVE accessions (reported in their paper as 4,467 MassIVE datasets). UltraMSdata extends coverage beyond metabolomics-specific MS^2^ data to include comprehensive scenario including drug discovery, environmental monitoring, clinical, and other application domains.

After downloading, we applied three major preprocessing steps for UltraMSdata construction which is different from the DreaMS/GeMS construction. First, UltraMSdata retained spectra from both positive and negative ionization modes, whereas DreaMS pretraining used only positive-mode centroid spectra. To our knowledge, this constitutes the first inclusion of negative-ion-mode data in a mass-spectrometry foundation model pretraining corpus. Second, we used the total-binned-XIC instrument-accuracy estimate as an inclusion criterion with standard deviation ≤ 10^−3^, a relaxation from the 10^−4^ GeMS-A criterion used in DreaMS pretraining, to admit more diverse instrumentation at the cost of a modest instrument accuracy reduction. Third, many public spectra carry an unknown spectrum-type label. To identify centroid spectra from these files and append them to the pretraining corpus, we replaced the DreaMS/MZmine-style base-peak-span heuristic with a geometry-based centroid/profile estimator. Specifically, the DreaMS estimator infers the spectrum type from a local measurement around the most intense peak: it examines the half-height span of the base peak and uses the presence of zero-intensity points to distinguish profile from thresholded-profile data. This approach is efficient but fragile at repository scale, because a single base peak is not necessarily representative of the sampling geometry across all fragment signals. A regularly sampled or thresholded profile spectrum can therefore appear centroid-like if its base peak happens to be narrow.

The UltraMSdata estimator addresses this failure mode by combining two orthogonal geometric signals. First, the coefficient of variation of adjacent *m/z* spacings separates regular-grid profile acquisition from event-like centroid peak lists. Second, the number of effective sampled points per local peak distinguishes unfolded peak shapes from single-point centroid events. The resulting classifier labels unknown spectra as centroid, profile, thresholded profile, ambiguous, or too sparse. This allowed UltraMSdata to retain centroid-like public spectra despite missing type metadata, while excluding profile-like records.

The overall algorithm is:

**Algorithm S1. Construction of the UltraMSdata**

1. **Data acquisition.** The GNPS2 file-level USI export was retrieved and the corresponding open-format LC-MS/MS files (.mzML/.mzXML) were downloaded from the MassIVE public repository (https://massive.ucsd.edu). Fragment directories matching fragment_* under the frozen download snapshot were enumerated. Checkpointed processing was enabled so that completed and excluded directories were recorded.
2. **Spectrum parsing and metadata extraction.** Because the GNPS2 archive is large, downloads were frequently interrupted by server bandwidth limits, leaving some files blank or truncated. Checkpoint-based recovery was applied over multiple rounds. Files that remained corrupt after all recovery attempts were excluded. The remaining raw files were parsed with the OpenMS Python bindings (pyopenms). File-level properties, including the retention-time ordering flag and the total-binned-XIC instrument-accuracy estimate, were retained for downstream filtering.
3. **Spectrum-level inclusion filtering.** We retained spectra satisfying the following criteria: MS level ≤ 2, 3– 500 fragment peaks, charge from −2 to +2, intensity amplitude ≥10, total-binned-XIC standard deviation ≤ 10^−3^, precursor *m/z ≤* 1500, and fragment *m/z ≤* 1500. Files with non-ordered retention time were skipped. The parsed spectra and file-level properties were assembled with pandas/NumPy and materialized as Apache Parquet shards via pyarrow.
4. **Centroid/profile classification.** Centroid spectra were retained and profile spectra excluded. For spectra reported as unknown, the type was estimated from peak geometry. Let Δ*m_i_* = *m_i_*_+1_ *m_i_* after removing non-positive spacings, and compute CV*_m/z_* = sd(Δ*m*)*/*mean(Δ*m*). Let *ɛ* = 10^−6^ max(*I*) and define points-per-peak as the number of points with intensity *> ɛ* divided by the number of local maxima *I_i_*_−1_ *< I_i_ I_i_*_+1_ above *ɛ*. Unknown spectra with fewer than five points were marked too sparse to classify and retained. Regularly sampled spectra (CV*_m/z_ <* 0.1) were rejected when points-per-peak *>* 20 (profile) or *>* 5 (thresholded profile), and retained only when classified as ambiguous. Event-like spectra (CV*_m/z_* ≥ 0.1) were retained as centroid when points-per-peak *<* 5, and otherwise retained as ambiguous.
5. **Training normalization.** The Parquet shards were read with pyarrow.parquet. Intensities were normalized to [0, 1] by dividing by the within-spectrum maximum. The 150 highest-intensity fragments were retained and sorted by increasing *m/z* using NumPy/PyTorch.

The final UltraMSdata corpus used for pretraining contained 160,641,162 tandem mass spectra from GNPS/MassIVE, comprising 149,767,344 positive-ion spectra, 10,793,129 negative-ion spectra, and 80,689 spectra with other or missing ion-polarity annotations. Spectra with missing ion-polarity annotations were retained for masked-peak reconstruction but excluded from the ion-polarity prediction task.

#### S1.2 Development, stability and transferability of chemical representations

*Development of chemical representations during training*

To further characterize how UltraMS captures chemical properties during training, we tracked the development of its representations across masked-peak reconstruction pretraining, adaptation with the auxiliary task of retention-time prediction, and joint training with the auxiliary tasks of retention-time and polarity prediction (Supplementary Fig. 1). We evaluated successive model states using a fixed set of 90,737 spectra from 9,533 molecules, none of which were included in the training corpus. Reconstruction accuracy was evaluated at three mass scales: 10, 1 and 0.1 Da. To examine the molecular information encoded in the representations, we additionally measured molecular fingerprint similarity, molecular identification accuracy, retention-time correlation and ionization-polarity discrimination. Molecular fingerprints were estimated using a linear model fitted on embeddings from 7,626 molecules and evaluated on a separate set of 1,907 molecules. Molecular identification was assessed by retrieving another spectrum of the same molecule, with accuracy defined as the fraction of queries for which the highest-ranked match had the correct molecular identity.

As training progressed, UltraMS increasingly captured the intrinsic chemical features needed to distinguish molecular identities, even when the reconstruction loss fluctuated. Molecular identification accuracy rose from 85.9% at the end of masked-peak reconstruction pretraining to 87.7% after incorporating the auxiliary task of retention-time prediction and 88.0% after further incorporating polarity prediction during joint training. This progression shows that incorporating the two auxiliary tasks of retention-time and polarity prediction strengthened the molecular information encoded by UltraMS despite fluctuations in the reconstruction objective.

*Stability of chemical representations under spectral perturbations*

We next examined whether UltraMS preserves its representations when observed spectra are perturbed through missing peaks, intensity variation or shifts in spectral peak *m/z* values (Supplementary Fig. 2). We aimed to simulate changes encountered in real-world applications, such as differences between instruments or variations in experimental conditions that alter the mass, intensity or occurrence of spectral peaks, and to test whether representations learned by different foundation models would be disrupted by these practical challenges. We randomly sampled 10,000 spectra from the MSnLib test set and applied four types of perturbation to each spectrum: random removal of 10–50% of spectral peaks, Gaussian noise added to normalized intensities with standard deviations of 0.05–0.20, random variation in spectral peak *m/z* values within ±0.05 to ±0.5 Da, and retention of only the 50, 30, 20 or 10 most intense peaks. UltraMS and DreaMS were evaluated using the same original and perturbed spectra. Representation stability was quantified by the cosine similarity between the global embedding of each original spectrum and that of its perturbed counterpart.

**Supplementary Figure 1.**
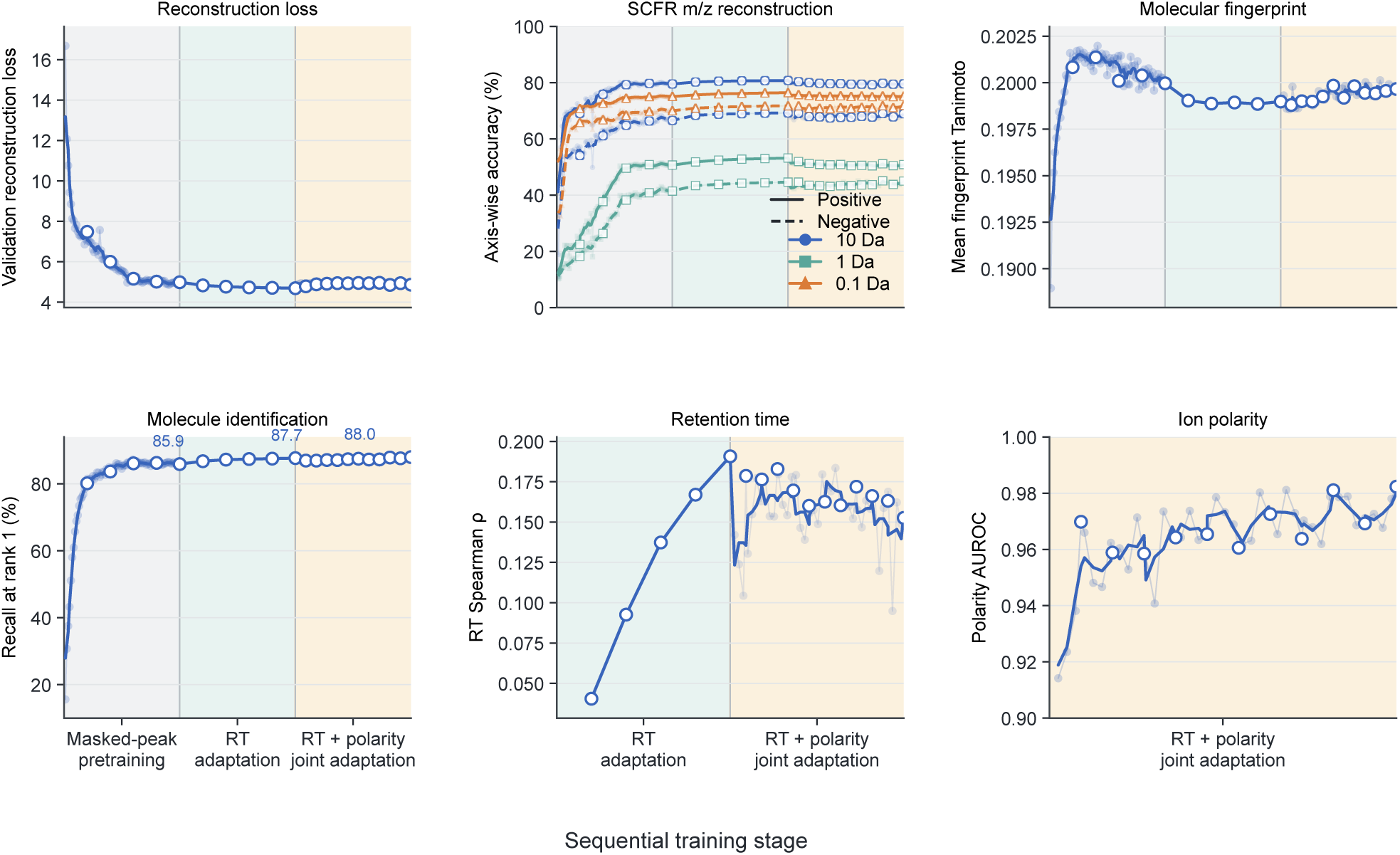
Development of chemical representations during UltraMS training. Trajectories of reconstruction loss, reconstruction accuracy at three mass scales, molecular fingerprint similarity, molecular identification accuracy, retention-time correlation and ionization-polarity discrimination. The three training stages comprise five epochs of masked-peak reconstruction pretraining, five epochs of adaptation with the auxiliary task of retention-time prediction, and eleven epochs of joint training with masked-peak reconstruction and the auxiliary tasks of retention-time and ionization-polarity prediction. Colours distinguish reconstruction accuracy at the 10, 1 and 0.1 Da mass scales, while solid and dashed lines distinguish positive- and negative-ion spectra. Molecular fingerprint similarity was calculated as the mean Tanimoto similarity between predicted and reference Morgan fingerprints. Molecular identification accuracy was evaluated as recall at rank 1 using 17,896 query spectra for which another spectrum of the same molecule was available for retrieval. Retention-time prediction was evaluated by calculating Spearman’s correlation between predicted and measured retention times for 14,187 spectra. Ionization-polarity prediction was evaluated using AUROC on 17,935 spectra.

**Supplementary Figure 2.**
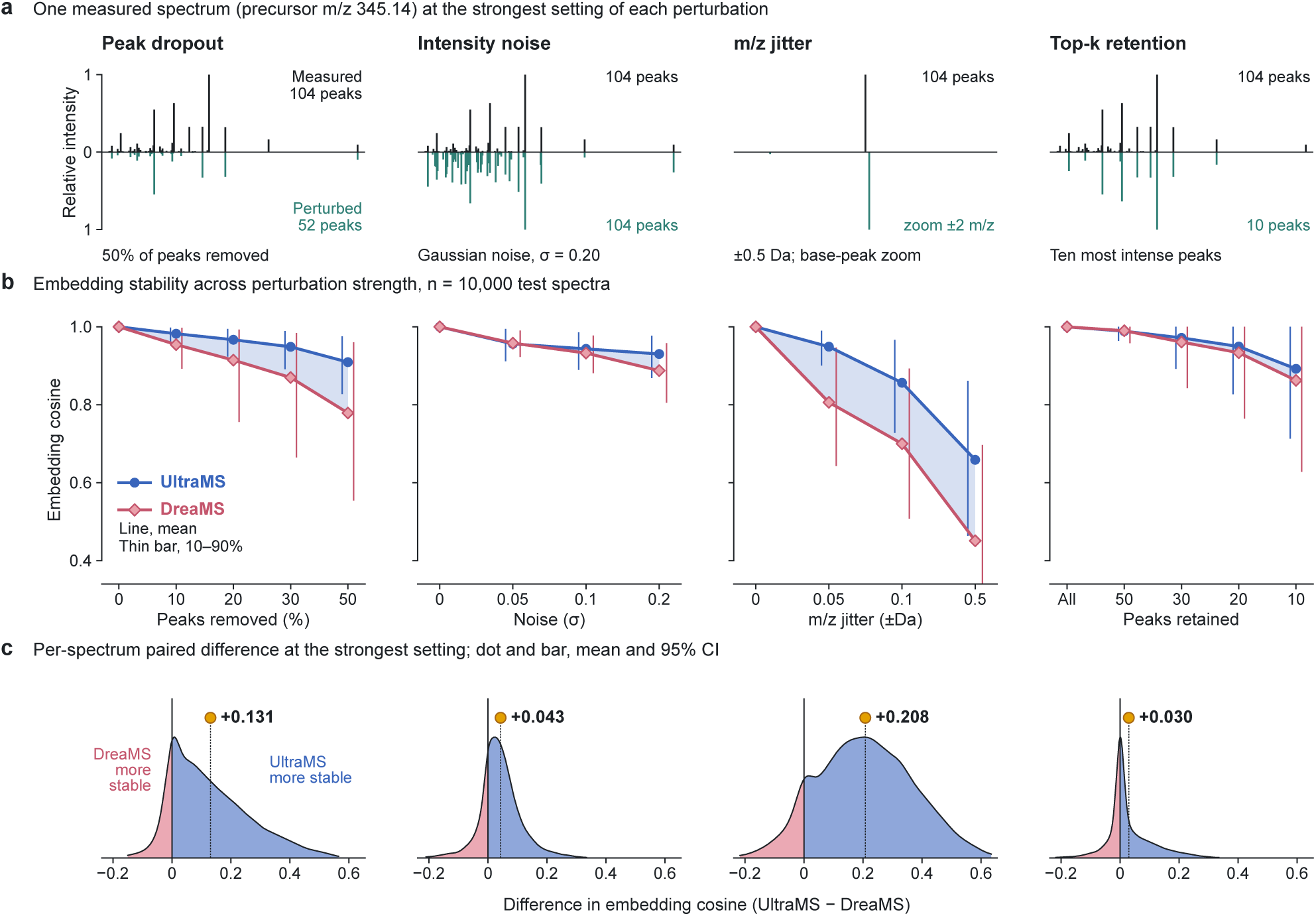
Stability of global representations under spectral perturbations. a, An example illustrating the four types of perturbation. **b,** Cosine similarity between the original and perturbed global embeddings across increasing perturbation strengths; shading marks the gap between the mean curves of the two models. **c,** Distributions of the per-spectrum differences in embedding similarity between UltraMS and DreaMS under the strongest perturbation settings. Positive values (blue) indicate greater stability for UltraMS and negative values (pink) greater stability for DreaMS. Coloured points and horizontal intervals show the mean differences and their 95% confidence intervals, calculated from 2,000 bootstrap resamples of the spectra.

We show that UltraMS retained greater similarity to the original representation under the strongest setting of each perturbation. After removal of half of the peaks, the mean embedding similarity was 0.909 for UltraMS and 0.779 for DreaMS. With intensity noise of *σ* = 0.20, the corresponding values were 0.930 and 0.888. Random shifts in spectral peak *m/z* values within 0.5 Da produced the largest separation between the models, with similarities of 0.659 and 0.451, respectively. When only the ten most intense peaks were retained, UltraMS maintained a similarity of 0.892, compared with 0.863 for DreaMS. These results show that UltraMS preserves a more consistent spectral representation space when peak information is incomplete or altered, supporting the stability of its learned chemical representations across variations in spectral observations.

##### Recognition of fragment structures when collision energies varied

We then examined whether spectral peak-level embeddings are able to preserve fragment identity when collision energy changes the surrounding fragmentation pattern (Supplementary Fig. 3). We analyzed 13 negative-ion molecular collision-energy series from MSnLib, using MAGMa annotations to identify shared fragment structures across spectra acquired from the same molecule at different collision energies. This yielded 965 cross-energy comparisons. For each query peak, spectral peaks in a spectrum acquired at another collision energy were ranked by embedding similarity. Identification accuracy was calculated as the fraction of comparisons in which the corresponding fragment with the same structure ranked first, and was then averaged equally across the molecular collision-energy series.

The illustrated collision-energy series shows four shared fragments across spectra acquired at 20, 30, 45 and 60 eV (Supplementary Fig. 3), despite pronounced changes in peak intensities and the overall spectral pattern. Across the 13 molecular collision-energy series, UltraMS identified the corresponding fragment at rank 1 in 91.2% of comparisons on average, compared with 14.6% for DreaMS. The mean difference was 76.7 percentage points, with a 95% confidence interval of 72.2–81.4 percentage points. This demonstrates that UltraMS peak embeddings retained a strong correspondence to fragment structure across changes in collision energy, linking chemically equivalent spectral peaks even when their spectral surroundings differed substantially.

**Supplementary Figure 3.**
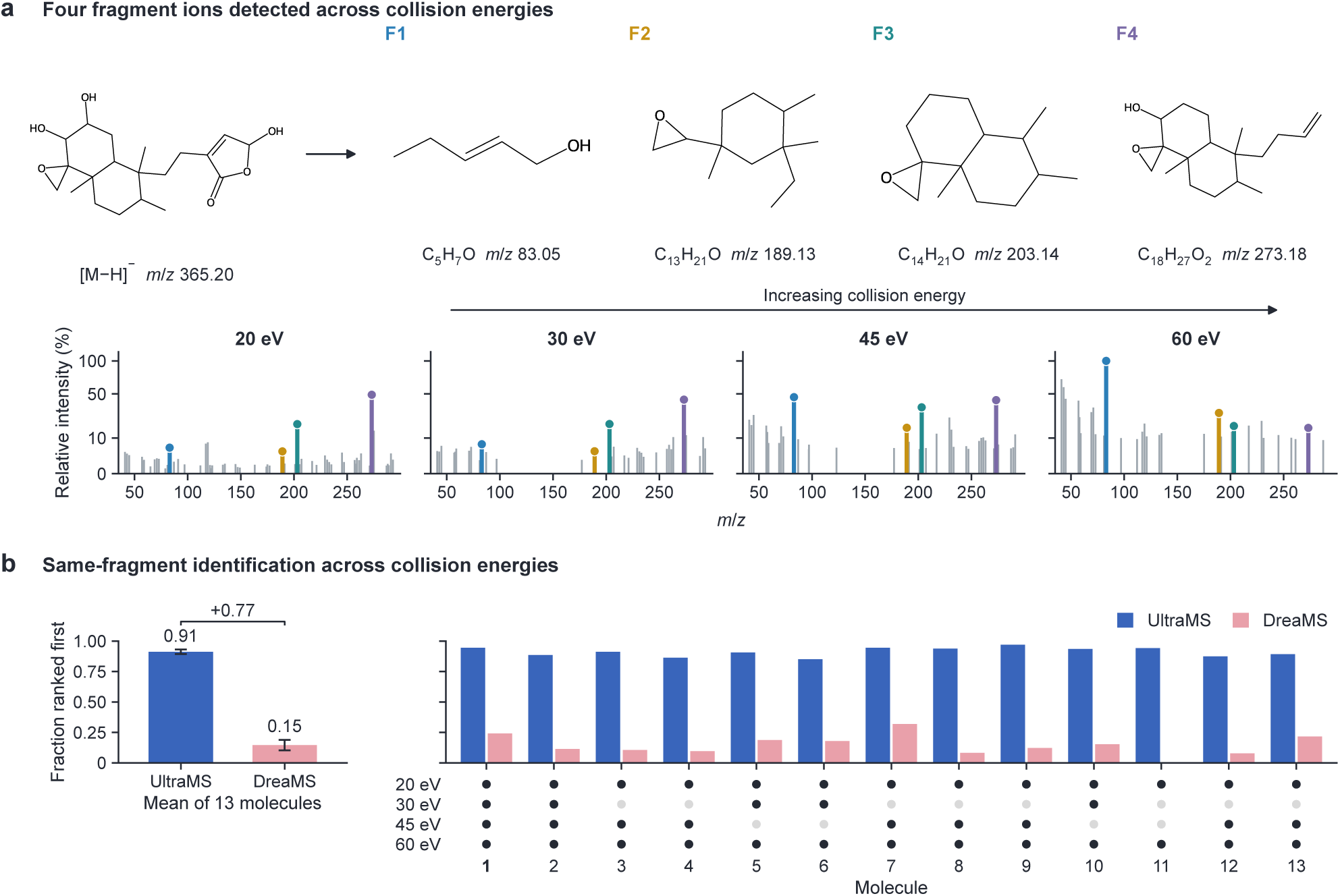
Recognition of unchanged fragment structures across collision energies. a, Spectra from one molecular collision-energy series acquired in negative-ion mode at 20, 30, 45 and 60 eV. Four MAGMa-annotated fragment structures observed at all four collision energies are linked by color to their corresponding peaks. **b,** Rank-1 identification accuracy for unchanged fragments across collision energies, averaged equally across 13 molecular collision-energy series containing 965 eligible comparisons. The confidence interval for the difference between models was calculated using 10,000 bootstrap resamples of the molecular collision-energy series.

##### Transfer of fragment identity across different molecules

Finally, we examined whether the structural information captured by spectral peak embeddings extends to fragments with the same structure originating from spectra of different molecules (Supplementary Fig. 4). We analysed 7,371 query peaks from negative-ion-mode spectra of 253 held-out molecules in MSnLib. For each query peak, we retrieved the ten most similar reference peaks with annotated structures from spectra of other molecules. A query was counted as successfully matched when at least one of these neighbours had the same MAGMa-annotated fragment structure. For each molecule, we calculated the fraction of its query peaks that were successfully matched and then averaged the differences between UltraMS and DreaMS across the 253 molecules.

The results show that UltraMS found structurally matching fragments across different molecules for 851 query peaks, compared with 219 for DreaMS. When matching rates were calculated separately for each molecule, UltraMS achieved a higher matching rate than DreaMS for 193 of the 253 molecules and a mean improvement of 8.9 percentage points across all 253 molecules, with each molecule given equal weight (95% confidence interval, 7.8–10.1 percentage points). Three examples illustrate how these correspondences connect chemically distinct molecules through shared sulfur dioxide, thiocarbamoyl and trifluoromethyl fragments (Supplementary Fig. 4). For these query fragments, UltraMS retrieved 10, 8 and 7 structurally matching fragments from other molecules among the top ten, respectively, whereas none of the spectral peaks retrieved by DreaMS had a compatible structure. These results show that UltraMS captures fragment-level chemical features that remain recognizable across both acquisition conditions and different molecular structures, providing a path for transferring fragment annotations between compounds.

**Supplementary Figure 4.**
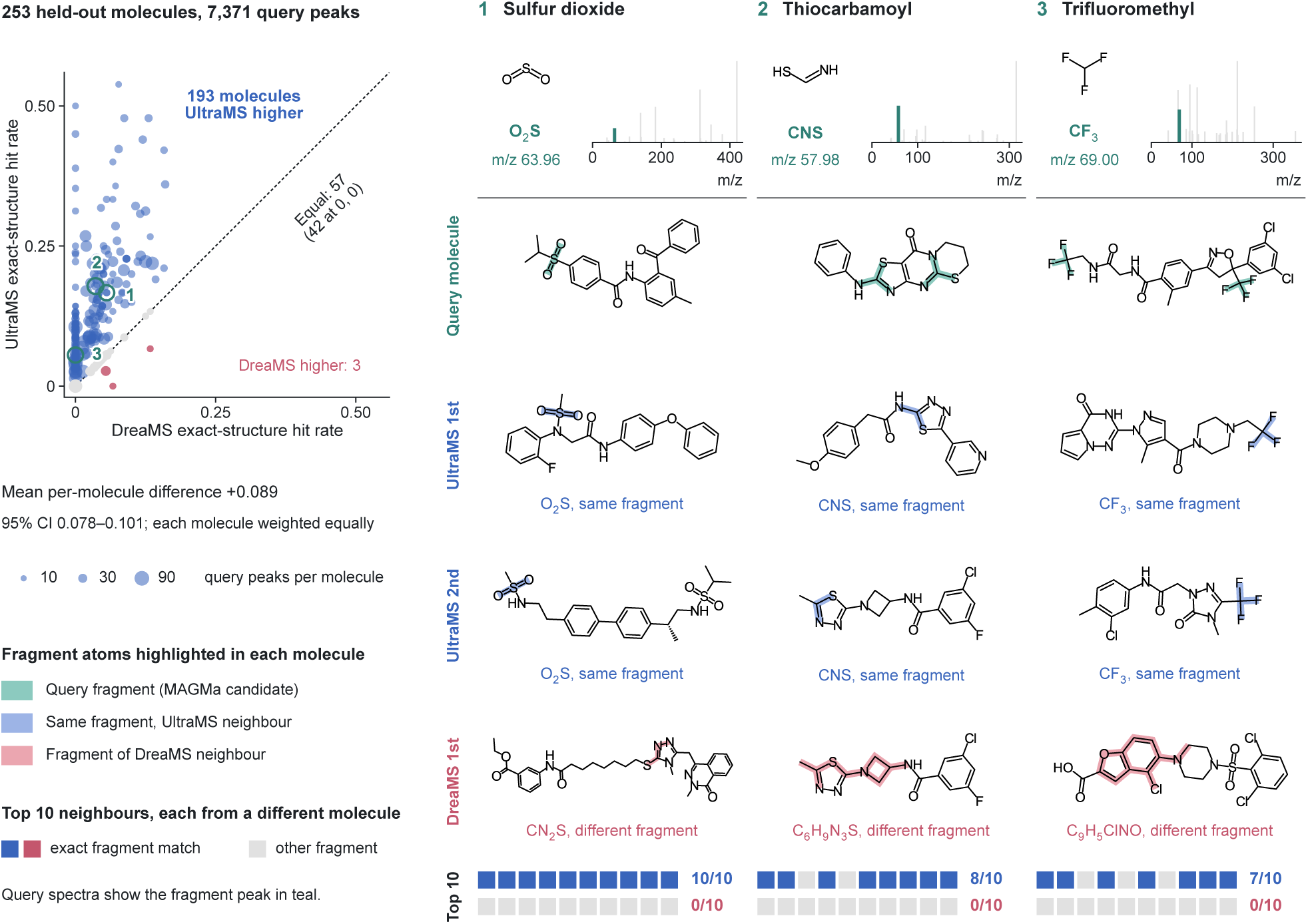
Transfer of fragment identity across different molecules. The upper panel shows the difference between UltraMS and DreaMS in the fraction of query peaks with an exact structural match among their ten nearest neighbours, calculated separately for 253 molecules. The lower panels illustrate fragment examples of sulfur dioxide, thiocarbamoyl and trifluoromethyl. Each example shows the query spectrum, the annotated fragment structure and the highest-ranked reference molecule retrieved by each model.

#### S1.3 Learned chemical semantics support molecular identification across diverse benchmarks

To apply UltraMS to spectrum-to-molecule identification, we extracted its global spectrum embedding and peak-level embeddings from each query spectrum and used a simple attention module to aggregate information across spectral peaks. The resulting spectral representations were used to predict molecular fingerprint information, and candidate molecules were ranked according to the similarity between their molecular fingerprints and the spectrum-derived predictions. Using this approach, we further examined whether the learned chemical semantics of UltraMS support molecular identification across different public benchmarks and candidate settings (Supplementary Fig. 5).

We evaluated UltraMS on the NPLIB1 natural-product retrieval benchmark and two molecule-retrieval tasks from MassSpecGym. The NPLIB1 evaluation contained 3,699 held-out query evaluations from three folds, representing 3,352 unique spectra and 1,844 molecules. The two MassSpecGym tasks each contained 16,440 test spectra. The mass-restricted task required the model to distinguish the correct molecule from candidates with similar molecular masses, whereas the formula-restricted task further limited the candidates to molecules sharing the same molecular formula. UltraMS and DreaMS were evaluated using the same query spectra and candidate sets within each task. Identification accuracy was measured by whether the correct molecule appeared among the first 1, 5 or 20 ranked candidates. Published results from fingerprint-based models and other molecular identification methods were included for the corresponding NPLIB1 and MassSpecGym tasks.

On NPLIB1, UltraMS achieved Top-1, Top-5 and Top-20 identification accuracies of 33.0%, 57.9% and 80.5%, respectively, compared with 27.9%, 54.6% and 79.1% for DreaMS. On the MassSpecGym mass-restricted task, UltraMS achieved 17.1%, 33.3% and 54.9%, compared with 13.2%, 26.3% and 45.7% for DreaMS. On the formula-restricted task, UltraMS achieved 14.1%, 29.0% and 51.7%, whereas DreaMS achieved 11.4%, 24.0% and 43.4%. UltraMS achieved the highest Top-1 identification accuracy among the displayed methods across all three tasks and consistently outperformed DreaMS at Top-1, Top-5 and Top-20. As the retrieval depth increased, UltraMS remained strongly competitive with architectures developed specifically for these molecular identification tasks, using only its pretrained spectral embeddings and a simple attention module to aggregate peak-level information. These results demonstrate that the learned chemical semantics of UltraMS enable accurate molecular identification across natural-product retrieval and candidate sets restricted by molecular mass or formula.

##### Following Top-1 candidates across molecules and acquisition conditions

Chemists judge a proposed structure not only as right or wrong but also by how close it comes to the true molecule, and they gain confidence when the same structure is supported under several acquisition conditions. To view spectrum-to-molecule identification in the same way, we followed the Top-1 candidate of every MSnLib test spectrum in the single-spectrum identification of Fig. 3a across all spectra of its molecule and measured its structural distance from the true molecule (Supplementary Fig. 6). The analysis covered all 57,437 test spectra of the 4,805 test molecules (median 10 spectra per molecule across adducts and collision energies) and the six models of Fig. 3a: UltraMS, DreaMS and the Linear, Codebook, Fourier and DeepSets baselines.

##### Classes of Top-1 candidates

Each Top-1 candidate was assigned to one of four classes. <u>Correct</u> denotes the Top-1 hits of Fig. 3a. <u>Same connectivity</u> denotes a candidate with the atomic connectivity of the true molecule that differs only in stereochemistry or representation. For all other candidates, we counted the bonds not shared with the true molecule, that is, the bonds of the two molecules minus twice the bonds of their maximum common substructure. The maximum common substructure, a single connected substructure, was computed with RDKit rdFMCS, matching elements and bond orders, matching ring bonds only to ring bonds and allowing only complete rings, with a time limit of 20 s per pair; for a candidate with several disconnected components, the largest component was used. <u>Close</u> denotes a candidate with the molecular formula of the true molecule and at most four unshared bonds, and <u>Distant</u> denotes all other candidates. Moving a terminal substituent such as a hydroxyl group to another atom of the scaffold leaves two bonds unshared, so a Close candidate is the true scaffold with up to two such moves. Bonds were compared on the molecular graph.

##### Incorrect Top-1 candidates are mostly distant isomers

For every model, an incorrect Top-1 candidate was almost always a structurally distinct molecule rather than a variant of the true one: 98.4% to 99.1% of incorrect candidates were Distant, differing from the true molecule by a median of 38 to 39 bonds (Supplementary Fig. 6a,b). Identification in this benchmark is therefore largely all-or-none: a model either recognizes the structure from the spectrum or ranks an unrelated isomer first, so the higher Top-1 accuracy of UltraMS reflects more molecules whose core structure is identified. The rare near-misses, Close or same-connectivity candidates, keep the scaffold and composition of the true molecule and differ only in stereochemistry or in the positions of a few substituents; they were most frequent for UltraMS (1.60% of incorrect candidates, compared with 1.24% for DreaMS and 0.89% to 1.03% for the baselines), and peak-level substructure annotations (Fig. 3c) can help to place the remaining substituents.

**Supplementary Figure 5.**
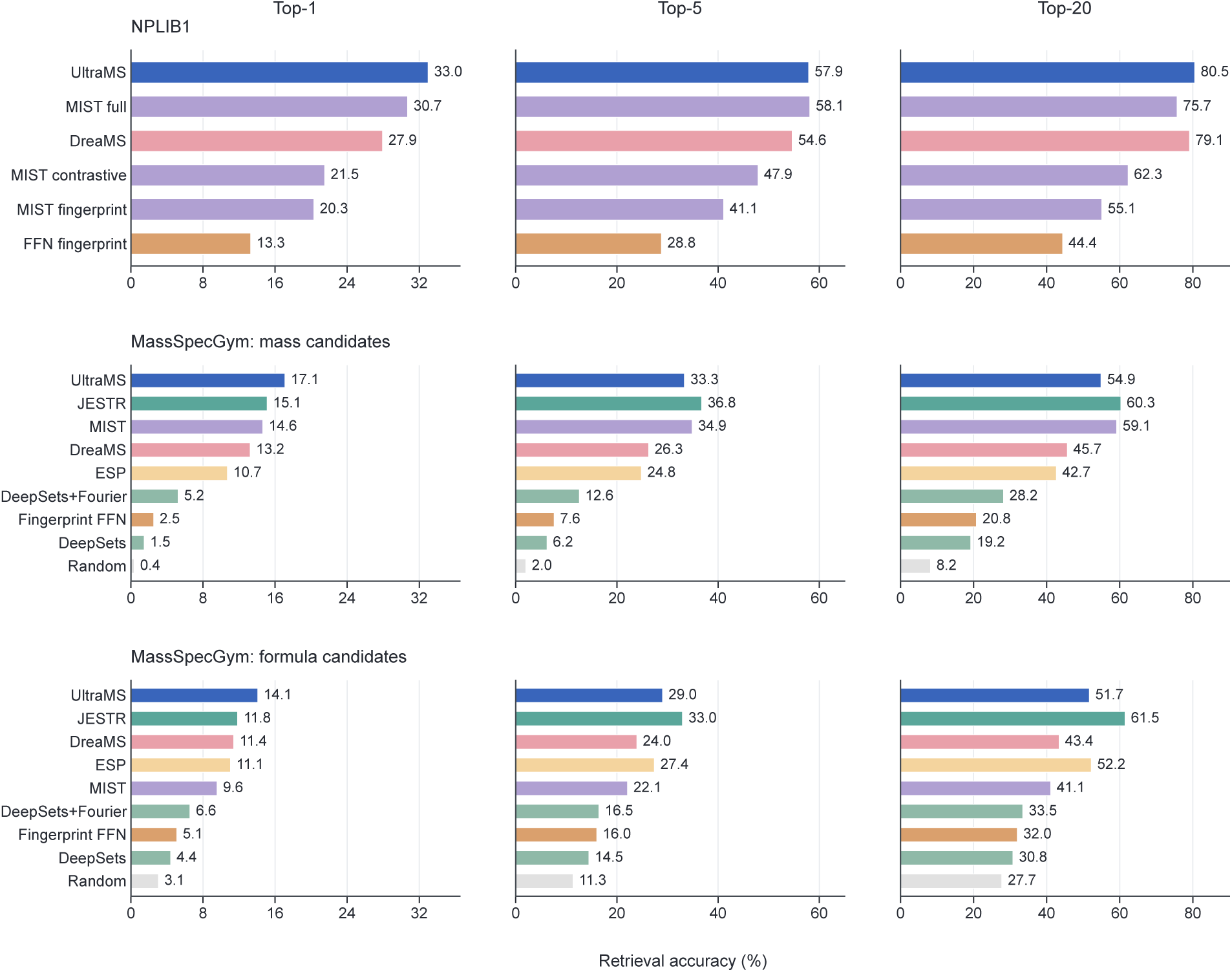
Molecular identification across diverse benchmark settings. Rows show the NPLIB1 natural-product retrieval benchmark, the MassSpecGym mass-restricted candidate task and the MassSpecGym formula-restricted candidate task. Columns show Top-1, Top-5 and Top-20 identification accuracy, defined as the percentage of query spectra for which the correct molecule appeared among the first 1, 5 or 20 ranked candidates. The UltraMS and DreaMS evaluation on NPLIB1 contained 3,699 held-out query evaluations from three folds, representing 3,352 unique spectra and 1,844 molecules. Each MassSpecGym task contained 16,440 test spectra.

##### UltraMS reveals the correct structure of most molecules from at least one acquisition condition

Each test molecule was measured as several spectra, a median of ten, that differ in adduct and collision energy. For every molecule, we counted the spectra in which the true molecule was ranked first and grouped the molecules accordingly into molecules identified from at least one of their spectra, molecules identified from more than half of their spectra and molecules identified from all of their spectra (Supplementary Fig. 6c). With UltraMS, 2,913 of the 4,805 molecules (60.6%) were identified from at least one of their spectra, more than with any other model (53.3% with DreaMS and 12.1% to 41.8% with the baselines); 1,121 molecules (23.3%; 14.2% with DreaMS) were identified from more than half of their spectra, and 239 molecules (5.0%; 2.1% with DreaMS) from all of their spectra. Most molecules were therefore identified from some of their spectra but not from others; cytidine, for example, was identified from 44 of its 71 spectra. Whether a single spectrum leads to the correct structure thus depends on the adduct and collision energy at which it was acquired, and multiple-spectrum voting (Fig. 3a), which pools all spectra of a molecule, makes use of every condition under which the molecule is identified.

##### Example molecules

Supplementary Fig. 6d–h shows five example molecules. For each molecule, the grid on the left has one row per model and one column per test spectrum; a cell is green if the model ranked the true molecule first for that spectrum, light green if it ranked first a candidate with the same connectivity, yellow if it ranked first a Close candidate and grey if it ranked first a Distant candidate. On the right are the true structure and the incorrect candidates that UltraMS or DreaMS ranked first most often. Two molecules were identified by UltraMS from all of their spectra: the steroidal alkaloid cyclopamine (seven spectra) and a cyclohexylpiperazine sulfonamide (PubChem CID 46396345; eight spectra). Two molecules were identified from only some of their spectra, and which spectra depended on the adduct and on the model. Cytidine has 71 test spectra with four adducts; UltraMS identified it from 44, including 18 of the 20 formate-adduct ([M + HCOO]^−^) spectra, and DreaMS from 36, including all 30 [M + H]^+^ spectra. A thiophene-carbonyl diazepane (PubChem CID 137889994) has eight test spectra; UltraMS and DreaMS each identified it from four, and only one of these spectra was shared, so that together the two models identified it from seven of the eight. The last molecule shows Close candidates, isomers that keep the scaffold of the true molecule but carry a substituent at another position. Iriflophenone was not identified from any of its eight spectra by any model, but in five of them UltraMS ranked first a trihydroxybenzophenone with the same rings and hydroxyl groups as the true molecule, with the hydroxyl groups at other positions of the trihydroxyphenyl ring.

We next considered a challenging protocol for spectrum-to-spectrum based molecule identification, in which a spectrum acquired at 20 eV was used to search a library of spectra acquired at 40 eV. Because the same molecule fragments substantially differently under low- and high-collision-energy conditions, the two spectra may share little m/z overlap, so successful retrieval requires the model to retain molecule-level fragmentation semantics rather than merely spectral-level similarity. We observe that UltraMS maintained the leading position across the major [M+H]^+^ and [M H]^−^ ion modes, achieving Recall@1 values of 94.7% and 92.4%, respectively, compared with 88.6% and 86.3% for DreaMS and only 22.9% and 24.7% for the strongest non-pretrained spectrum encoders. We further examined four major categories representing prevalent scientific scenarios and practical applications. For stereochemically complex compounds containing at least four stereocentres, which are common among steroid metabolites and natural products, UltraMS achieved Recall@1 values of 94.3% and 89.9% in positive and negative mode, respectively, supporting robust cross-energy annotation of structurally intricate metabolites. For molecules containing at least three aromatic rings, a category relevant to aromatic pharmaceuticals and environmental chemicals, UltraMS achieved 95.0% and 93.3%. For carboxylic-acid-containing compounds, which include organic acids, fatty acids, bile acids and numerous drug metabolites, UltraMS achieved 91.7% and 90.3%, exceeding DreaMS by 7.0 and 9.2 percentage points, respectively, an advantage that translates into more reliable annotation in metabolomics and drug-metabolite studies. For halogenated compounds containing F, Cl, Br or I, which are prominent among pharmaceuticals, agrochemicals and environmental contaminants, UltraMS achieved 96.0% and 93.5%, supporting robust cross-energy library matching in drug and environmental screening.

In the isomer discrimination analysis of Fig. 3c (bottom panels), spectra with substructure annotations supporting only the correct isomer outnumbered those supporting only the competing isomer by 16.3-fold and 11.4-fold in the correctly and incorrectly identified cases for UltraMS, respectively, compared with 6.3-fold and 4.3-fold for DreaMS.

In spectrum-to-spectrum library search on MSnLib (Fig. 3e), DreaMS required higher FDRs of 4.58%/7.23% for [M+H]^+^/[M H]^−^, as compared with 2.89%/5.29% for UltraMS, to reach their respective maximum recalls. In cross-ionization spectrum-to-spectrum library search (Fig. 3f), UltraMS yielded 256 and 257 additional correct Top-1 identifications compared with DreaMS for positive-ion and negative-ion queries, respectively, and placed the true molecule in the top ten for 510 and 572 more spectra.

**Supplementary Figure 6.**
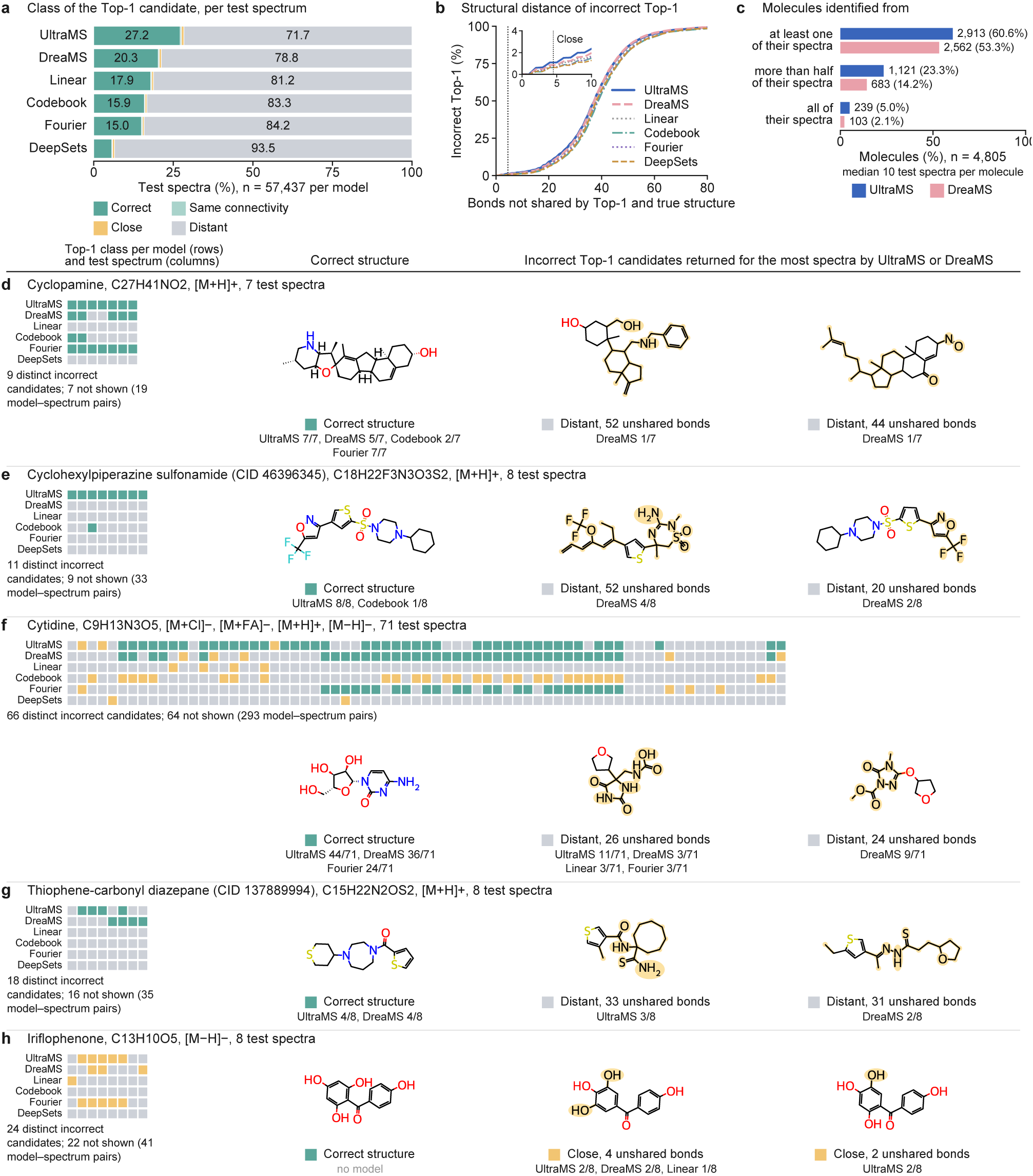
Top-1 candidates followed across molecules and acquisition conditions. **a**, Class of the Top-1 candidate for the 57,437 MSnLib test spectra and six models. Close, same molecular formula and at most four bonds not shared with the true structure; Same connectivity, differing only in stereochemistry or representation. **b**, Cumulative distribution of bonds not shared by incorrect Top-1 candidates and the true structure; dotted line, Close threshold. **c**, Test molecules (*n* = 4,805) identified from at least one, more than half or all of their test spectra. **d**–**h**, Example molecules. Left, Top-1 class per model (rows) and test spectrum (columns). Right, the correct structure and the two incorrect candidates most often ranked first by UltraMS or DreaMS, with atoms outside their common substructure with the correct structure highlighted.

**Supplementary Figure 7.**
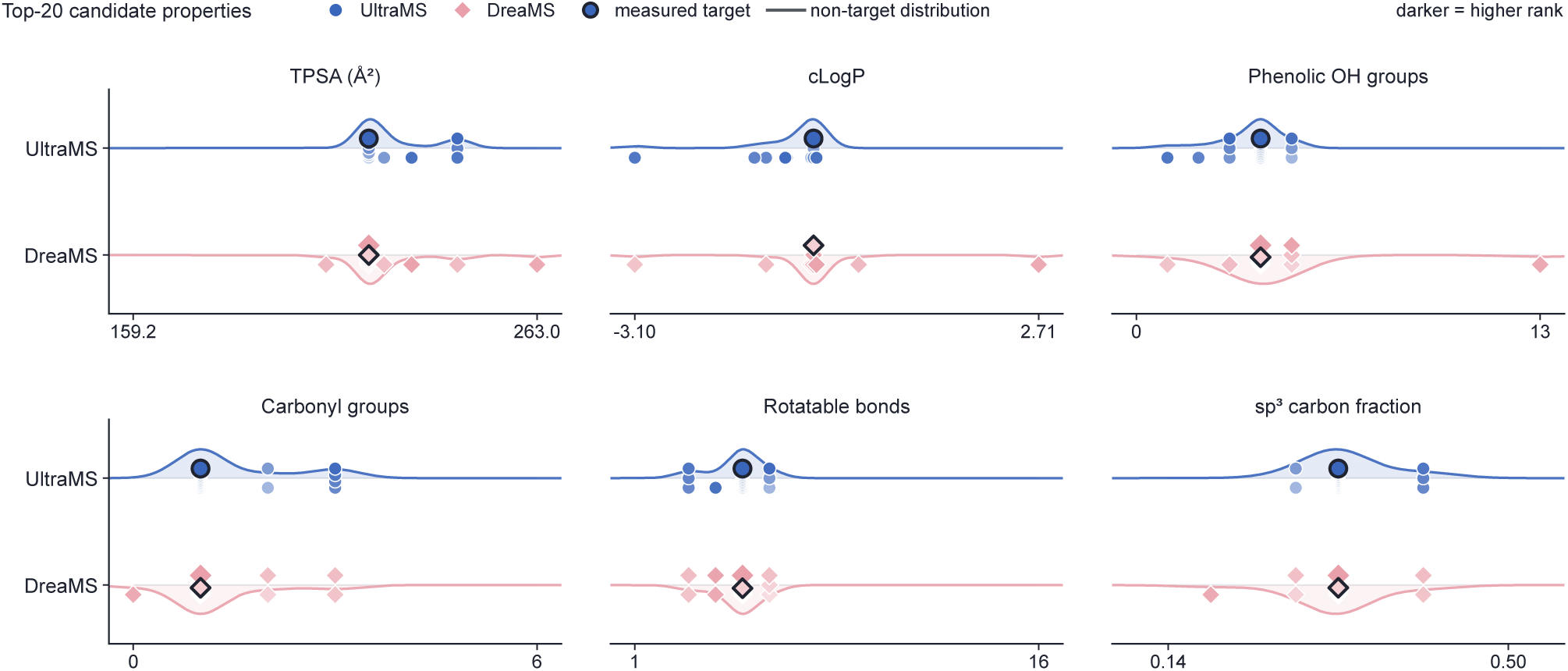
Property distributions of the top 20 candidates, with outlined symbols marking the ground-truth structure in. **Fig. 3c**.

**Supplementary Figure 8.**
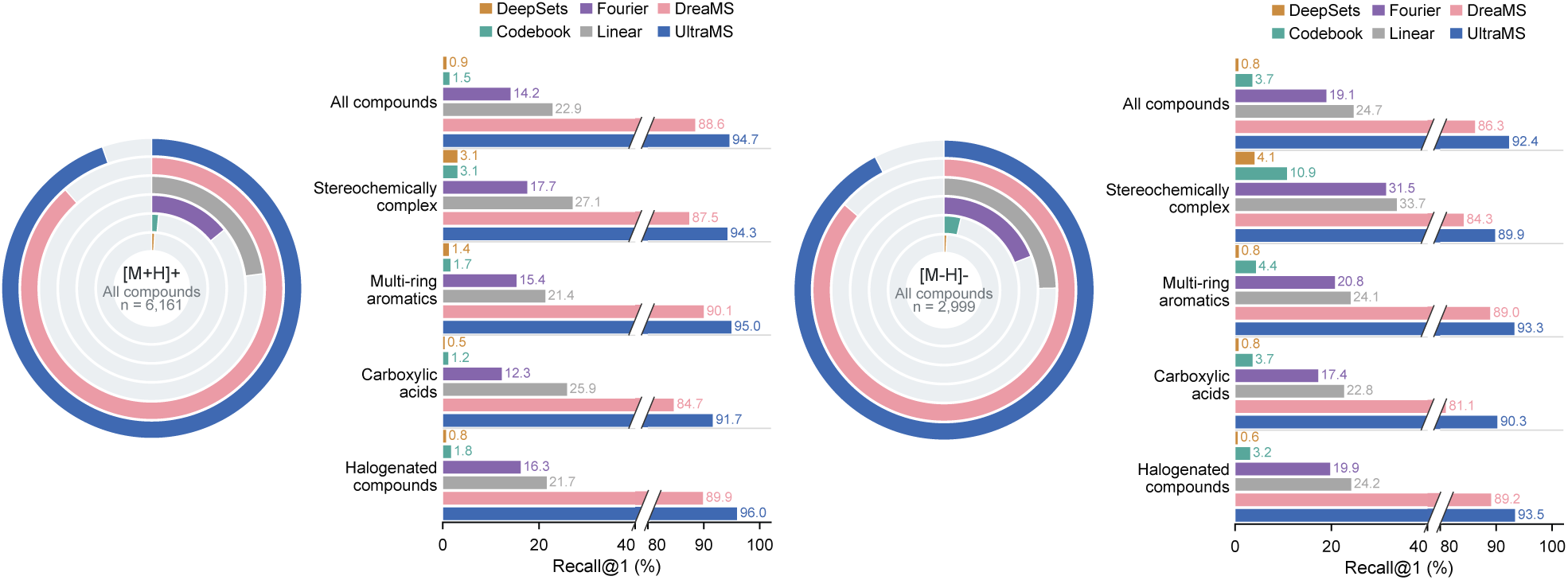
Robustness of cross-collision-energy library search under different acquisition conditions. Low-energy query spectra (*≤* 20 eV) are searched against high-energy reference spectra (*≥* 40 eV). UltraMS achieves the highest Recall@1 across both positive- and negative-ion modes, spanning diverse chemical classes including stereochemically complex compounds, polycyclic aromatics, carboxylic acids, and halogenated molecules. Ring plots summarize overall identification performance across all compounds; bar plots compare overall and class-specific identification accuracies.

#### S1.4 Spectral peak embeddings enable chemical interpretation of tandem mass spectra

##### Datasets, evaluation protocol and ground-truth substructures

We evaluated the spectral peak annotation transfer approach on the original MSnLib test partition, comprising 57,437 spectra from 4,805 parent molecules, with all 502,647 training and validation spectra as the reference library. UltraMS and DreaMS were applied as frozen pretrained encoders without further training. Spectral peaks of the query and reference spectra were annotated by MAGMa using their known parent molecules; these computational spectral peak substructures served only to assess the results. Specifically, within each query spectrum, we compared the cosine similarity of peak-level embeddings against the Tanimoto similarity (Morgan fingerprints) of their annotated substructures across all annotated peak pairs.

##### Intensity-weighted and edit-tolerant evaluation of transferred annotations

The advantage of UltraMS in exact substructure assignment (18.4% versus 5.0% of spectral peaks) persisted when considering spectral prominence: when spectral peaks were weighted by their intensity, accurately annotated spectral peaks accounted for nearly a fifth of the total spectral intensity in UltraMS (19.8%; 95% CI, 19.2–20.4%), compared to only 6.5% (95% CI, 6.1–6.9%) for DreaMS. Furthermore, allowing the transferred and true substructures to differ by up to two atom or bond edits yielded only a marginal gain (to 20.9% and 6.5%, respectively). This indicates that the transferred spectral peak annotations are driven by conserved, identical substructures across different molecules.

##### Transfer of substructure annotations to spectral peaks unannotated by MAGMa

A transferred substructure was retained only if its atomic connectivity and bond types were supported by the query parent molecule and its calculated *m/z* matched the observed *m/z* of the unannotated target peak within a 10 ppm mass tolerance. To evaluate whether transferred annotations also preserved broader chemical fidelity, we compared physicochemical descriptors between transferred and MAGMa-annotated substructures (Fig. 4d). UltraMS achieved lower macro-averaged mean absolute errors across all four descriptors: Bertz structural complexity (60.0 versus 143.5 for DreaMS), topological polar surface area (12.1 versus 20.7 Å^2^), log *P* (0.633 versus 1.011), and the number of rotatable bonds (1.18 versus 1.60).

#### S1.5 UltraMS reveals phenotype-associated metabolites in clinical carotid plaques

##### Literature evidence for glutathione in carotid plaque biology

Human carotid-plaque studies provided direct support for the protective role of the glutathione-dependent antioxidant system. In one study, the antioxidant system that uses and regenerates glutathione was severely weakened in carotid plaques, and patients whose plaques completely lacked detectable glutathione-peroxidase activity were more likely to have disease in multiple blood vessels or narrowing of the opposite carotid [37]. In another study of 48 carotid plaques, unstable plaques contained 64% less of a glutathione-dependent detoxification enzyme than stable plaques (*P* = 0.006) [38]. Experimental studies showed the same protective direction: glutathione supplementation reduced plaque area by 30% in mice [39], whereas stimulating glutathione production reduced oxidative damage and existing plaque burden [40].

##### Literature evidence for shikimate in carotid plaque biology

Shikimate (shikimic acid) was detected by UltraMS in 15 of 42 asymptomatic plaques but only 3 of 30 symptomatic plaques. An animal study showed that shikimate reduced blood-clot formation in the carotid without causing detectable bleeding [64]. This finding is directly relevant because carotid plaques commonly produce symptoms when a blood clot forms on a disrupted plaque [33]. Therefore, the UltraMS annotations offer computational insight into a plausible mechanism through which higher shikimate levels may be associated with reduced clotting activity, ultimately lowering the risk of symptomatic plaques.

##### Reassessment of the original picolinic acid assignment

The chromatograms provided an immediate clue. In electrospray ionization, the same molecule commonly appears as paired protonated and sodium-adduct ions [65]. If the *m/z* 141.0657 ion were picolinic acid [*M* + NH_4_]^+^, the corresponding neutral molecule would have a mass of approximately 123.032 Da, leaving both the co-eluting *m/z* 163.0478 peak and the observed 21.9821-Da mass difference unexplained. In contrast, MS-FINDER, an established in silico structure-elucidation tool that scores candidate molecules according to how well their structures explain the observed MS2 peaks under chemically defined molecular dissociation principles [41], favored MIAA over picolinic acid for all four plaque spectra. The paired ions can be well explained by MIAA [*M* + H]^+^ at *m/z* 141.0657 and MIAA [*M* + Na]^+^ at *m/z* 163.0478, and MS-FINDER scores for MIAA were nearly 0.5 higher on average than those for picolinic acid.

##### Literature evidence for the kynurenine pathway in carotid plaque biology

Beyond the individual abundance variations of kynurenine (KYN), anthranilic acid (AA) and kynurenic acid (KYNA), plaque-level profiles demonstrated consistent covariation between KYN and AA alongside a positive correlation for KYN–KYNA. Altering expression of kynurenine pathway enzymes in carotid plaques has been linked to local inflammation and plaque instability [45]. Additionally, elevated circulating levels of KYNA and AA have been independently associated with plaque instability [66] and severe carotid artery narrowing [67]. In light of this evidence, the higher abundance of KYN and its downstream products in symptomatic plaques raises the hypothesis that a similar metabolic imbalance occurred in the MTBLS12303 cohort, potentially contributing to adverse events.

S1.6 UltraAtlas reveals hidden molecular relationships across diverse mass spectra

##### UltraMS preserves molecular identity across spectral variation

On the 6,159 challenging queries presented in the main text, among the spectrum pairs in the top 10% of embedding similarity for each method, direct spectral cosine similarity matching and neural network encoding baselines reached mean ECFP4 Tanimoto similarities of 0.586–0.721 (Fig. 6b), and no method other than UltraMS and DreaMS exceeded a pooled Pearson *r* of 0.13 (Fig. 6c). To complement the analysis of the 6,159 challenging queries presented in the main text, we evaluated UltraMS on the remaining 16,756 less challenging queries from the same independent test set (Supplementary Fig. 9). For each query, the query spectrum itself was excluded, and all other spectra with the same ion polarity were ranked according to their similarity to the query. Molecular identification accuracy was calculated as the fraction of queries for which the highest-ranked spectrum came from the correct molecule. To determine whether the retrieved results also reflected structural similarity, we calculated the ECFP4 Tanimoto similarity between the query molecule and the molecule represented by the highest-ranked spectrum.

Across these 16,756 queries, raw cosine identified the correct molecule for 87.75% of queries, showing that direct peak overlap already provided a strong identification signal in this subset. DreaMS increased the identification accuracy to 93.61%, while UltraMS reached 95.67%. Furthermore, the molecules retrieved by UltraMS were more structurally similar to the query molecules, with a mean top-ranked ECFP4 Tanimoto similarity of 0.967, compared with 0.951 for DreaMS and 0.910 for raw cosine similarity. These results show that UltraMS improves molecular identification even when direct spectral similarity is already informative, while also retrieving molecules that more closely resemble the query structure.

**Supplementary Figure 9.**
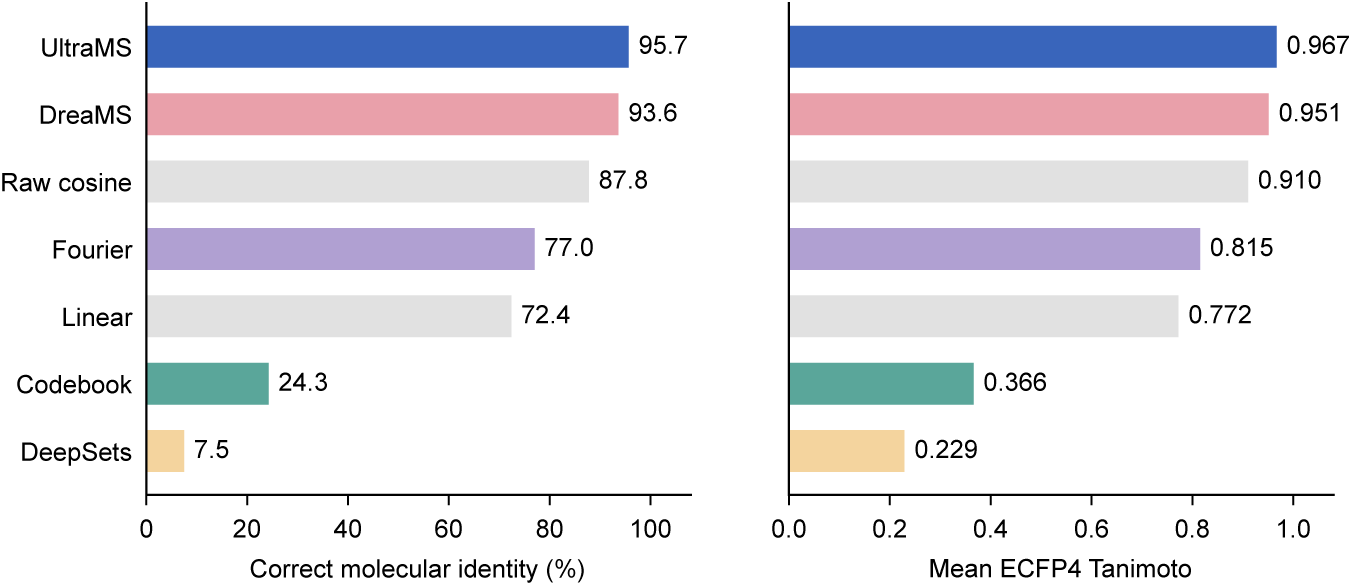
Molecular identification when direct spectral similarity is informative. The left panel shows the percentage of queries for which the highest-ranked spectrum came from the correct molecule. The right panel shows the mean ECFP4 Tanimoto similarity between the ground-truth molecule and the molecule represented by the highest-ranked spectrum.

##### Chemical classes form coherent regions across the UltraAtlas landscape

To determine whether the molecular organization revealed by UltraAtlas reflected shared chemistry rather than direct spectral resemblance alone, we examined whether UltraMS could efficiently infer structural and biochemical classes from large numbers of untargeted mass spectra (Supplementary Figs. 10–12). First, we defined two complementary label systems: ten structural classes defined according to RDKit principles and 20 ChEBI biochemical classes. Classes were mutually exclusive within each system, such that each molecule belonged to one structural class and one biochemical class when available. The normalized 512-dimensional UltraMS embeddings of all spectra from the same molecule were averaged to obtain one molecular representation. Molecular-class prediction models were then trained for the RDKit and ChEBI label systems, respectively.

Class organization was evaluated using five molecule-disjoint folds, with each molecule used once for evaluation and excluded from the corresponding training set. For each class, we selected the probability threshold that retained the largest number of molecular assignments while keeping the proportion of incorrect assignments at or below 5%. We found that all ten RDKit structural classes could be assigned while maintaining an error rate of no more than 5% (Supplementary Fig. 10). Phosphate/cofactor-like and sugar/polyol-like molecules showed the broadest coverage, with recalls of 86.17% and 74.12%, respectively. Additionally, among the 20 ChEBI biochemical classes shown in Supplementary Figs. 11 and 12, 19 could be assigned while maintaining an error rate of no more than 5%. Together, these results show that UltraMS embedding is capable to organize diverse mass spectra according to their shared structural and biochemical features.

##### UltraAtlas transfers structural knowledge to large-scale unannotated spectra

To evaluate the retrieval completeness of UltraAtlas, we compared its two-stage search with direct search across all 163.7 million spectra in UltraMSdata on 10,000 external spectrum queries from GeMS-C: UltraAtlas returned the same spectrum as full-library search for 99.93% of queries, with the remaining results falling firmly within the top five candidates of full-library search. We next examined whether UltraAtlas could extend its chemical organization to large-scale external mass spectra beyond the reference datasets used in its construction. We sampled 1,000,000 spectra from GeMS-C across different data sources, ion polarities and precursor *m/z* ranges (Supplementary Fig. 13). After removing duplicate spectra, we further excluded queries whose retrieved Atlas matches had a raw spectral cosine similarity greater than or equal to 0.95, ultimately leaving 750,464 spectra for the external-coverage analysis (Supplementary Fig. 13a). We show that UltraAtlas maintained high search efficiency, taking 13.52 ms per query spectrum with a cold cache, which required the relevant data to be loaded from local storage, and 10.11 ms per query with a warm cache, in which the data had already been loaded into the server’s memory (Supplementary Fig. 13b). Additionally, the searched spectra covered a broad precursor *m/z* range, with most spectra distributed between 100 and 800 (Supplementary Fig. 13c).

We then examined whether the Atlas matches returned for these external spectra represented structurally related molecules (Supplementary Fig. 14). We began with 22,915 reference spectra from 2,830 molecules with known structures and compared UltraMS spectral similarity with ECFP4 molecular-structure similarity, obtaining a molecule-level Pearson correlation of 0.614 (Supplementary Fig. 14a). A five-fold molecule-disjoint calibration was then used to convert UltraMS spectral similarity into expected ECFP4 similarity. After establishing this relationship between UltraMS embedding similarity and molecular-structure similarity, we applied the calibrated mapping to the 750,464 external spectra. The Atlas returned matches with a mean expected ECFP4 similarity of 0.972. By comparison, randomly paired Atlas spectra with the same ion polarity and similar precursor mass had a mean expected similarity of only 0.155 (Supplementary Fig. 14b,c). The mean difference was 0.817 (95% confidence interval, 0.808–0.831; *P* = 1.0×10^−5^). These results show that UltraAtlas can recommend structurally relevant reference spectra for large-scale external queries even after highly similar spectral matches are removed. UltraAtlas therefore holds the potential to transfer molecular knowledge from structurally characterized reference spectra to the largely unannotated mass-spectral landscape.

##### UltraAtlas uncovers diverse chemical classes among previously unclassified scalp metabolic products

We next examined how UltraAtlas complements the original analyses performed using GNPS, SIRIUS and CANOPUS when applied to the untargeted scalp metabolomics dataset (Supplementary Fig. 15). The scalp study comprised 74 participants analysed across four LC–MS modes. Spectral features were combined and deduplicated by acquisition mode and feature identifier, yielding 32,538 molecule features with associated tandem mass spectra. Original chemical-class labels were obtained from the GNPS, SIRIUS and CANOPUS analyses. For UltraAtlas, the chemical class of the Atlas recommendation was transferred when its UltraMS cosine similarity met the positive-or negative-ion threshold established at an empirical FDR of 5%.

**Supplementary Figure 10.**
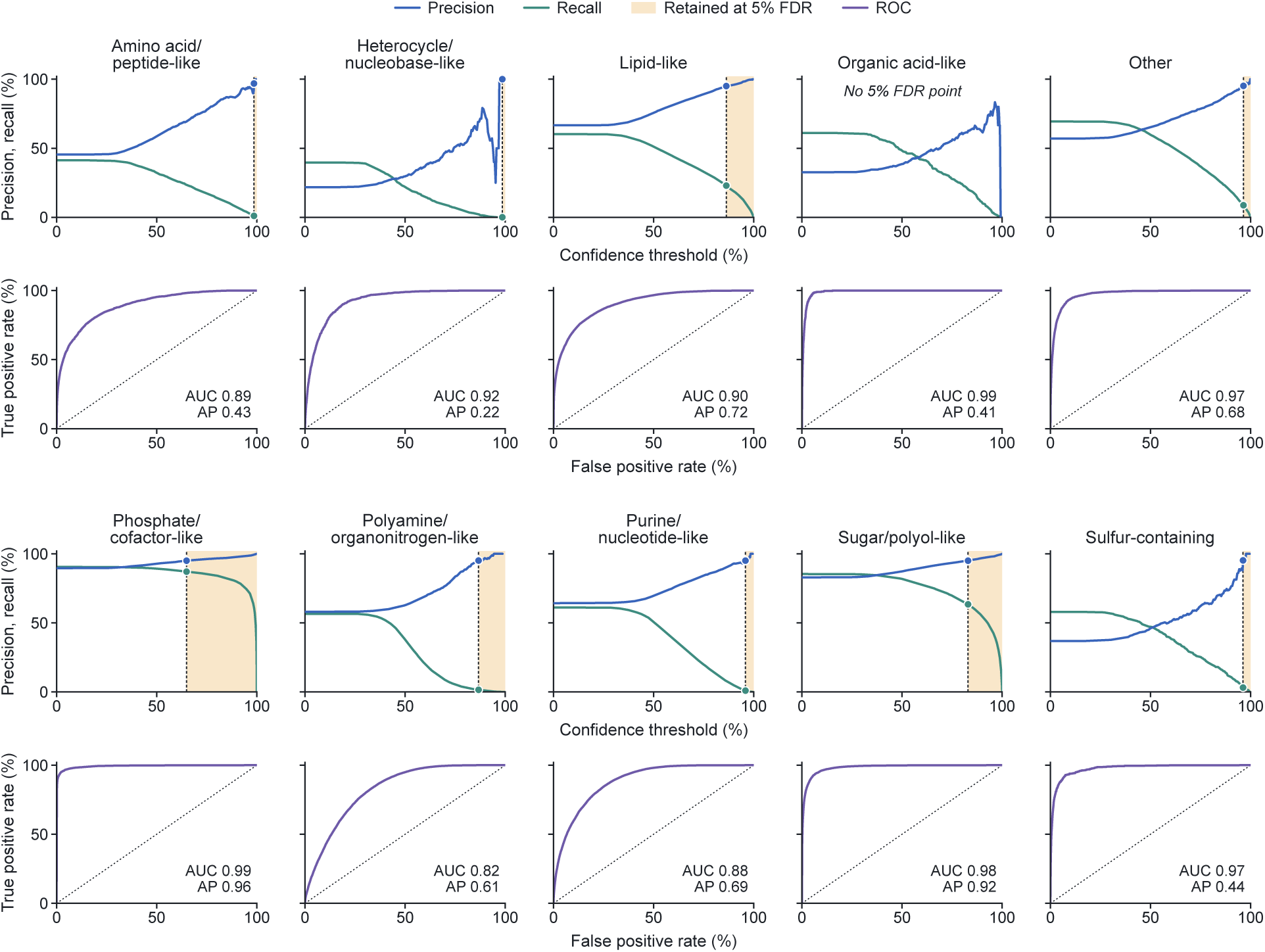
RDKit structural-class assignment. Precision and recall across probability thresholds, together with the corresponding ROC curves, for ten RDKit structural classes evaluated by five-fold molecule-disjoint cross-validation. Dotted lines mark the selected class-specific thresholds at a 5% error rate, and open circles indicate precision and recall at these thresholds.

**Supplementary Figure 11.**
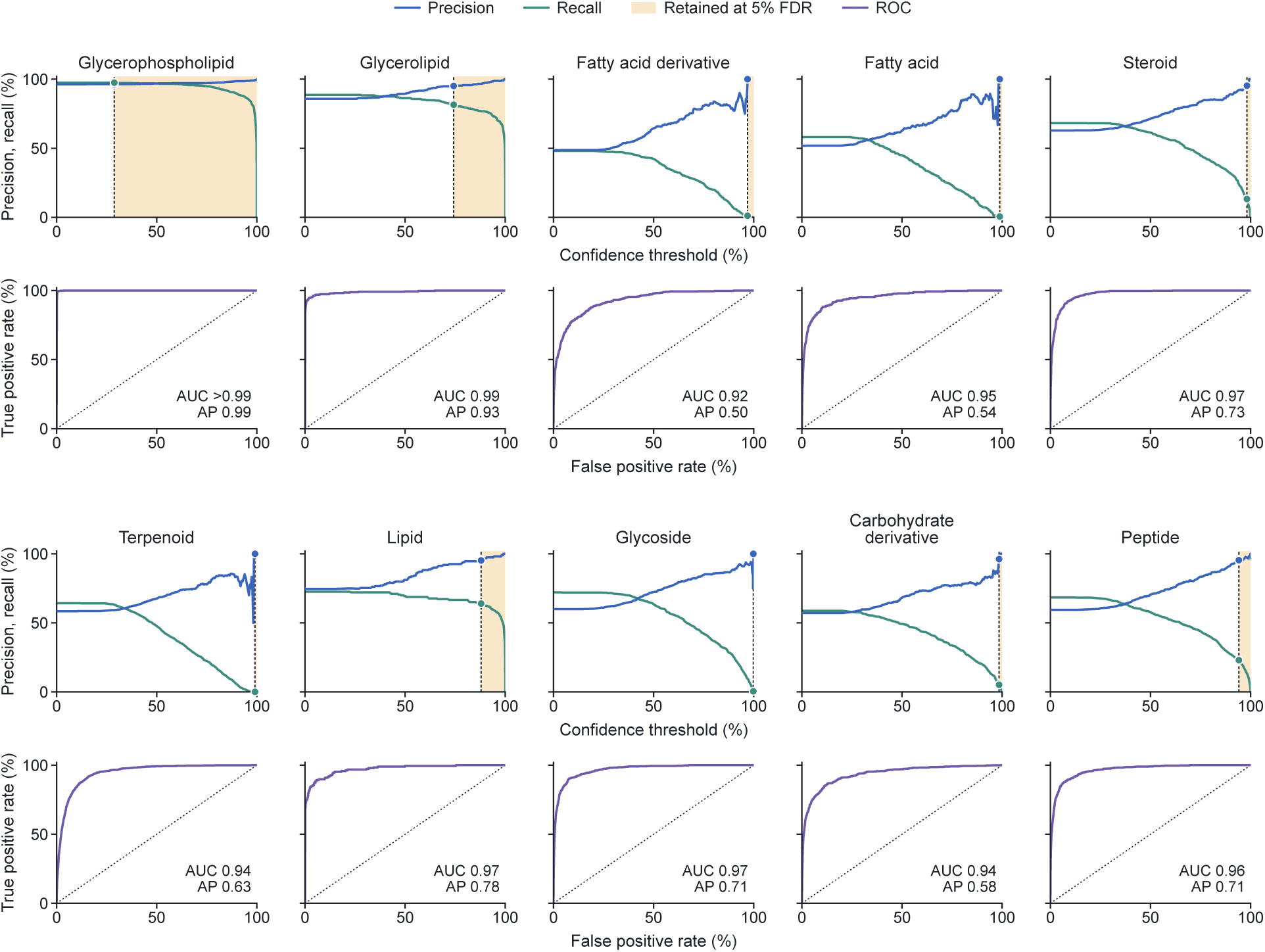
ChEBI biochemical-class assignment, part. **I.** Precision and recall across probability thresholds, together with the corresponding ROC curves, for ten ChEBI biochemical classes evaluated by five-fold molecule-disjoint cross-validation. Dotted lines mark the selected class-specific thresholds at a 5% error rate, and open circles indicate precision and recall at these thresholds.

**Supplementary Figure 12.**
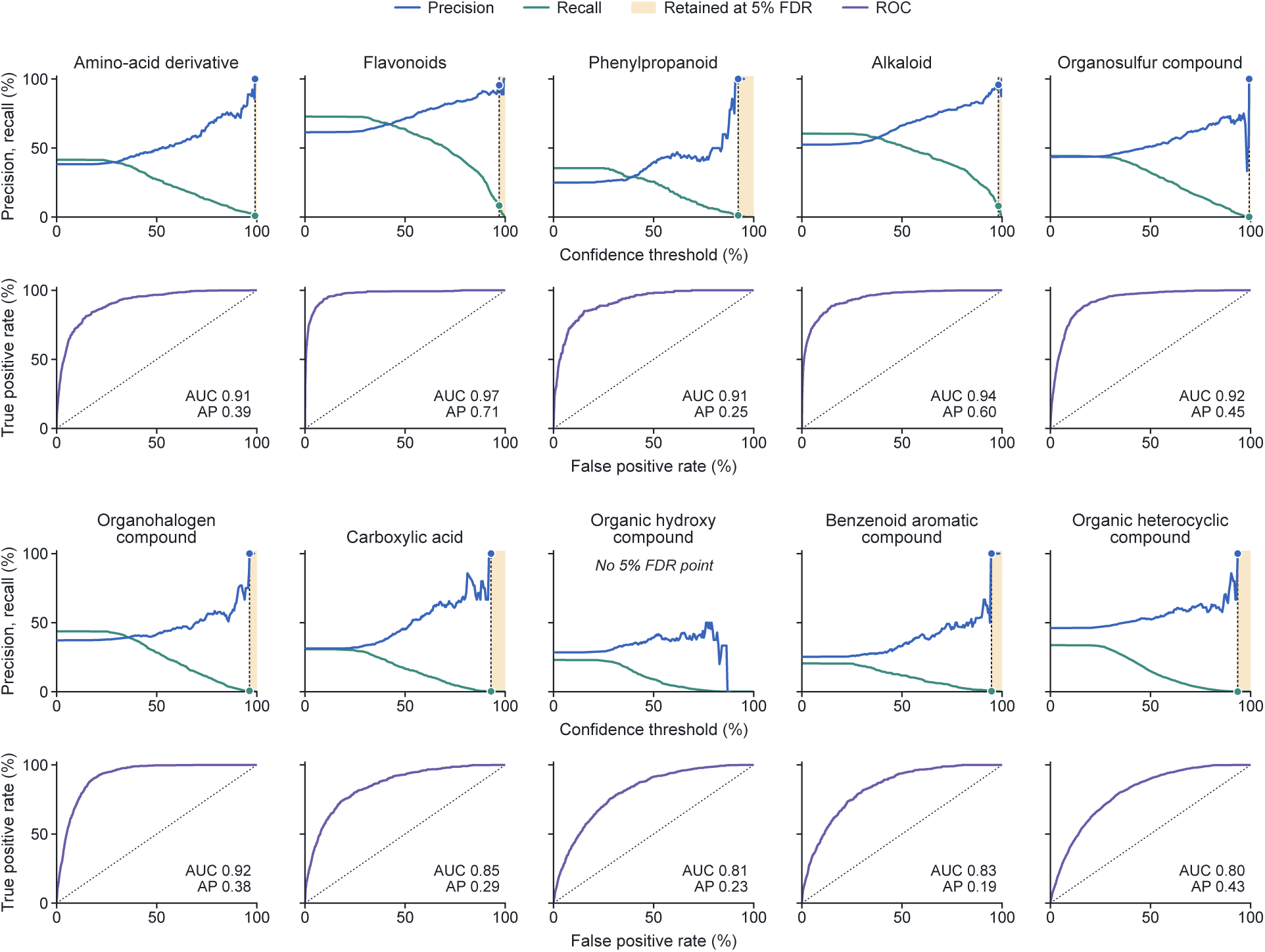
ChEBI biochemical-class assignment, part II. Precision and recall across probability thresholds, together with the corresponding ROC curves, for additional ChEBI biochemical classes evaluated by five-fold molecule-disjoint cross-validation. Dotted lines mark the selected class-specific thresholds at a 5% error rate, and open circles indicate precision and recall at these thresholds. Particularly, for organohalogen, no predictions could be retained at a 5% error rate.

**Supplementary Figure 13.**
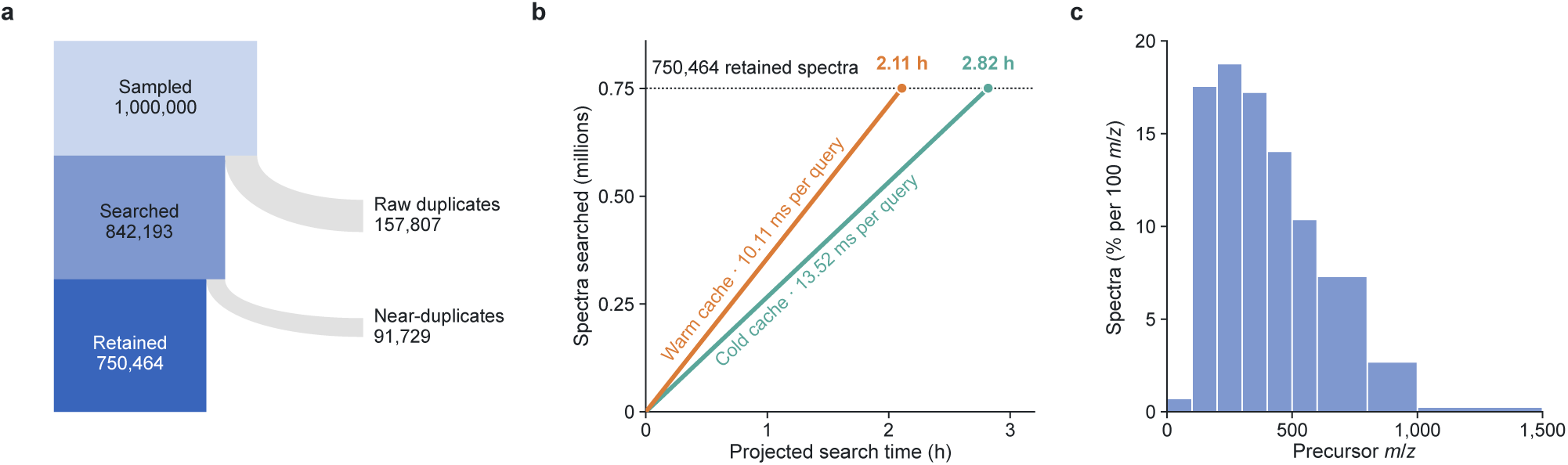
Large-scale search of external spectra. **a**, Numbers of GeMS-C spectra initially sampled, searched after raw-spectrum deduplication and retained after the exclusion of near-identical query–Atlas matches. Band widths are proportional to the numbers of spectra; grey branches indicate the removed spectra. **b**, Projected time for the retained external spectra, derived from the search time per query spectrum under cold- and warm-cache conditions. Reported search times refer to the final spectrum-matching step. **c**, Precursor *m/z* distribution of the searched spectra; bar widths follow the *m/z* bins and heights give the percentage of spectra per 100 *m/z*.

We show that the original tools provided chemical-class labels for 16,236 molecule features (49.9%), whereas UltraAtlas assigned chemical classes to 10,391 molecule features (31.9%). Combining these annotations increased the coverage to 21,099 molecule features (64.8%), with UltraAtlas adding class labels for 4,863 molecule features that lacked an original label. The gain was observed across all four acquisition modes and ranged from 10.5% in C18 negative-ion mode to 77.6% in HILIC negative-ion mode. These results show that UltraAtlas substantially expands the chemical-class information obtained from untargeted mass spectra, bringing previously unclassified molecule features into the chemical organization of the scalp metabolite.

##### UltraAtlas places scalp microbe–metabolite associations in chemical context

The chemical regions linked to individual skin microbes in Fig. 6i agree with the established ecological niches and nutritional requirements of the scalp microbiome: *S. epidermidis* occupies a metabolic niche rich in phosphatidylcholines and amide-linked fatty acids [51]; *Propionibacteriaceae* become more abundant as skin oil and its neutral-lipid components increase [51]; and *Malasseziaceae* must obtain fatty acids from their surroundings owing to the absence of fatty-acid synthase [68]. For the sphingomyelin family in Fig. 6k, all five individual SM family molecule features also showed positive associations with *Corynebacteriaceae* relative abundance, in addition to the family-level association. To profile microbial functional potential, we quantified the abundances of KEGG Orthology (KO) groups and Enzyme Commission (EC) numbers in the paired scalp shotgun metagenomes, yielding 4,895 candidate functions that were tested for association with the SM family signal.

#### S1.7 Model training, adaptation and retrieval settings

We primarily used two UltraMS model checkpoints. The first was the self-supervised UltraMS model with retention-time and ion-polarity adaptation, which provided general spectrum-level and spectral-peak-level representations. The second was a contrastively enhanced UltraMS model trained to place spectra from the same molecule close to one another. This model provided the 512-dimensional representations used for the spectrum-to-spectrum retrieval experiments in Fig. 6 and for the construction of UltraAtlas. We additionally examined a model obtained by adapting the self-supervised UltraMS model on the public MoNA molecular-identity contrastive dataset. This MoNA-adapted model was used for the embedding visualization and was further explored in the supplementary NPLIB1 experiments, where it was compared with DreaMS contrastive training enhanced version using the same training protocol.

##### Self-supervised UltraMS representation

The general UltraMS model was trained using the UltraMSdata pretraining corpus containing 160,641,162 tandem mass spectra. When more than 150 valid fragmention spectral peaks were present, the 150 spectral peaks with the highest normalized intensities were retained and ordered by increasing *m/z*. Each spectral peak, comprising the fragment *m/z* and normalized intensity, was encoded using Continuous Multi-resolution Peak Encoding (CMPE). The resulting continuous spectral-peak representations were then processed by a 16-layer transformer with a hidden dimension of 1,024, 16 attention heads, a feed-forward dimension of 4,096 and dropout of 0.1.

A learned CLS spectrum token was placed at the beginning of each spectrum sequence. The final hidden state of this token was used as the 1,024-dimensional representation of the complete spectrum, whereas the hidden states aligned with individual spectral peaks were retained as spectral-peak-level embeddings. Thus, the CLS spectrum embedding and the spectral-peak embeddings originated from the same transformer but represented chemical information at different levels.

During self-supervised training, 15% of the valid fragment-ion spectral peaks were sampled without replacement, with at least one spectral peak masked in each spectrum. Both the *m/z* and intensity channels of every selected spectral peak were hidden. The original values were reconstructed using Sequentially Conditioned Fine-grained Reconstruction (SCFR), which decomposed *m/z* prediction into successive resolutions of 10, 1 and 0.1 Da.

Self-supervised training jointly optimized CMPE, the transformer and the reconstruction heads for five epochs using AdamW. The initial and minimum learning rates were 3×10^−5^ and 5×10^−7^, respectively, with a weight decay of 0.01, a warm-up ratio of 0.03 and gradient clipping at 5.0. Training was distributed across six NVIDIA A100 GPUs.

Retention time and ion polarity were subsequently introduced as auxiliary physicochemical objectives while masked spectrum modelling was retained. The retention-time head received the CLS spectrum embedding and was optimized using Huber loss. Before calculating the loss, each retention time in seconds was divided by 600 s. Missing retention-time labels and labels that did not satisfy the predefined self-consistency criterion were excluded from the corresponding minibatch loss. The ion-polarity head also received the CLS spectrum embedding and was optimized using binary cross-entropy.

**Supplementary Figure 14.**
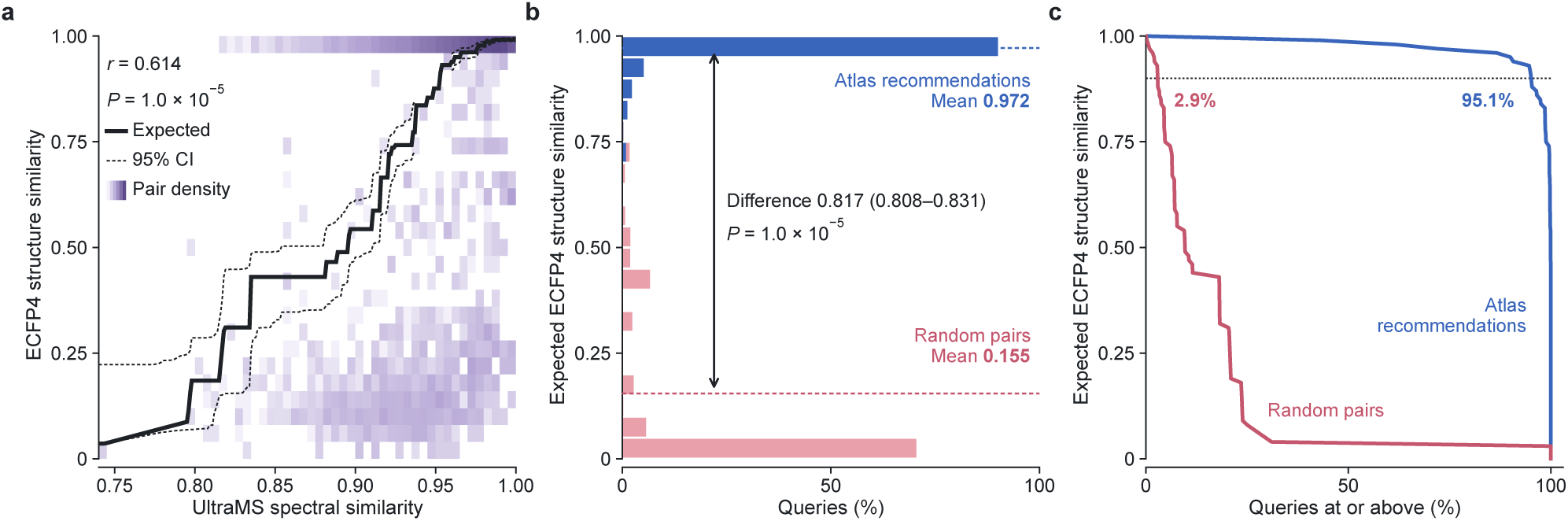
Structural similarity of UltraAtlas recommendations. **a**, Relationship between UltraMS spectral similarity and ECFP4 Tanimoto similarity in reference molecules (22,915 spectra from 2,830 molecules). Shading indicates the density of reference pairs; the solid line indicates the expected structure similarity and dashed lines its 95% confidence interval. *r*, molecule-level Pearson correlation; *P*, permutation test. **b**, Expected structure similarity of Atlas recommendations and of randomly paired spectra matched by ion polarity and precursor mass for 750,464 external spectra; dashed lines indicate the means, and the difference is shown with its 95% confidence interval. **c**, Proportions of Atlas recommendations and mass-matched random pairs remaining above each expected structure-similarity threshold; dotted line, 0.90.

**Supplementary Figure 15.**
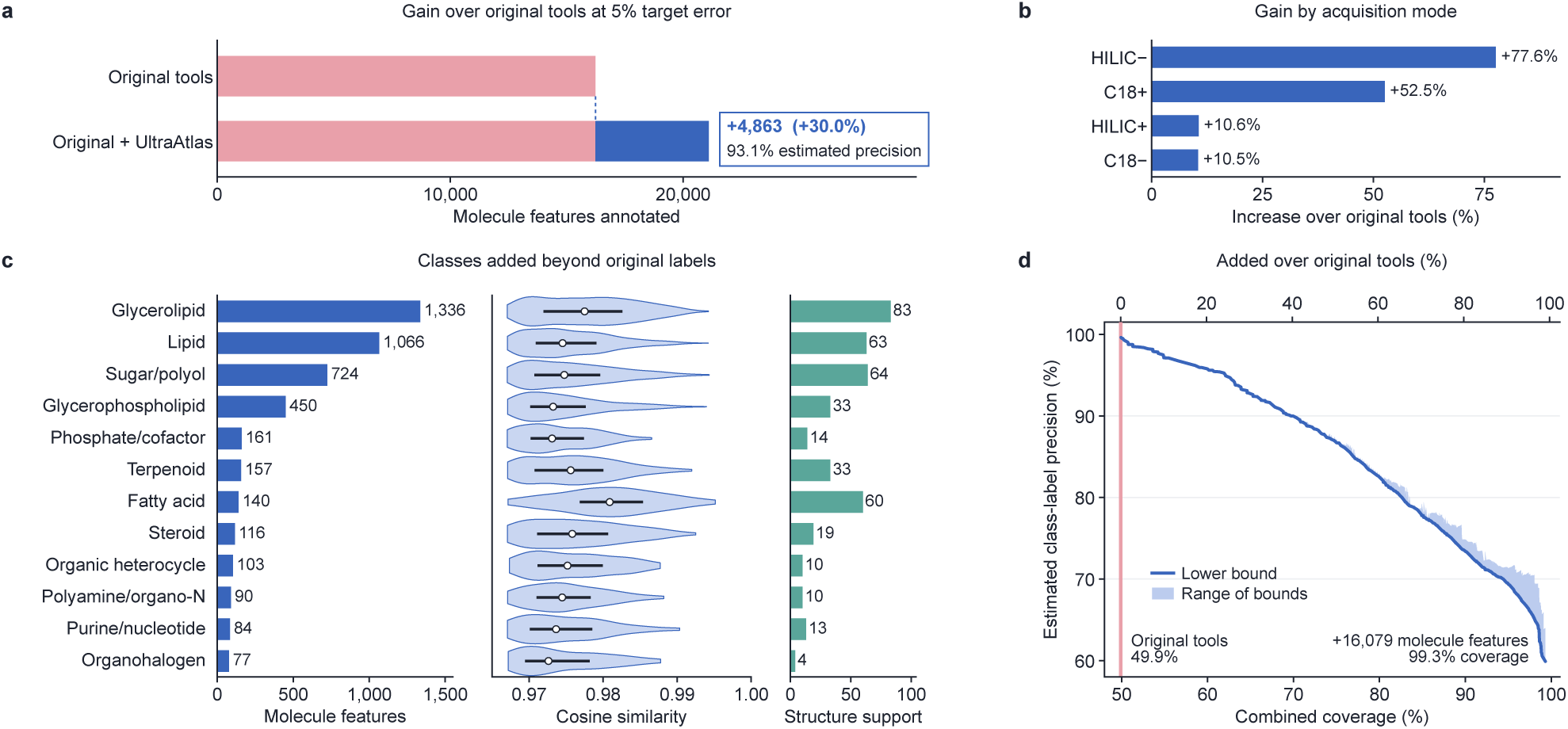
**a**, Molecule features assigned a chemical class by the original GNPS, SIRIUS and CANOPUS analyses, alone and combined with UltraAtlas, among 32,538 molecule features. UltraAtlas adds 4,863 molecule features, a 30.0% increase over the 16,236 originally labelled, with an estimated precision of 93.1% (95% CI 91.4–94.6%). **b**, Increase over the original labels within each acquisition mode. **c**, The twelve most frequent chemical classes added by UltraAtlas among molecule features without an original class label, with the cosine similarity to the Atlas recommendation (white points, medians; bars, interquartile ranges) and the number of these molecule features that also receive a structure-level recommendation. Each molecule feature is counted once. Lipid, organic heterocycle and organosulfur combine a ChEBI class with its directly corresponding RDKit structural class. **d**, Estimated chemical-class precision against combined coverage as the classifier and cosine-similarity thresholds are varied. The top axis expresses coverage as the increase over the original labels, and the vertical line marks the original 49.9%. In **a**–**c**, classifier thresholds are set at an empirical 5% false discovery rate on reference molecules, and the cosine-similarity threshold is calibrated to a 5% target error on reference spectra, on which precision is also estimated. C18, reversed-phase chromatography; HILIC, hydrophilic interaction liquid chromatography; + and *−*, ionization polarity.

Among spectra with usable polarity labels, the auxiliary polarity dataset contained 149,767,344 positive-ion spectra and 10,793,129 negative-ion spectra. To prevent the positive-ion majority from dominating this objective, all 10,793,129 negative-ion spectra were combined with an equal number of positive-ion spectra sampled using random seed 42. Spectra outside this balanced subset remained available for masked spectrum modeling but did not contribute to the polarity loss. The combined objective was

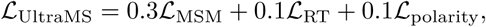

where *L*_MSM_ denotes the masked spectrum modeling loss. Adaptation used AdamW with an initial learning rate of 5×10^−6^, a minimum learning rate of 1×10^−7^, a weight decay of 0.01, a warm-up ratio of 0.05 and gradient clipping at 5.0. The resulting model was used for the principal representation and molecular-identification experiments.

##### Downstream tasks of the self-supervised model

For ion-mode and neutral-loss prediction, the self-supervised UltraMS model was frozen, and a task-specific multilayer perceptron was trained using its CLS spectrum embeddings. Heteroatom prediction additionally incorporated an attention-weighted combination of the spectral-peak embeddings. The model predicted the numbers of S, Cl, F and Br atoms in the precursor molecule, and the learned attention weights were subsequently used to examine whether the most informative spectral peaks represented fragments containing the corresponding element. For the spectral-peak similarity analysis, each annotated query spectral peak was compared with spectral peaks from other spectra according to the cosine similarity between their transformer hidden states. The structural similarity between the query fragment and its closest matches was then evaluated. This analysis used the hidden state of each individual spectral peak directly, without using the CLS spectrum embedding or pooling information from other spectral peaks.

The primary spectrum-to-molecule experiments in the main text used ChemBERTa to represent molecular structures. Canonical molecular SMILES strings were encoded using the fixed ChemBERTa-100M-MLM model, and the final hidden state of its CLS token provided a 768-dimensional molecular embedding. The 1,024-dimensional UltraMS CLS spectrum embedding was mapped into this 768-dimensional ChemBERTa space using a residual multilayer projection. The projection contained a linear skip branch and a nonlinear branch composed of a linear layer, GELU activation, layer normalization, dropout and a second linear layer. The outputs of the two branches were added to produce the final 768-dimensional spectrum embedding. Both the projected UltraMS embedding and the ChemBERTa molecular embedding were *L*_2_-normalized before their cosine similarity was calculated. During training, each minibatch contained matched spectrum–molecule pairs. The projection was trained to increase the similarity between each spectrum and its corresponding molecule while decreasing its similarity to the other molecules in the same minibatch. The UltraMS backbone remained frozen throughout this process. This experiment therefore measured how accurately the pretrained UltraMS CLS spectrum embedding could be matched to molecular structures represented by ChemBERTa.

The supplementary spectrum-to-molecule analyses used molecular fingerprints as a complementary representation of molecular structure. Separate neural-network predictors were trained to infer 2,048-bit, radius-2 Morgan fingerprints from the UltraMS spectrum representations. For each query spectrum, the predicted fingerprint was compared by cosine similarity with the Morgan fingerprints of all molecules in its predefined candidate set, and the candidates were ranked according to this similarity. These experiments tested whether UltraMS representations could recover explicit molecular substructures that distinguished the correct molecule from the other candidates.

For the spectrum-to-spectrum library-search experiment, both query and reference spectra were encoded using the self-supervised UltraMS model. Each CLS spectrum embedding was first layer-normalized and passed through a two-layer projection consisting of a linear layer, GELU activation and a second linear layer. The resulting vector was *L*_2_-normalized to produce a 256-dimensional representation of the spectrum. Training examples consisted of pairs of spectra assigned to the same canonical molecule. For each spectrum in a minibatch, the other spectrum from the same molecule was treated as the correct match, whereas spectra from different molecules served as incorrect matches. The model was trained in both matching directions using the InfoNCE loss with a temperature of 0.07. CMPE and the lower transformer layers remained frozen, whereas the final two transformer layers and the projection were trainable. Training used AdamW for 20 epochs with a batch size of 256, a projection learning rate of 3 × 10^−4^, a transformer learning rate of 3 × 10^−5^, a weight decay of 0.01 and a warm-up ratio of 0.1.

##### Contrastively enhanced UltraMS for broad scale library search and UltraAtlas

The second principal UltraMS model was trained to construct a spectrum space that remained coherent across diverse spectra acquired from the same molecule. Its structure-annotated training resources combined MassSpecGym, MSnLib and Spectraverse. The three processed resources contained 1,255,534 spectra, of which 1,196,493 passed the model-input checks. Following partitioning and replicate-eligibility filtering, 1,125,365 spectra from 73,268 canonical molecular structures, corresponding to 64,507 molecular connectivities, were available for supervised contrastive training.

Training batches contained 64 molecules and up to four spectra per molecule, giving 256 spectrum views per batch. Spectra from the same molecule were treated as positive views, whereas spectra from different molecules provided negative comparisons. The CLS spectrum embedding was passed through a projection head to produce a 512-dimensional vector, which was *L*_2_-normalized before comparison. The model was optimized using a multi-positive supervised contrastive objective with temperature 0.07. The final two transformer layers and the projection head were trainable, whereas the remaining UltraMS parameters were frozen. Training used 20 epochs, an encoder learning rate of 1×10^−5^, a projection-head learning rate of 3×10^−4^, weight decay of 0.01, a warm-up ratio of 0.1, gradient clipping at 1.0 and random seed 42.

For the fixed Fig. 6 retrieval comparison, UltraMS and DreaMS were evaluated using their prespecified final epoch-20 states. UltraAtlas construction and its downstream applications used frozen 512-dimensional spectrum representations from this contrastively enhanced UltraMS training line. We refer to this representation as the UltraAtlas spectrum space. No model parameter was updated during Atlas construction, external-spectrum retrieval, carotid-plaque analysis or scalp-metabolomics analysis.

The chemical-class readout used by UltraAtlas was separate from the contrastively enhanced UltraMS spectrum model. The frozen 512-dimensional representations of replicate spectra from each reference molecule were averaged and normalized to obtain one molecule-level representation. This representation was supplied to a multilayer perceptron containing layer normalization, a 512-to-256 transformation, GELU activation, dropout and the class-specific output layer. An incoming spectrum was first compared with spectra represented in UltraAtlas. Class-level evidence was then transferred from the retained Atlas reference spectrum according to the predefined similarity and ion-mode criteria. The incoming spectrum was not assigned a class by directly applying the class readout to the query itself.

##### MoNA contrastive dataset adaptation

For the UMAP visualization of the negative-ion Spectraverse spectra and the supplementary NPLIB1 experiments, we additionally examined UltraMS representations after molecular-identity adaptation using the public MoNA contrastive dataset released with DreaMS [18]. The dataset contained 25,903 [M + H]^+^ spectra, comprising 22,911 training spectra and 2,992 validation spectra. Valid contrastive examples could be constructed for 19,644 training and 1,817 validation reference spectra. The training and validation partitions shared neither two-dimensional molecular structures, as defined by the first 14 characters of the InChIKey, nor Murcko scaffolds.

For each reference spectrum, another spectrum from the same molecule was used as the positive example. A spectrum from a different molecule was then selected such that its neutral exact mass differed from that of the reference molecule by no more than 0.05 Da. The model was trained to make the reference spectrum more similar to the same-molecule spectrum than to the mass-matched different-molecule spectrum, with a cosine-similarity margin of 0.1. The final four transformer layers and the projection layer were updated for three epochs while keeping the remaining layers frozen. The model checkpoint was selected according to its validation ranking accuracy.

##### Spectral and structural similarity metrics

Raw spectral cosine similarity, learned embedding cosine similarity, and ECFP4 Tanimoto similarity captured distinct levels of information. Specifically, raw spectral cosine measured direct peak overlap between fragment ions, embedding cosine quantified proximity in the learned representation space, and ECFP4 Tanimoto similarity evaluated ground-truth molecular structural similarity.

#### S1.8 Benchmark datasets and evaluation protocol

The benchmark suite was organized according to the capability being evaluated. Representation benchmarks examined masked spectral-peak reconstruction, encoding of ion mode, recognition of neutral losses, prediction of heteroatom content and structural organization of the spectral-peak embeddings. Molecular identification was evaluated using spectrum-to-molecule candidate ranking and spectrum-to-spectrum library search. Unless otherwise stated, UltraMS and all baseline models were evaluated using the same query spectra, molecular candidate sets and reference spectral libraries.

##### Masked spectral-peak reconstruction

Masked spectral-peak reconstruction was evaluated using spectra sampled from GeMS-A10 and the MassSpecGym test partition [27]. We initially sampled 5,000 spectra from each dataset. A spectrum was included in the comparison only when it could be processed by both models and a reconstruction result was successfully obtained from each. After excluding spectra that did not meet these requirements, 4,999 GeMS-A10 spectra and 4,187 MassSpecGym spectra remained for evaluation. Within each spectrum, 15% of the valid fragment-ion spectral peaks were masked without replacement, with at least one spectral peak masked in every spectrum. Reconstruction accuracy was defined as the proportion of masked spectral peaks for which the predicted *m/z* differed from the ground-truth value by no more than 0.05 Da. The mean absolute *m/z* error across the masked spectral peaks was also reported.

##### Evaluation of physicochemical and spectral-peak-level tasks

Ion-polarity information was evaluated using balanced MSnLib partitions [26]. For each ion polarity, 100,000 spectra were used for training, 3,000 for validation and 5,000 for testing, yielding a balanced test set of 10,000 spectra. The UltraMS and DreaMS encoders remained frozen. For UltraMS, the classifier received the 1,024-dimensional CLS spectrum embedding; for DreaMS, it received the corresponding 1,024-dimensional precursor-token embedding. Each representation was passed through the same multilayer perceptron, with hidden dimensions of 512 and 256, to predict the ion polarity. Performance was summarized using overall accuracy, precision, recall and *F*_1_ score. We additionally include two classical classifiers, SVM and Random Forest, as baselines that use *m/z* ratio and intensity as input features to identify which peaks contain the target element and to judge element existence directly from raw spectra. To identify which peaks contain the target element, each peak was scored by its intensity multiplied by the positive SVM coefficient or random-forest feature importance of its *m/z* bin.

Neutral-loss recognition was evaluated using MassSpecGym. Seven predefined neutral losses were considered: H_2_O, NH_3_, CO_2_, HF, HCl, CO and CH_3_. A spectrum received a positive label for a given neutral loss when it contained a fragment-ion spectral peak within 0.02 Da of the precursor *m/z* minus the corresponding neutralloss mass. For each model, its whole-spectrum embedding was concatenated with a learned attention-weighted combination of its spectral-peak embeddings. The resulting 2,048-dimensional representation was passed through a multilabel classifier with a 1,024-dimensional hidden layer and seven output units. Both spectral encoders remained frozen, and only the attention module and classifier were trained. We adopted the public MassSpecGym benchmark with the same train/test split. Approximately 164,000 spectra were used for training, with approximately 16,000 spectra each used for validation and testing. Average precision was calculated for all seven neutral losses, and ROC–AUC was also reported when a complete ROC curve was available.

In pharmaceutical chemistry, approximately 25% of FDA-approved small-molecule drugs contain at least one halogen atom [69], particularly fluorine, which is monoisotopic so that no obvious signal can be gained through isotopic patterns. Additionally, its bond with carbon resists cleavage under typical collision energies, leading to no distinct diagnostic signal that can be used to infer its existence. In addition, many commercially available medicines contain sulfur [70], and many widely used pesticides and herbicides are sulfur-bearing compounds whose residues require routine monitoring in food and environmental samples [71, 72]. To test whether learned representations can resolve these chemically distinct and analytically challenging elements, we evaluated heteroatom prediction targeting the numbers of S, Cl, F and Br atoms in molecules across MassSpecGym spectra. The UltraMS and DreaMS encoders remained frozen, while an element-specific attention module assigned a score to each fragment-ion spectral peak and combined the spectral-peak embeddings for molecular-level prediction. The resulting models were trained for 15 epochs by minimizing the mean squared error between the predicted and true atom counts, and their performance on held-out spectra was summarized using *R*^2^. For each target element and each value of *k*, a prediction was counted as correct when at least one of the *k* highest-scored spectral peaks was assigned to a fragment containing that element. In the overall evaluation, both UltraMS and DreaMS demonstrate substantial advantages over all baselines (UltraMS/DreaMS test *R*^2^: S 0.300/0.223, Cl 0.188/0.123, F 0.118/0.079, Br 0.231/0.200), while classical models operating on raw features yield near-zero or even negative *R*^2^ scores (Fig. 2h).

The structural relationship of the spectral-peak embeddings was evaluated using MAGMa-annotated MSnLib spectra. After filtering spectra and fragment assignments for annotation quality, the dataset contained 176,138 annotated spectral-peak embeddings from 7,539 spectra. We filter out spectra with annotation ratios less than 35%, yielding 60,000 reliably annotated spectrum peaks in total. We use peaks from 20% of these spectra as queries to search their neighbors leveraging the spectrum peak embeddings from UltraMS and DreaMS. To compare with other methods, we use the remaining 80% data to train two additional embedding models using SVM and Random Forest to project spectrum peaks to embeddings by using spectrum peak information including *m/z*, intensity and spectral meta-information (precursor *m/z*, collision energy, number of spectral peaks, adduct type and ion mode). For each query spectral peak, spectral peaks from all other spectra were ranked by the cosine similarity between their embeddings. The highest-ranked matches were compared with the query using four complementary measures: exact agreement of the assigned fragment formula, agreement of the fragment family, ECFP4 Tanimoto similarity between the assigned fragment structures and absolute difference in *m/z*.

##### Negative-ion embedding evaluation

The negative-ion embedding analysis used the released Spectraverse partitions [21]. We retained only negative-ion spectra and excluded molecules that were also present in the MoNA adaptation dataset by comparing the first 14 characters of their InChIKeys. One spectrum was then retained for each remaining molecule, yielding 661 molecule–spectrum pairs.

Molecules were grouped by structural similarity using Butina clustering of RDKit molecular fingerprints. The spectrum embeddings were visualized using UMAP and coloured according to these molecular groups. We evaluated how well the spectrum embeddings separated the molecular groups using normalized mutual information, adjusted Rand index and silhouette score.

##### Spectrum-to-molecule benchmark datasets

Spectrum-to-molecule identification was evaluated primarily on MSnLib [26], with MassSpecGym [27] and NPLIB1 used as supplementary benchmarks. The original MSnLib release contains more than 2.3 million MS*^n^* spectra for 30,008 unique molecular structures [26]. We used the MS^2^ export files of the revised MSnLib Zenodo release (DOI: 10.5281/zenodo.16984129), covering nine compound sub-libraries in positive- and negative-ion modes; in addition to individual scans, these files contain spectra merged across scans of the same or all collision energies and pseudo-MS^2^ spectra summarizing complete MS*^n^* fragmentation trees. We merged all spectra from these files and partitioned them at the level of molecular formula: the 21,107 unique molecular formulas were divided into training, validation and test sets at a ratio of 8:1:1, and all spectra sharing a molecular formula were assigned to the same partition. The resulting MSnLib benchmark contained 560,084 spectra, comprising 446,667 training, 55,980 validation and 57,437 test spectra.

NPLIB1, originally released as the CANOPUS training benchmark, provided three data splits with predefined molecular candidate lists [9, 10]. Each MS2 block was treated as one query and matched against its corresponding candidate list. Within each split, the training, validation and test sets did not share spectra, parent spectra or molecules. After removing queries without a valid candidate list, the three test sets contained 3,699 query blocks in total. Top-1 accuracy was calculated across all 3,699 query blocks, giving equal weight to each block. Molecule-level and parent-spectrum-level results were also reported.

##### Detail of the MSnLib benchmark partition

For Spectrum-to-molecule task, we partitioned MSnLib by molecular formula rather than by individual molecule, which is stricter than a conventional molecule-disjoint split. Molecules with the same formula, such as constitutional isomers and stereoisomers, are the most difficult candidates to distinguish by MS/MS, and they are common in MSnLib: 71.8% of library molecules share their formula with at least one other molecule. Under a random molecule-level split, 68.7% of test molecules would have a same-formula isomer in the training set (mean of ten random splits with partition sizes matched to the formula split). The formula partition instead places all molecules sharing a formula in the same partition, so that no molecular formula (0/21,107) or molecule (0/47,662) is shared between partitions, and no test molecule shares its InChIKey connectivity (first 14 characters) with a training molecule (0/4,889). At the same time, the partition keeps the structural overlap with training at the level of a molecule-level split. The fraction of test molecules sharing a Bemis–Murcko scaffold with the training set is the same as under a random molecule-level split (36.7% in both cases), and 72.8% of test scaffolds do not occur in training. In this benchmark, no molecules sharing a test-set formula are available during training. Accurate identification therefore cannot rely on memorizing familiar structures; instead, it requires extracting structure-specific fragmentation evidence directly from the spectrum itself. Performance on MSnLib thus directly evaluates how well a model has captured the general relationship between chemical structure and MS/MS fragmentation—a capability essential for annotating the uncharacterized molecules that constitute the vast majority of spectra in untargeted metabolomics.

##### MSnLib spectrum-to-spectrum library search

The MSnLib library-search benchmark evaluated protonated and deprotonated spectra separately. The first task contained only [M + H]^+^ spectra, and the second contained only [M − H]^−^ spectra, so that spectral matching was not confounded by differences in precursor ion type. Unlike the formula-level partition used for spectrum-to-molecule identification, this benchmark used a separate molecule-level partition, because the two tasks address different questions. Spectrum-to-molecule identification asks whether a model can recognize a molecule it has never encountered among structurally similar candidates; all molecules sharing a molecular formula were therefore withheld together. Library search instead asks whether a query spectrum can be matched to other spectra of the same molecule in a reference library, which reflects how spectral libraries are used in practice: reference spectra of the target molecule are present, and the challenge is to recognize them among the spectra of many other molecules, most critically those of its same-formula isomers. A molecule-level partition matches this setting. Each test molecule and all of its spectra were withheld from model training, whereas same-formula isomers of test molecules could appear in the training set and remained in the reference library as competing matches. Because successful library search depends on distinguishing the spectra of a molecule from those of its isomers, exposure to same-formula molecules during training is desirable rather than a source of leakage: a model that has learned how structural differences are expressed in fragmentation should be better able to separate these isomers at test time, while the test molecule itself is never seen. Within each task, molecules (canonical isomeric SMILES) were divided into training, validation and test sets in a 70%/15%/15% ratio, comprising 211,745, 44,881 and 45,010 [M + H]^+^ spectra and 98,401, 20,927 and 20,865 [M − H]^−^ spectra, respectively. One spectrum from each test molecule was selected as the query, and all remaining spectra in the same ion-mode task formed the reference library.

For the [M + H]^+^ task, 6,202 query spectra were searched against 295,434 reference spectra, and 6,201 queries had at least one spectrum from the same molecule in the library. For the [M − H]^−^ task, 3,193 query spectra were searched against 137,000 reference spectra, and 3,160 queries had at least one spectrum from the same molecule in the library. Identification recall was calculated using these 6,201 and 3,160 queries, respectively. A Top-1 match was considered correct when the query and the highest-ranked reference spectrum had the same canonical SMILES.

We also evaluated library search across substantially different collision energies. Molecules were included when they had at least one spectrum acquired at 20 eV and at least one spectrum acquired at 40 eV. These molecules were divided into training, validation and test sets in a 70%/15%/15% ratio. The training set was used to train the models, the validation set was used to select the model checkpoint, and performance was reported only on the test set. For each test molecule, a spectrum acquired at ≤ 20 eV was used as the query and spectra acquired at ≥ 40 eV formed the reference library. The test set contained 6,161 molecules for the [M + H]^+^ task and 2,999 molecules for the [M − H]^−^ task.

At a similarity threshold *τ*, a query was accepted when the similarity of its highest-ranked library match was at least *τ* . Empirical false-discovery rate (FDR) and identification recall were calculated as

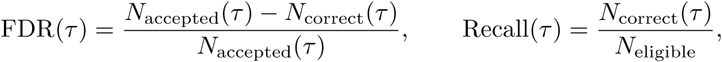

where *N*_accepted_(*τ*) is the number of accepted queries, *N*_correct_(*τ*) is the number whose highest-ranked match came from the same molecule, and *N*_eligible_ is the number of queries with at least one same-molecule spectrum in the library. We reported the recall obtained at 5% empirical FDR and the mean of the highest recall values obtained between 0% and 5% FDR.

To test whether a similarity threshold selected on one set of molecules also worked on unseen molecules, we repeated the analysis using 200 random splits. In each repeat, molecules were divided equally into validation and test sets. We selected the lowest similarity threshold that kept empirical FDR at or below 5% in the validation set and then applied the same threshold to the test set. The resulting test-set FDR and recall measured how well the selected threshold transferred to a different set of molecules.

##### Cross-ionization spectrum-to-spectrum library search

Cross-ionization spectrum-to-spectrum library search used the molecule-level library-search partition described above. Each model was adapted by contrastive training on pairs consisting of one positive-ion and one negative-ion spectrum of the same molecule, using only the 8,265 molecules assigned to the training set in both ion modes; checkpoint selection used the 380 molecules assigned to the validation set in both ion modes. UltraMS spectra were represented by concatenating the CLS embedding with the mean of the spectral peak embeddings, and DreaMS spectra by the precursor-token embedding; each representation was mapped by a projection head to an *L*_2_-normalized 256-dimensional embedding. Both models were trained for 20 epochs with a symmetric InfoNCE loss (temperature 0.07), a batch size of 256 and the last two encoder layers unfrozen, using AdamW (learning rates of 3×10^−4^ for the projection head and 3×10^−5^ for the encoder, weight decay 0.01) with 10% warm-up and cosine decay. The checkpoint with the highest mean bidirectional Recall@1 on the validation molecules was retained. At test time, all 6,202 positive-ion test queries were searched against the complete negative-ion reference library (137,000 spectra from 21,260 molecules), and all 3,193 negative-ion test queries against the complete positive-ion reference library (295,434 spectra from 41,347 molecules). Each reference molecule was scored by the highest cosine similarity between the query and any of its reference spectra, and molecules were ranked by this score. Recall@*k* was calculated over the 2,421 positive-ion and 2,476 negative-ion queries whose correct molecule was present in the opposite-polarity library.

##### Difficulty-stratified benchmark for spectrum-to-spectrum search

The benchmark combined structure-annotated spectra from MassSpecGym, MSnLib and Spectraverse. The three datasets initially contained 1,255,534 processed spectra. After filtering and separating the molecules used for training and testing, 1,125,365 spectra from 73,268 molecules were used for contrastive training. The test set contained 22,915 spectra from 2,830 molecules that were not present in the training set.

We divided the test spectra into challenging and less challenging queries. The challenging set contained 6,159 queries for which the most similar reference spectrum from the same molecule had a raw-spectrum cosine similarity below 0.4. Each query also had a competing reference spectrum from a different molecule with the same ion polarity and a precursor-*m/z* difference of no more than 0.05 Da. Additionally, the raw-spectrum cosine similarity of this competing spectrum differed from that of the correct reference by no more than 0.02. These criteria produced cases in which the correct and competing molecules could not be easily distinguished using precursor mass or direct spectral-peak overlap. Apart from this, the remaining 16,756 test spectra formed the less challenging set.

For the challenging set, we first calculated the AUROC for distinguishing the correct same-molecule reference from the competing different-molecule reference. We then measured the mean ECFP4 Tanimoto similarity of the molecular pairs within the top 10% of spectrum-embedding similarities. Finally, we calculated the Pearson correlation between spectrum-embedding cosine similarity and molecular ECFP4 Tanimoto similarity.

For the less challenging set, we reported the percentage of queries whose highest-ranked spectrum came from the correct molecule. We also calculated the mean ECFP4 Tanimoto similarity between the correct molecule and the molecule represented by the highest-ranked spectrum. Raw-spectrum cosine similarity was included as a training-free comparison in both sets.

UltraMS and DreaMS were trained using the same training spectra and same-molecule spectrum groups. Both models produced 512-dimensional spectrum embeddings, updated their final two transformer layers and were trained for 20 epochs using the same random seed. The epoch-20 model was evaluated for both methods. All comparison methods used the same test spectra and reference spectra.

##### Evaluation on chemical class classification, structural calibration, and large-scale search

Chemical-class recognition used structure-annotated spectra from MassSpecGym, MSnLib and Spectraverse. We defined ten broad structural classes using RDKit and 24 biochemical classes using ChEBI parent terms. A class was included when it contained at least 100 molecules and 500 spectra. All ten RDKit classes and 21 ChEBI classes met these requirements. After dividing the molecules into training and test folds, 20 ChEBI classes had enough molecules for model evaluation.

For each molecule, the 512-dimensional spectrum embeddings from all of its spectra were averaged and normalized. Chemical-class prediction was evaluated using five-fold cross-validation, with all spectra from the same molecule kept in the same fold. Each molecule was therefore predicted by a model trained on different molecules. For each class, we retained the largest group of predictions for which incorrect assignments accounted for no more than 5% of the group. All ten RDKit classes and 19 of the 20 evaluated ChEBI classes produced at least one retained prediction. Class performance was also summarized using coverage, ROC–AUC and average precision.

We next tested whether similarity between UltraMS spectrum embeddings reflected similarity between molecular structures. This analysis used 22,915 spectra from 2,830 molecules that were not used for contrastive training. For each spectrum, we recorded the UltraMS cosine similarity to its highest-ranked reference spectrum and calculated the ECFP4 Tanimoto similarity between their known molecules. We fitted a regression model to predict molecular ECFP4 Tanimoto similarity from UltraMS spectrum cosine similarity. In five-fold evaluation, the regression was fitted on four molecule-disjoint folds and used to predict the fifth. After this evaluation, it was fitted using all five folds to estimate expected ECFP4 Tanimoto similarity for external Atlas matches. Calibration accuracy was measured using mean absolute error and molecule-level Pearson correlation. To test whether the observed correlation between UltraMS spectrum-embedding cosine similarity and molecular ECFP4 Tanimoto similarity exceeded chance, we randomly reassigned the molecular ECFP4 Tanimoto values within groups of the same ion polarity and precursor-*m/z* bin, then recalculated the correlation. We repeated this procedure 100,000 times.

For external evaluation, we sampled one million spectra from GeMS-C across different data sources, ion polarities and precursor-*m/z* ranges. After removing duplicate spectra, 842,193 spectra were searched against UltraAtlas. Query–Atlas pairs with a raw-spectrum cosine similarity above 0.95 were removed, leaving 750,464 pairs for structural analysis. Each Atlas match was compared with a randomly selected background spectrum of the same ion polarity and similar precursor mass. We separately tested how closely the two-stage UltraAtlas search reproduced an exact search of the full spectral library. The same 10,000 external GeMS-C spectra were searched using both methods. We recorded how often the two methods returned the same Top-1 reference spectrum. When their Top-1 results differed, we also checked the rank of the UltraAtlas result in the exact full-library search.

